# Greater male than female variability in regional brain structure across the lifespan

**DOI:** 10.1101/2020.02.17.952010

**Authors:** Lara M Wierenga, Gaelle E Doucet, Danai Dima, Ingrid Agartz, Moji Aghajani, Theophilus N Akudjedu, Anton Albajes-Eizagirre, Dag Alnæs, Kathryn I Alpert, Ole A Andreassen, Alan Anticevic, Philip Asherson, Tobias Banaschewski, Nuria Bargallo, Sarah Baumeister, Ramona Baur-Streubel, Alessandro Bertolino, Aurora Bonvino, Dorret I Boomsma, Stefan Borgwardt, Josiane Bourque, Anouk den Braber, Daniel Brandeis, Alan Breier, Henry Brodaty, Rachel M Brouwer, Jan K Buitelaar, Geraldo F Busatto, Vince D Calhoun, Erick J Canales-Rodríguez, Dara M Cannon, Xavier Caseras, Francisco X Castellanos, Tiffany M Chaim-Avancini, Christopher RK Ching, Vincent P Clark, Patricia J Conrod, Annette Conzelmann, Fabrice Crivello, Christopher G Davey, Erin W Dickie, Stefan Ehrlich, Dennis van ’t Ent, Simon E Fisher, Jean-Paul Fouche, Barbara Franke, Paola Fuentes-Claramonte, Eco JC de Geus, Annabella Di Giorgio, David C Glahn, Ian H Gotlib, Hans J Grabe, Oliver Gruber, Patricia Gruner, Raquel E Gur, Ruben C Gur, Tiril P Gurholt, Lieuwe de Haan, Beathe Haatveit, Ben J Harrison, Catharina A Hartman, Sean N Hatton, Dirk J Heslenfeld, Odile A van den Heuvel, Ian B Hickie, Pieter J Hoekstra, Sarah Hohmann, Avram J Holmes, Martine Hoogman, Norbert Hosten, Fleur M Howells, Hilleke E Hulshoff Pol, Chaim Huyser, Neda Jahanshad, Anthony C James, Jiyang Jiang, Erik G Jönsson, John A Joska, Andrew J Kalnin, Karolinska Schizophrenia Project (KaSP) Consortium, Marieke Klein, Laura Koenders, Knut K Kolskår, Bernd Krämer, Jonna Kuntsi, Jim Lagopoulos, Luisa Lazaro, Irina S Lebedeva, Phil H Lee, Christine Lochner, Marise WJ Machielsen, Sophie Maingault, Nicholas G Martin, Ignacio Martínez-Zalacaín, David Mataix-Cols, Bernard Mazoyer, Brenna C McDonald, Colm McDonald, Andrew M McIntosh, Katie L McMahon, Genevieve McPhilemy, Dennis van der Meer, José M Menchón, Jilly Naaijen, Lars Nyberg, Jaap Oosterlaan, Yannis Paloyelis, Paul Pauli, Giulio Pergola, Edith Pomarol-Clotet, Maria J Portella, Joaquim Radua, Andreas Reif, Geneviève Richard, Joshua L Roffman, Pedro GP Rosa, Matthew D Sacchet, Perminder S Sachdev, Raymond Salvador, Salvador Sarró, Theodore D Satterthwaite, Andrew J Saykin, Mauricio H Serpa, Kang Sim, Andrew Simmons, Jordan W Smoller, Iris E Sommer, Carles Soriano-Mas, Dan J Stein, Lachlan T Strike, Philip R Szeszko, Henk S Temmingh, Sophia I Thomopoulos, Alexander S Tomyshev, Julian N Trollor, Anne Uhlmann, Ilya M Veer, Dick J Veltman, Aristotle Voineskos, Henry Völzke, Henrik Walter, Lei Wang, Yang Wang, Bernd Weber, Wei Wen, John D West, Lars T Westlye, Heather C Whalley, Steven CR Williams, Katharina Wittfeld, Daniel H Wolf, Margaret J Wright, Yuliya N Yoncheva, Marcus V Zanetti, Georg C Ziegler, Greig I de Zubicaray, Paul M Thompson, Eveline A Crone, Sophia Frangou, Christian K Tamnes

**Author notes:** **Corresponding author**: Lara M. Wierenga.

## Abstract

For many traits, males show greater variability than females, with possible implications for understanding sex differences in health and disease. Here, the ENIGMA (Enhancing Neuro Imaging Genetics through Meta-Analysis) Consortium presents the largest-ever mega-analysis of sex differences in variability of brain structure, based on international data spanning nine decades of life. Subcortical volumes, cortical surface area and cortical thickness were assessed in MRI data of 16,683 healthy individuals 1-90 years old (47% females). We observed significant patterns of greater male than female between-subject variance for all subcortical volumetric measures, all cortical surface area measures, and 60% of cortical thickness measures. This pattern was stable across the lifespan for 50% of the subcortical structures, 70% of the regional area measures, and nearly all regions for thickness. Our findings that these sex differences are present in childhood implicate early life genetic or gene-environment interaction mechanisms. The findings highlight the importance of individual differences within the sexes, that may underpin sex-specific vulnerability to disorders.

## Introduction

For a diverse set of human traits and behaviors, males are often reported to show greater variability than females (Hyde 2014). This sex difference has been noted for aspects of personality (Borkenau, McCrae, and Terracciano 2013), cognitive abilities (Arden and Plomin 2006; Johnson, Carothers, and Deary 2008; Roalf et al. 2014), and school achievement (Baye and Monseur 2016). A fundamental question is to what degree these sex differences are related to genetic mechanisms or social factors, or their interactions. Lehre et al. (2009) found compelling evidence for an early genetic or in utero contribution, reporting greater male variability in anthropometric traits (e.g. body weight and height, blood parameters) already detectable at birth. Recent studies suggest greater male variability also in brain structure and its development (Forde et al. 2019; Ritchie et al. 2018; Wierenga et al. 2017; 2019), but studies with larger samples that cover both early childhood and old age are critically needed. Specifically, we do not know when sex differences in variability in brain structure emerge and whether they change with development and throughout life. Yet, data on this could inform us on the origins and factors that influence this phenomenon. For this reason, we set out to analyze magnetic resonance imaging (MRI) data from a large sample of individuals across a very wide age range (n = 16,683, age 1-90) to robustly characterize sex differences in variability of brain structure and test how these differences interact with age.

Many prior studies report sex differences in brain structure, but the specificity, regional pattern and functional relevance of such effects are not clear (Herting et al. 2018; Koolschijn and Crone 2013; Marwha, Halari, and Eliot 2017; Ruigrok et al. 2014; Tan et al. 2016). One reason could be that most studies have examined mean differences between the sexes, while sex differences in variability remain understudied (Del Giudice et al. 2016; Joel et al. 2015). As mean and variance measure two different aspects of the distribution (center and spread), knowledge on variance effects may provide important insights into sex differences in the brain. Recent studies observed greater male variance for subcortical volumes and for cortical surface area to a larger extent than for cortical thickness (Ritchie et al. 2018; Wierenga et al. 2017; 2019). However, further studies are needed to explore regional patterns of variance differences, and, critically, to test how sex differences in variability in the brain unfold across the lifespan.

An important question pertains to the mechanisms involved in sex differences in variability. It is hypothesized that the lack of two parental X-chromosomal copies in human males may directly relate to greater variability and vulnerability to developmental disorders in males compared to females (Arnold 2012). All cells in males express an X-linked variant, while female brain tissues show two variants. In females, one of the X-chromosomes is randomly silenced, as such neighboring cells may have different X related genetic expression (Wu et al. 2014). Consequently, one could expect that in addition to greater variability across the population, interregional anatomical correlations may be stronger in male relative to female brains. This was indeed observed for a number of regional brain volumes in children and adolescents, showing greater within-subject homogeneity across regions in males than females (Wierenga et al. 2017). These results remain to be replicated in larger samples as they may provide clues about mechanisms and risk factors in neurodevelopmental disorders (e.g. attention-deficit/hyperactivity disorder and autism spectrum disorder) that show sex differences in prevalence (Bao and Swaab 2010), age of onset, heritability rates(Costello et al. 2003), or severity of symptoms and course (Goldstein, Seidman, and O’brien 2002).

In the present study, we performed mega-analyses on data from the ENIGMA (Enhancing NeuroImaging Genetics through Meta-Analysis) Lifespan working group (Dima et al., 2020; Frangou et al., 2020; Jahanshad and Thompson 2016). A mega-analysis allows for analyses of data from multiple sites with a single statistical model that fits all data and simultaneously accounting for the effect of site. Successfully pooling lifespan data was recently shown in a study combining 18 datasets to derive age trends of brain structure (Pomponio et al. 2020). This contrasts with meta-analysis where summary statistics are combined and weighted from data that is analyzed at each site (van Erp et al. 2019). MRI data from a large sample (n = 16,683) of participants aged 1 to 90 years was included. We investigated subcortical volumes and regional cortical surface area and thickness. Our first aim was to replicate previous findings of greater male variability in brain structure in a substantially larger sample. Based on prior studies (Forde et al. 2019; Ritchie et al. 2018; Wierenga et al. 2017; 2019) and reports of somewhat greater genetic effect on surface area than thickness (Eyler et al. 2011; Kremen et al. 2013), we hypothesized that greater male variance would be more pronounced for subcortical volumes and cortical surface area than for cortical thickness, and that greater male variance would be observed at both upper and lower ends of the distribution. Our second aim was to test whether observed sex differences in variability of brain structure are stable across the lifespan from birth until 90 years of age, or e.g. increase with the accumulation of experiences (Pfefferbaum, Sullivan, and Carmelli 2004). Third, in line with the single X-chromosome hypothesis, we aimed to replicate whether males show greater interregional anatomical correlations (i.e. within-subject homogeneity) across brain regions that show greater male compared to female variance (Wierenga et al. 2019).

## Methods

### Participants

The datasets analyzed in the present study were from the Lifespan working group within the ENIGMA Consortium (Jahanshad and Thompson 2016). There were 78 independent samples with MRI data, in total including 16,683 (7,966 males) healthy participants aged 1-90 years from diverse ethnic backgrounds (see detailed descriptions at the cohort level in Table 1). Samples were drawn from the general population or were healthy controls in clinical studies. Screening procedures and the eligibility criteria (e.g. head trauma, neurological history) may be found in Supplemental Table 1. Participants in each cohort gave written informed consent at the local sites. Furthermore, at each site local research ethics committees or Institutional Review Boards gave approval for the data collection, and all local institutional review boards permitted the use of extracted measures of the completely anonymized data that were used in the present study.

### Imaging Data Acquisition and Processing

For definition of all brain measures, whole-brain T1-weighted anatomical scan were included. Detailed information on scanner model and image acquisition parameters for each site can be found in Supplemental Table 1. T1 weighted scans were processed at the cohort level, where subcortical segmentation and cortical parcellation were performed by running the T1-weighted images in FreeSurfer using versions 4.1, 5.1, 5.3 or 6.0 (see Supplemental Table 1 for specifications per site). This software suite is well validated and widely used, and documented and freely available online (surfer.nmr.mgh.harvard.edu). The technical details of the automated reconstruction scheme are described elsewhere (Dale, Fischl, and Sereno 1999; Fischl et al. 1999; 2002). The outcome variables included volumes of seven subcortical structures: accumbens, caudate, pallidum, putamen, amygdala, hippocampus, and thalamus (Fischl et al. 2002), and cortical surface area and thickness measures (Dale et al. 1999; Fischl et al. 1999) of 68 regions of the cerebral cortex (Desikan-Killiany atlas) (Desikan et al. 2006). Quality control was also implemented at the cohort level following detailed protocols (http://enigma.ini.usc.edu/protocols/imaging-protocols). The statistical analyses included 13,696 participants for subcortical volumes, 11,338 for surface area measures, and 12,533 participants for cortical thickness analysis.

### Statistical Analysis

Statistical analyses were performed using R Statistical Software. The complete scripts are available in the Supplemental materials in the SI Appendix. In brief, we first adjusted all brain structure variables for cohort, field strength and FreeSurfer version effects. As age ranges differed for each cohort this was done in two steps: initially, a linear model was used to account for cohort effects and non-linear age effects, using a third-degree polynomial function. Next, random forest regression modelling (Breiman 2001) was used to additionally account for field strength and FreeSurfer version. See Supplemental Figure 1 for adjusted values. This was implemented in the R package *randomForest*, which can accommodate models with interactions and non-linear effects.

**Figure 1.**
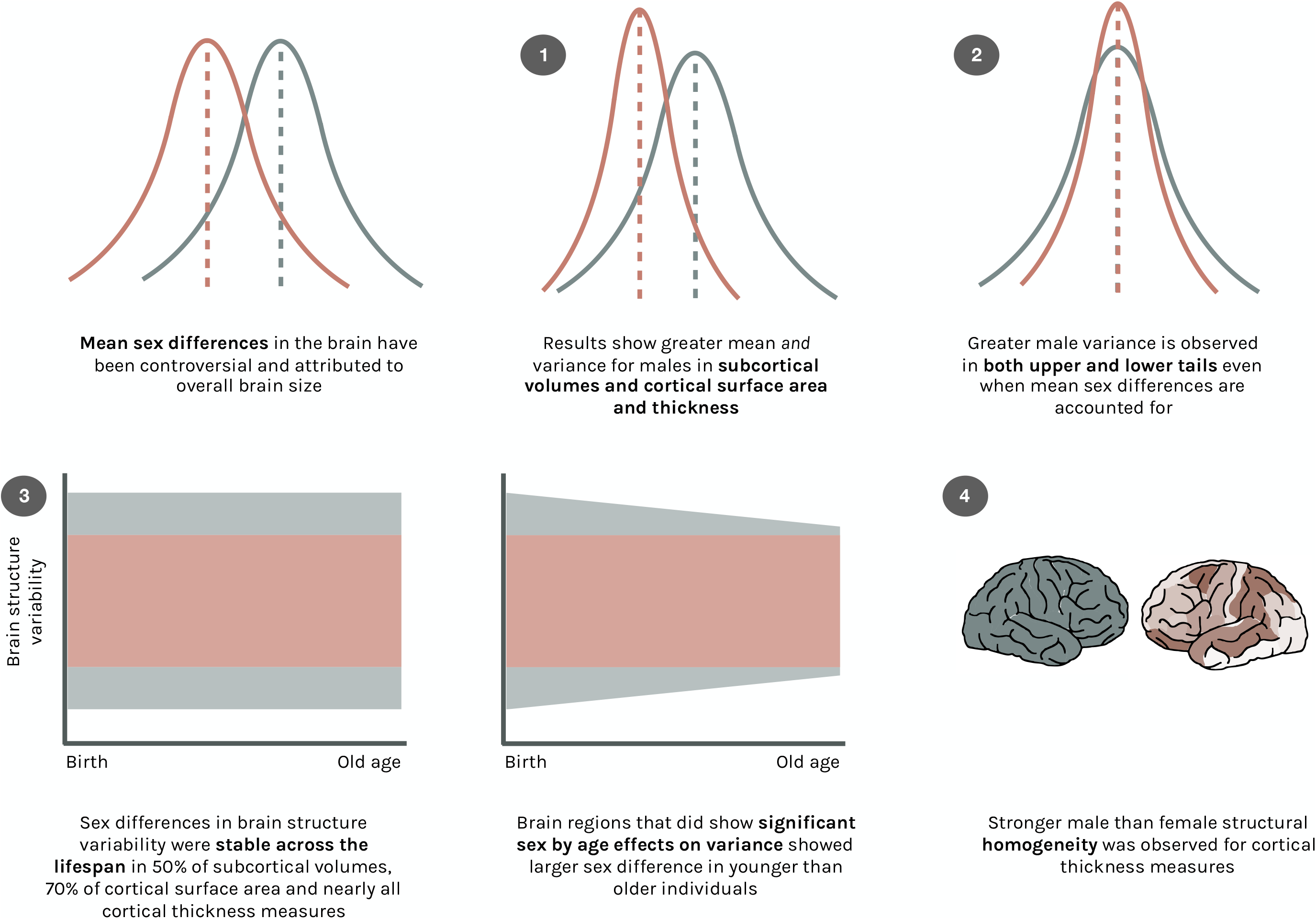
Sex differences in volumetric measures of subcortical volumes (left), cortical surface area (center), and cortical thickness (right). Shown are effect sizes (Cohen’s d-value) of FDR corrected mean sex differences. Greater mean values for males are displayed in blue, greater mean values for females are displayed in red. Darker colors indicate larger effect sizes.

### Mean differences

Mean sex differences in brain structure variables were tested using t-tests (FDR corrected, see(Benjamini and Hochberg 1995)) and effect sizes were estimated using Cohen’s *d*-value. A negative effect size indicates that the mean was higher in females, and a positive effect size indicates it was higher in males. The brain structure variables were adjusted for age and covariates described above. Graphs were created with R package ggseg (Mowinckel and VIdal-Pineiro, 2019).

### Variance ratio

Variance differences between males and females were examined, after accounting for age and other covariates as described above. Fisher’s variance ratio (VR) was estimated by dividing variance measures for males and females. VR was log transformed to account for VR bias (Katzman and Alliger 1992; Lehre et al. 2009). Letting *y*_*i*_ denote the observed outcome for observation number *i* and *y*^_*i*_ its predicted outcome, the residuals were then formed:

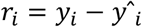

The residual variance *Var* _*males*_ and *Var* _*females*_ were computed separately for males and females, and used to form the test statistic

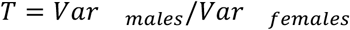

For each outcome, a permutation test of the hypothesis that the sex specific standard deviations were equal, was performed. This was done by random permutation of the sex variable among the residuals. Using β permutations, the *p*-value for the *k*-th outcome measure was computed as

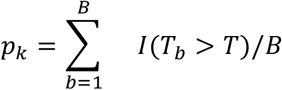

where *I*(*T*_*b*_ ≥ *T*;) is an indicator function that is 1 when *T*_*b*_ ≥ *T*;, and 0 otherwise. Thus, the *p*-value is the proportion of permuted test statistics (*T*_*b*_) that were greater than the observed value *T*; of the test statistic above. Here *B* was set to 10,000. FDR corrected values are reported as significant.

### Shift Function

To assess the nature of the variability difference between males and females, shift functions were estimated for each brain measure that showed significant variance differences between males and females using quantile regression forests (Meinshausen 2006; Rousselet, Pernet, and Wilcox 2017), implemented in the R package quantregForest (see Wierenga et al. 2017) for a similar approach). First, as described above, brain measures were accounted for site, age, field strength and FreeSurfer version. Next, quantile distribution functions were estimated for males and females separately after aligning the distribution means. Let *q* be a probability between 0 and 1. The quantile function specifies the values at which the volume of a brain measure will be at or below any given *q*. The quantile function for males is given as *Q*(*q*|*males*) and for females as *Q*(*q*|*females*). The quantile distance function is then defined as:

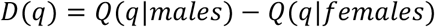

A bootstrap method was used to estimate the standard error of the quantile difference functions, which was used to form approximate 95% confidence intervals. If the quantile distance function is a straight-line parallel to the *x* axis, this indicates a stable difference between the sexes across the distribution and thus no detectable difference in variability. A positive slope indicates greater male variance. More specifically, this would indicate that the males with the largest values have relatively larger values than females with the largest values, and males with the smallest values are relatively smaller values than the females with the smallest values. A negative slope of the quantile distance function would indicate larger variability in females at both ends of the distribution.

### Variance change with age

To study whether the sex differences in variance are stable across the age range we used the residuals of the predicted outcome measure and each individual *i*:

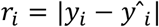

The absolute value of *r*_*i*_ was then used in a regression model. It was next explored whether there was a significant (FDR corrected) age by sex interaction effect using a linear model 1 and quadratic model 2:

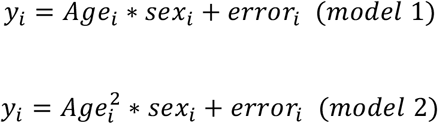

### Anatomical correlation analysis

Inter-regional anatomical associations were assessed by defining the correlation between two brain structures, after accounting for age and other covariates as described above. Anatomical correlation matrices were estimated as previously applied in several structural MRI studies for males and females separately (see e.g. Baaré et al. 2001; Lerch et al. 2006). Next, the anatomical correlation matrix for females was subtracted from the anatomical correlation matrix for males, yielding a difference matrix.

Thus, the Pearson correlation coefficient between any two regions *i* and *j* was assessed for males and females separately. This produced two group correlation matrices *M*_*ij*_ and *F*_*ij*_ where *i,j*, = 1,2,....,*N*, where *N* is the number of brain regions.

Sex specific means and standard deviations were removed by performing sex specific standardization. The significance of the differences between ***M***_***ij***_ and ***F***_***ij***_ was assessed by the difference in their Fisher’s ***z***-transformed values, and ***p***-values were computed using permutations. Whether these significantly differed between the sexes was tested using a Chi-square test.

## Results

### Sex Differences in Mean and Variance

All brain measures were adjusted for cohort, field strength, FreeSurfer version and (non-linear) age. As a background analysis, we first assessed whether brain structural measures showed mean differences between males and females to align our findings to previous reports (Figure 1, Table 2A-C). All subcortical volumes were significantly larger in males, with effect sizes (Cohen’s *d*-values) ranging from 0.41 (left accumbens) to 0.92 (right thalamus), and an average effect size of 0.7. In follow-up analyses with total brain volume as an additional covariate we found a similar pattern, although effect sizes were smaller (Supplemental Table S2A). Also for cortical surface area, all regions showed significantly larger values in males than females, with effect sizes ranging from 0.42 (left caudal anterior cingulate area) to 0.97 (left superior temporal area), on average 0.71.

When total surface area was included as an additional covariate, a similar pattern was observed, although effect sizes were smaller (Supplemental Table S2B). Cortical thickness showed significant mean sex differences in 43 (out of 68) regions, of which 38 regions showed larger thickness values in females than males. These were mostly frontal and parietal regions. The largest effect size, however, was only 0.12 (right caudal anterior cingulate cortex). When total average cortical thickness was included as an additional covariate, nine regions showed a male advantage that was not observed in the raw data analysis, and six of the 38 regions showing female advantage did not reach significance (Supplemental Table S2C).

We then tested for sex differences in variance of brain structure, adjusted for cohort, field strength, FreeSurfer version and (non-linear) age (Figure 2, Tables 2A-C). All subcortical volumes had significantly greater variance in males than females. Log transformed variance ratios ranged from 0.12 (right accumbens) to 0.36 (right pallidum), indicating greater variance in males than females. Similar results were also observed when total brain volume was taken into account (Supplemental Table S2A). Cortical surface area also showed significantly greater variance in males for all regions: variance ratios ranged from 0.13 (left caudal anterior cingulate cortex) to 0.36 (right parahippocampal cortex). This pattern was also observed when total surface area was included in the model (Supplemental Table S2B). Cortical thickness showed significantly greater male variance in 41 out of 68 regions, with the greatest variance ratio being 0.11 (left precentral cortex). Notably, 37 of these 41 regions did not show significantly larger mean thickness values in males. When additionally accounting for total average thickness, we found greater male variance in 39 regions and greater females variance in 5 regions. Also here, significant variance ratios were present in the absence of mean sex differences (Supplemental Table S2C).

**Figure 2.**
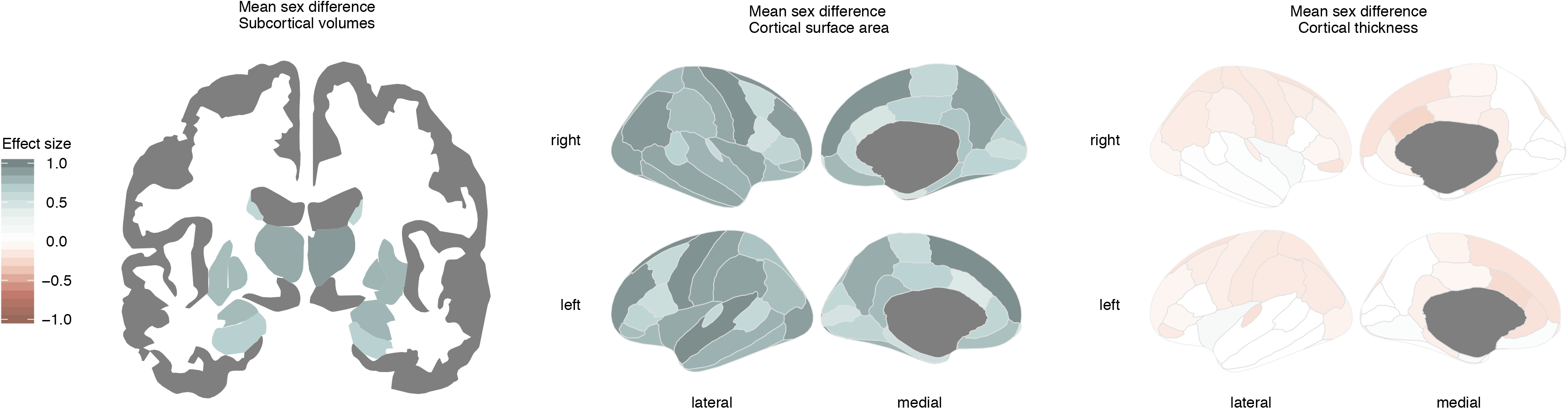
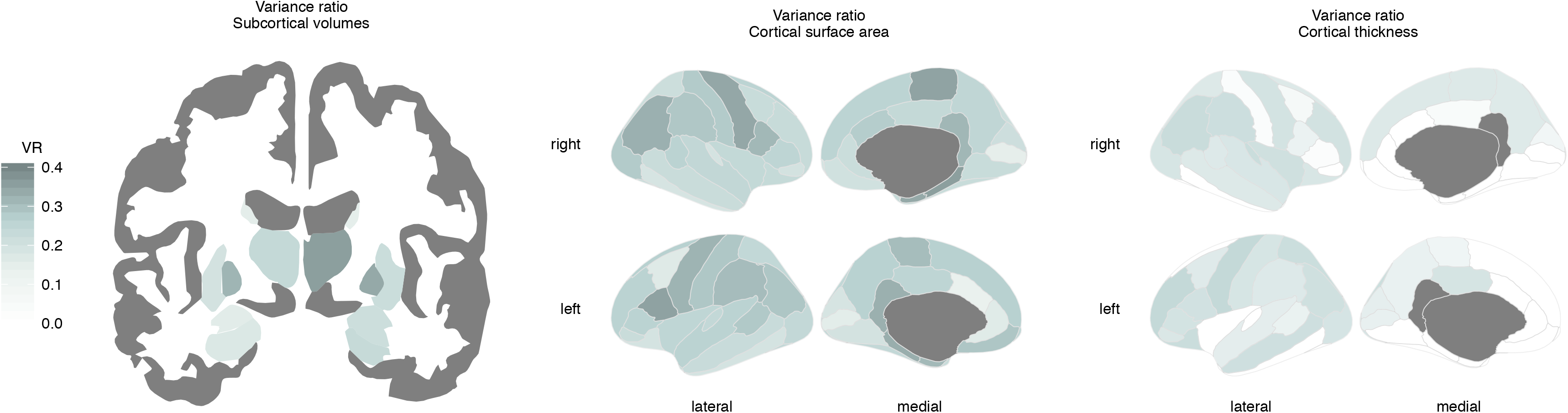
Sex differences in variance ratio for subcortical volumes (Left), cortical surface area (center), and cortical thickness (right). Shown are log transformed variance ratios, where significant larger variance ratio for males than females is displayed in blue ranging from 0 to 1. Darker colors indicate a larger variance ratio.

Next, we directly tested whether the regions showing larger variance effects were also those showing larger mean differences, by correlating the variance ratios with the vector of *d*-values (Supplemental Figure 2). There was a significant association for subcortical volumes (r(12) = 0.7, *P*-value = 0.005), but no significant relation for regional cortical surface area (r(66) = 0.18, *P*-value = 0.14), or thickness (r(66) = −0.21, *P*-value = 0.09).

### Greater Variance in Males at Upper and Lower Tails

In order to characterise how the distributions of males and females differ, quantiles were compared using a shift function(Rousselet et al. 2017). As in the previous models, brain measures were adjusted for cohort, field strength, FreeSurfer version and age. In addition, the distribution means were aligned. Results showed greater male variance at both upper and lower tails for regions that showed significant variance differences between males and females. The top three variance ratio effects for subcortical volume, cortical surface area and cortical thickness are shown in Figure 3.

**Figure 3.**
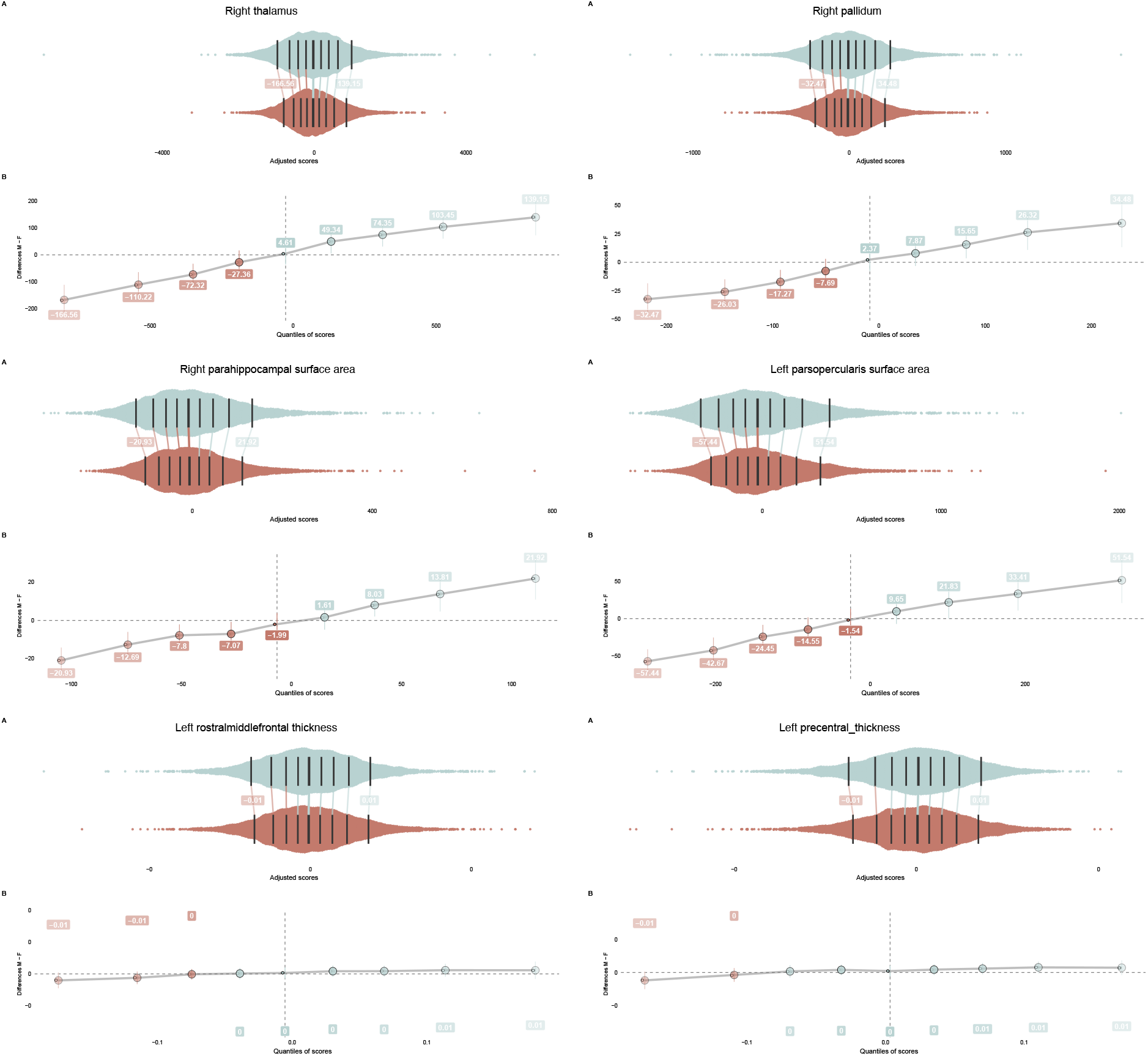
Jittered marginal distribution scatterplots (A) are displayed together with their shift function (B) for the top three variance ratio effects of subcortical volumes (top), cortical surface area (middle) and cortical thickness (right). The central, darkest line on each distribution is the median, note that main sex effects are removed. The other lines mark the deciles of each distribution. The shift values are included, which refer to the number of units that the male (upper) distribution would have to be shifted to match the female (lower) distribution. Confidence intervals are included for each of these shift values.

### Variance Difference Between Sexes Across Age

We next tested whether the sex differences in variance interacted with age (Figure 4 and supplemental Figure 3). In this set of analyses, brain measures were adjusted for cohort, field strength, and FreeSurfer version. For 50% of the subcortical volume measures there was a significant interaction, specifically for the bilateral thalami, bilateral putamen, bilateral pallidum and the left hippocampus (Table 3A, Figure 5). Cortical surface area showed significant interaction effects in 30% of the cortical regions (Table 3C, Figure 5). In both cases, younger individuals tended to show greater sex differences in variance than older individuals. For cortical thickness, an interaction with age was detected only in the left insula (Table 3B, Figure 5). This region showed greater male than female variance in the younger age group, whereas greater female variance was observed in older individuals.

**Figure 4.**
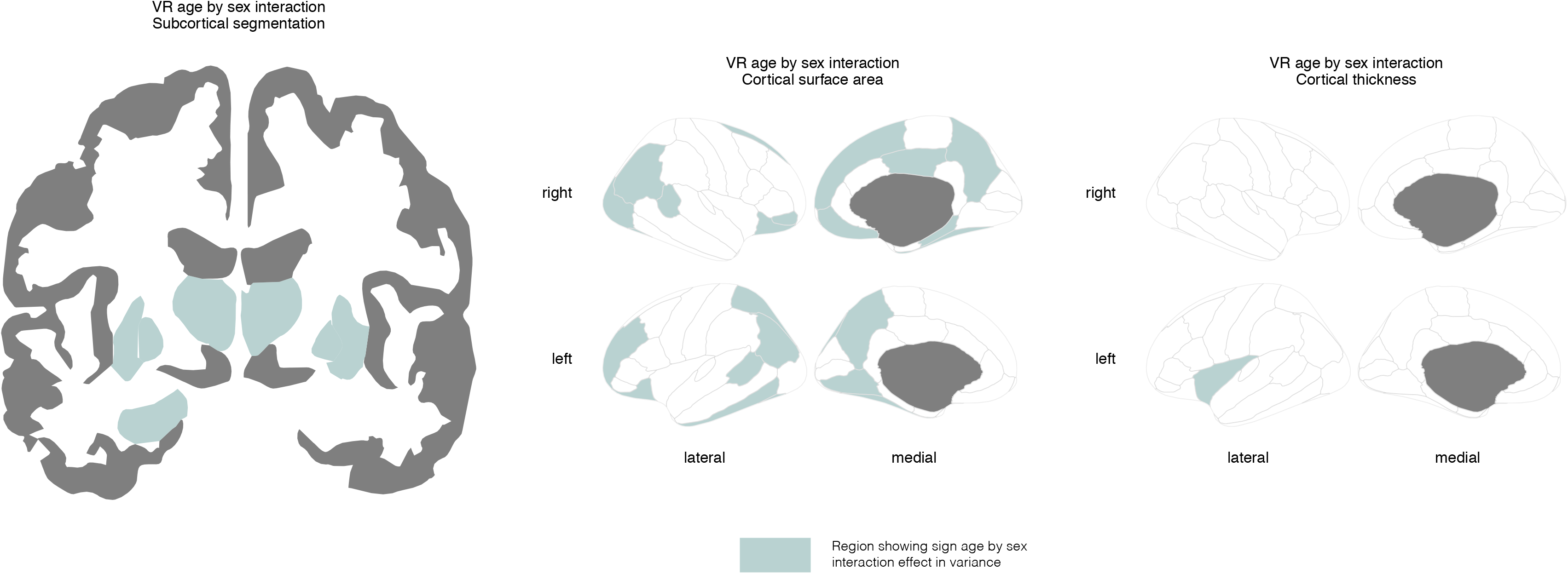
Regions where sex differences in variability of brain structure interacted with age displayed for subcortical volumes (left), cortical surface area (center), and cortical thickness (right).

**Figure 5.**
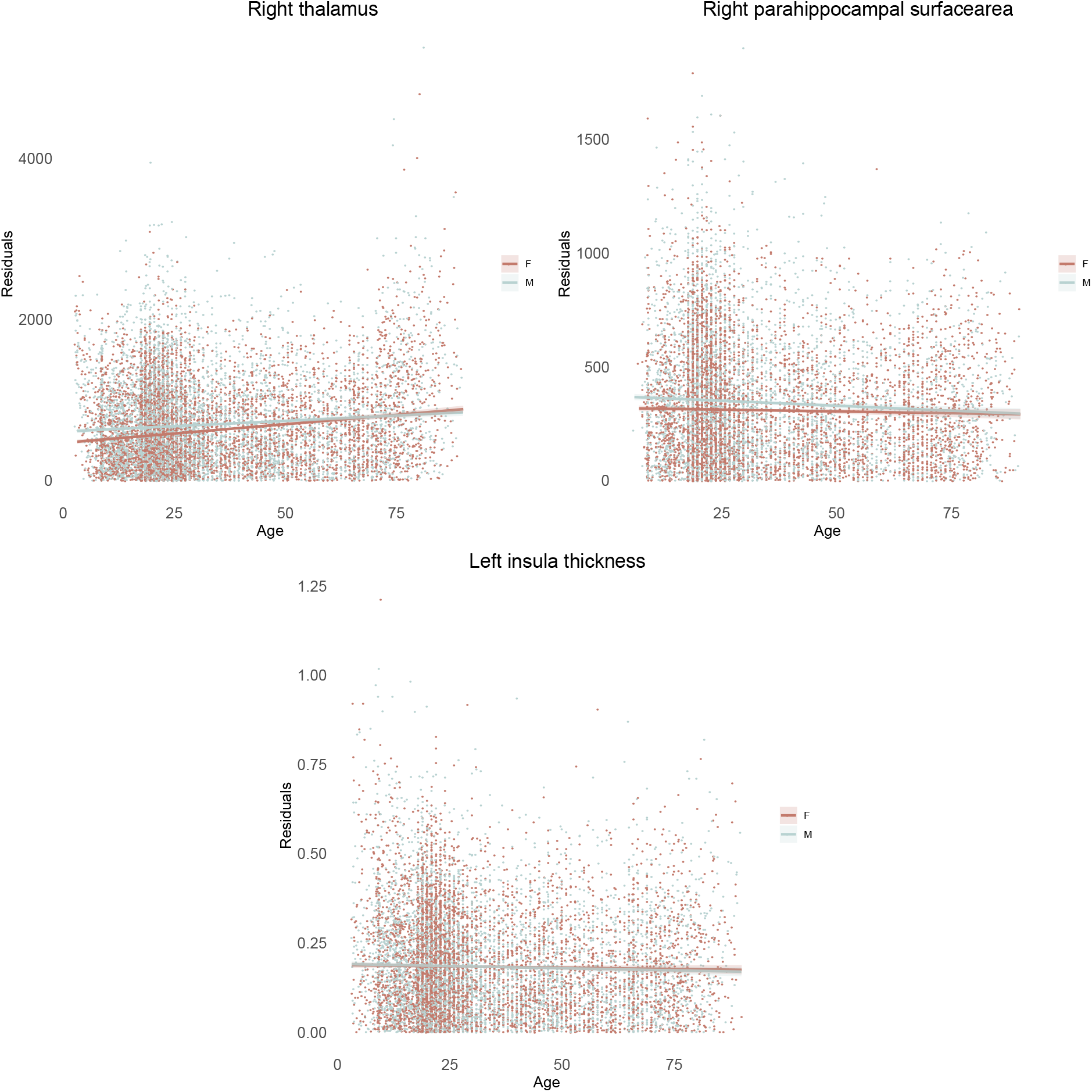
Sex differences in variability interacted with age in 50% of the subcortical volumes, 30% of the surface area measures, and only one thickness measure. Three representative results are shown: right thalamus volume (top left), surface area of the right parahippocampal gyrus (top right) and thickness of the left insula (bottom center). Absolute residual values are modeled across the age range. Effects showed larger male than female variance in the younger age group, this effect attenuated with increasing age.

Next, these analyses were repeated using a quadratic age model (Supplemental Tables 3A-C). None of the subcortical or cortical surface area measures showed quadratic age by sex interaction effects in variance. Cortical thickness showed significant quadratic age by sex effects in two regions; left superior frontal cortex and right lateral orbitofrontal cortex.

### Sex Differences in Anatomical Correlations

Finally, we tested whether females showed greater diversity than males in anatomical correlations by comparing inter-regional anatomical associations between males and females. Using permutation testing (B = 10000), the significance of correlation differences between males and females was assessed.

Of the 91 subcortical-subcortical correlation coefficients, 2% showed significantly stronger correlations in males, while, unexpectedly, 19% showed stronger correlations in females (tested two-sided) (Figure 6A). A chi-square test of independence showed that this significantly differed between males and females, *X*^2^(1, *N* = 18) = 10.889, *p* < .001. For surface area, no significant difference between males and females were observed: significantly stronger male homogeneity was observed in 4% of the 2,278 unique anatomical correlations, and similarly females also showed significantly stronger correlations in 4% of the anatomical associations (Figure 6B). For thickness, stronger male than female homogeneity was observed in 21% of the correlations, while stronger female correlations were observed in <1% of the correlations (Figure 6C). This difference was significant, *X*^2^(1, *N* = 484) = 460.300, *p* < .001.

**Figure 6 A-C.**
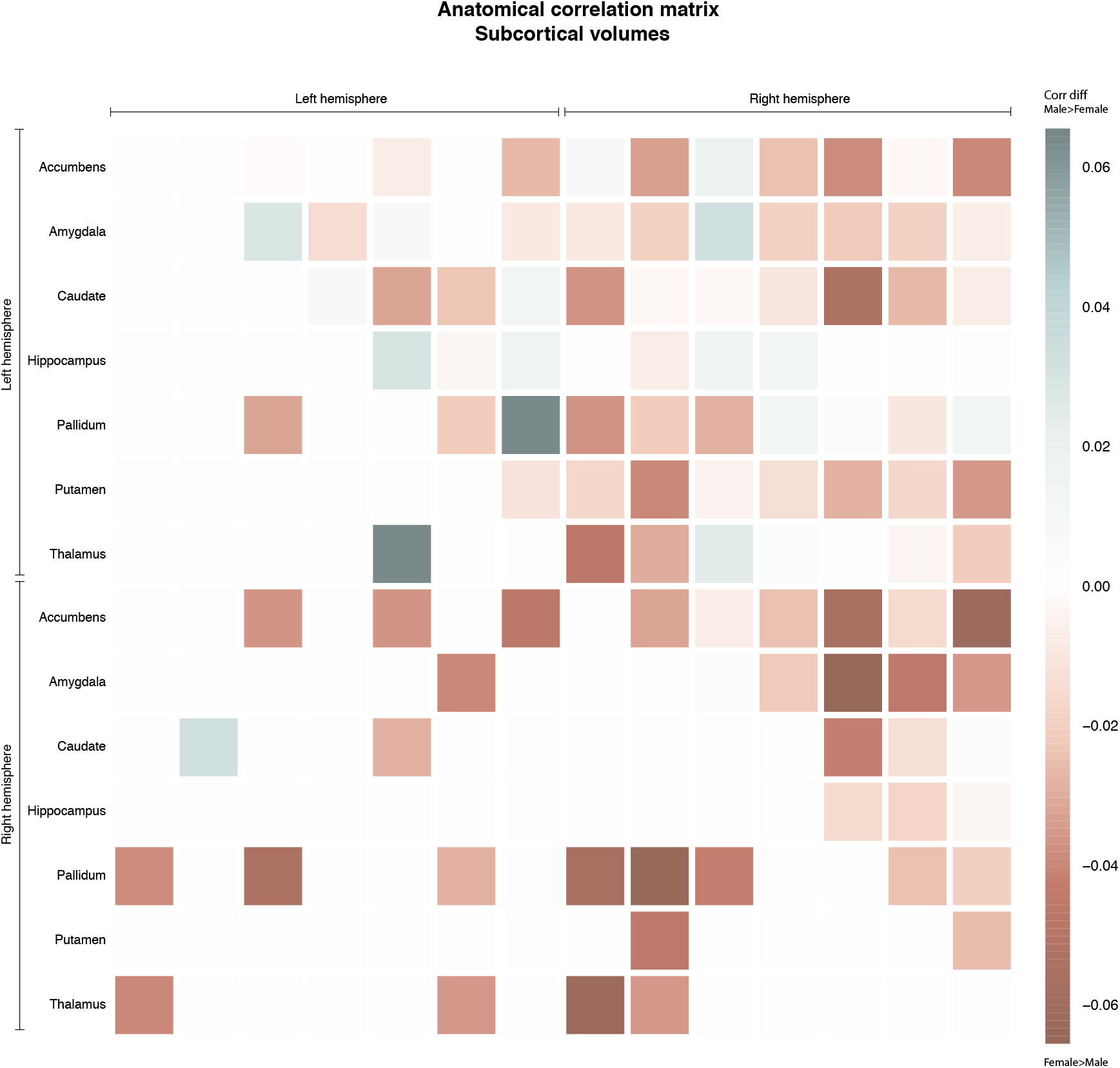
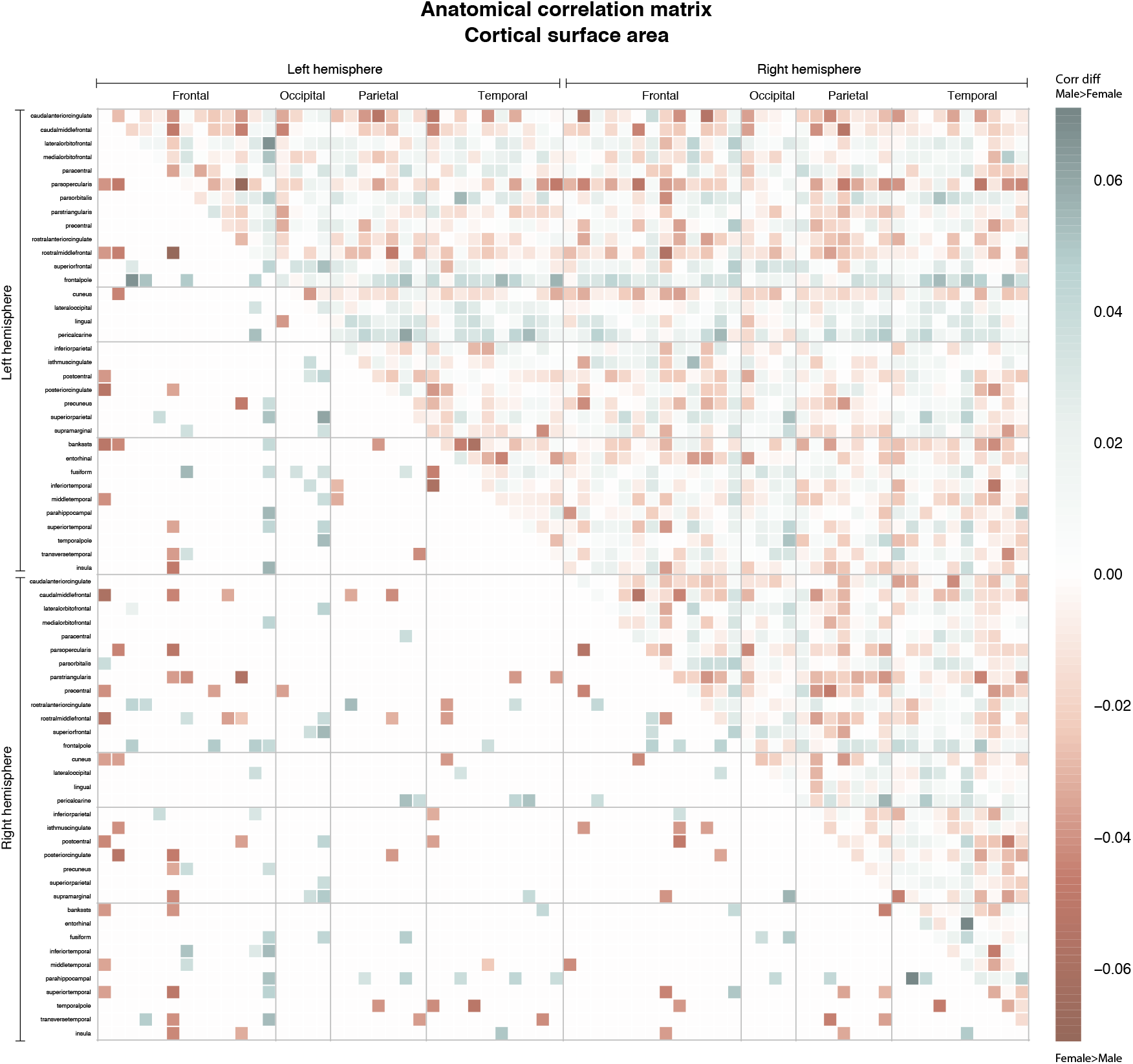
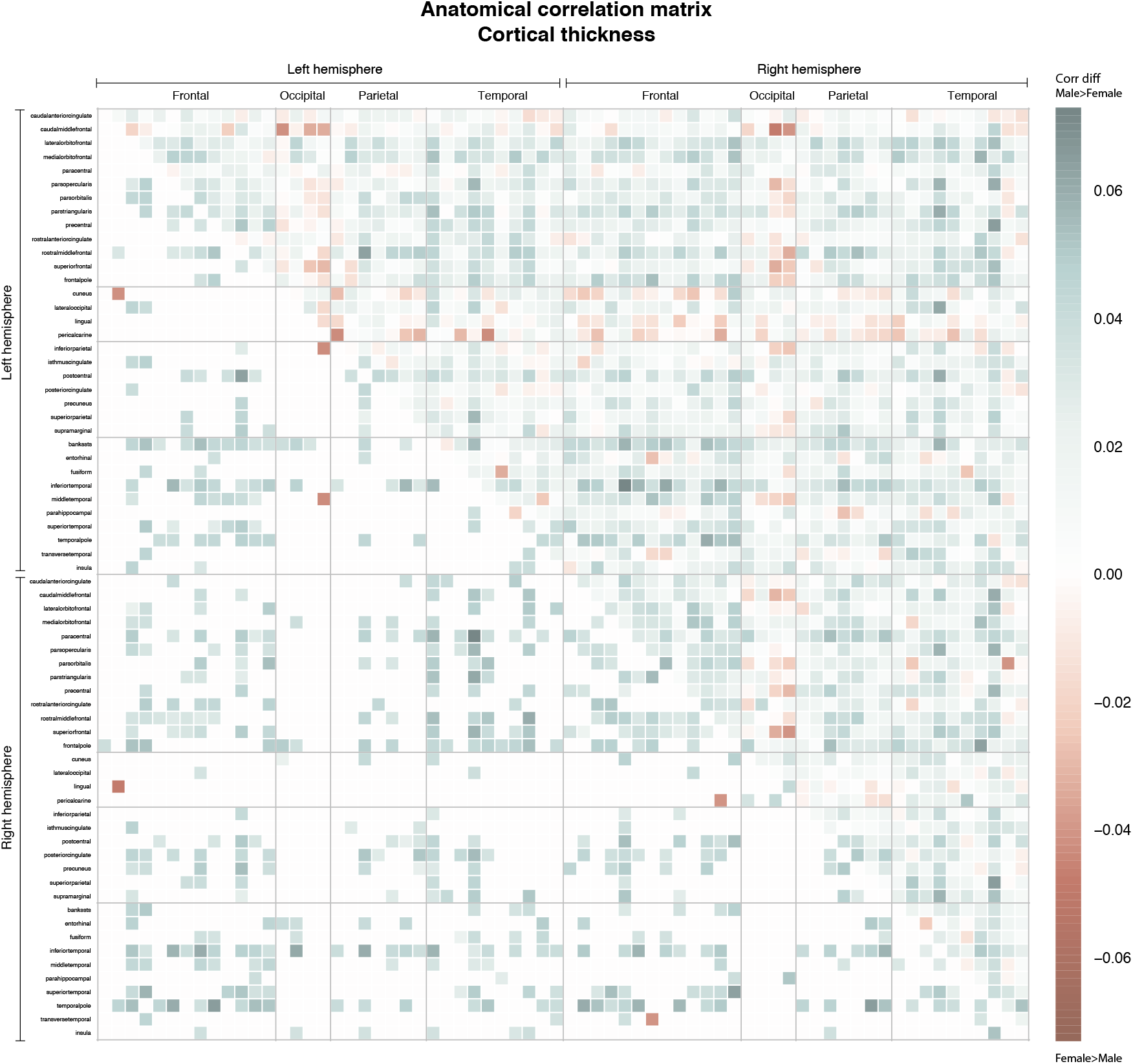
Stronger anatomical correlations for males than females are indicated in blue (larger homogeneity in males than females), while stronger correlations for females are displayed in red (larger homogeneity in females than males). The bottom left half shows the significant variance ratio’s only, using two sided permutation testing. Results are displayed for subcortical volumes (A), surface area (B) and cortical thickness (C). Cortical regions are ordered by lobe and hemisphere (left frontal, left occipital, left parietal, left temporal, right frontal, right occipital, right parietal, right temporal).

## Discussion

In this study, we analyzed a large lifespan sample of neuroimaging data from 16,683 participants spanning nine decades of life starting at birth. Results confirmed the hypothesis of greater male variability in brain structure (Forde et al. 2019; Ritchie et al. 2018; Wierenga et al. 2017; 2019). Variance differences were more pronounced for subcortical volumes and regional cortical surface area than for regional cortical thickness. We also corroborated prior findings of greater male brain structural variance at both upper and lower tails of brain measures (Wierenga et al. 2017). These variance effects seem to describe a unique aspect of sex differences in the brain that does not follow the regional pattern of mean sex differences. A novel finding was that sex differences in variance appear stable across the lifespan for around 50% of subcortical volumes, 70% of cortical surface area measures and almost all cortical thickness measures. Unexpectedly, regions with significant change in variance effects across the age range showed decreasing variance differences between the sexes with increasing age. Finally, we observed greater male inter-regional homogeneity for cortical thickness, but not for surface area or subcortical volumes, partly replicating prior results of greater within-subject homogeneity in the male brain (Wierenga et al. 2017). Unexpectedly, subcortical regions showed stronger interregional correlation in females than in males.

Greater male variance was most pronounced in brain regions involved in planning, regulation and inhibition of motor movements (pallidum, right inferior parietal cortex and paracentral region), episodic memory (hippocampus), and multimodal sensory integration (thalamus) (Aron, Robbins, and Poldrack 2004; Burgess, Maguire, and O’Keefe 2002; Grillner et al. 2005). In addition, the early presence of sex differences in brain structural variability may be indicative of genetic effects, in line with findings in a pediatric sample (Wierenga et al. 2017). We also observed that sex differences in structural variation are either stable or may reduce in old age. Longitudinal designs are, however, needed to address the mechanisms underlying this observation.

The expression of greater male variability in both upper and lower tails of the distribution may be related to architectural and geometric constraints that are critical for a delicate balance for effective local-global communication. For example, neurons only partly regulate their size, and the number of neural connections does not vary strongly with neocortical size across species (Stevens 1989). Although axon size and myelin can compensate firing rates in larger brains by speeding up conduction time, there is a limited energy budget to optimize both volume and conduction time (Buzsáki, Logothetis, and Singer 2013). As such, extreme brain structure (in both directions) may come at a cost. This is in line with recent findings that show that extreme neural activity patterns may induce suboptimal expressions of mental states (Northoff and Tumati 2019). Interestingly, it has been found that individuals with autism spectrum disorder show atypical patterns of brain structure and development in both the upper and lower range (Zabihi et al. 2019), suggesting a possible link between greater male variability and vulnerability for developmental disorders (see also Alnæs et al. 2019)). Together with our findings, this opens up new approaches to understanding sex biased developmental disorders, beyond group-level mean differences.

Although most results showed stable sex differences with increasing age, half of the subcortical regions and a quarter of the cortical surface area measures showed decreasing sex differences in variance. What stands out is that in all these regions, sex differences in variance were largest in young compared to older age. This is indicative of early mechanisms being involved. Furthermore, for subcortical regions, the patterns showed larger volumetric increases in females then in males. For surface area, interaction effects showed mostly stable variance across age in females, but decreases in variability in males. The observation that there were no significant quadratic interactions makes it unlikely that pubertal hormones may affect greater male variance. Yet, the decrease in male variance in older age, may be indicative of environmental effects later in life. Alternative explanation may be the larger number of clinical or even death rates in males that may lead to some sex difference in survival (Chen et al. 2008; Ryan et al. 1997).

Factors underlying or influencing sex differences in the brain may include sex chromosomes, sex steroids (both perinatal or pubertal), and the neural embedding of social influences during the life span (Dawson, Ashman, and Carver 2000). Although we could not directly test these mechanisms, our findings of greater male variance, that are mostly stable across age, together with the greater male inter-regional homogeneity for cortical thickness are most in line with the single X-chromosome expression in males compared to the mosaic pattern of X-inactivation in females (Arnold 2012). Whereas female brain tissue shows two variants of X-linked genes, males only show one. This mechanism may lead to increased male vulnerability, as is also seen for a number of rare X-linked genetic mutations (Chen et al. 2008; Craig, Haworth, and Plomin 2009; Johnson, Carothers, and Deary 2009; Reinhold and Engqvist 2013; Ryan et al. 1997). <u>None of the other sex effects mentioned above predict these specific inter and intra-individual sex differences in brain patterns</u>. Future studies are, however, needed to directly test these different mechanisms. Furthermore, the observation that greater male homogeneity was only observed in cortical thickness, but not cortical surface area or subcortical volumes, may speculatively indicate that X-chromosome related genetic mechanisms may have the largest effect on cortical thickness measures.

This paper has several strengths including its sample size, the age range spanning nine decades, the inclusion of different structural measures (subcortical volumes and cortical surface area and thickness) and the investigation of variance effects. These points are important, as most observed mean sex differences in the brain are modest in size (Joel and Fausto-Sterling 2016). We were able to analyze data from a far larger sample than those included in recent meta-analyses of mean sex differences (Marwha et al. 2017; Ruigrok et al. 2014; Tan et al. 2016), and a very wide age range covering childhood, adolescence, adulthood and senescence. The results of this study may have important implications for studies on mean sex differences in brain structure, as analyses in such studies typically assume that group variances are equal, which the present study shows might not be tenable. This can be particularly problematic for studies with small sample sizes (Rousselet et al. 2017).

The current study has some limitations. First, the multi-site sample was heterogeneous and specific samples were recruited in different ways, not always representative of the entire population. Furthermore, although structural measures may be quite stable across different scanners, the large number of sites may increase the variance in observed MRI measures, but this would be unlikely to be systematically biased with respect to age or sex. In addition, variance effects may change in non-linear ways across the age-range. This may be particularly apparent for surface area and subcortical volume measures, as these showed pronounced non-linear developmental patterns through childhood and adolescence (Tamnes et al. 2017; Wierenga et al. 2018). Also, the imbalanced number of subjects across the age range may have diminished variability effects in the older part of the age range. The present study has a cross-sectional design. Future studies including longitudinal data are warranted to further explore the lifespan dynamics of sex differences in variability in the brain. Last, one caveat may be the effect of movement on data quality and morphometric measures. As males have been shown to move more than females in the scanner (Pardoe, Kucharsky Hiess, and Kuzniecky 2016), this may have resulted in slight under estimations of brain volume and thickness measures for males (Reuter et al. 2015). Although quality control was conducted at each site using the standardized ENIGMA cortical and subcortical quality control protocols (http://enigma.ini.usc.edu/protocols/imaging-protocols/), which involve a combination of statistical outlier detection and visual quality checks and a similar number of males and females had partially missing data (52.4% males), we cannot exclude the possibility that in-scanner subject movement may have affected the results. Nevertheless, we do not think this can explain our finding of greater male variance in brain morphometry measures, as this was seen at both the upper and lower ends of the distributions.

## Conclusions

The present study included a large lifespan sample and robustly confirmed previous findings of greater male variance in brain structure in humans. We found greater male variance in all brain measures, including subcortical volumes and regional cortical surface area and thickness, at both the upper and the lower end of the distributions. The results have important implications for the interpretation of studies on (mean) sex differences in brain structure. Furthermore, the results of decreasing sex differences in variance across age opens a new direction for research focusing on lifespan changes in variability within sexes. Our findings of sex differences in regional brain structure being present already in childhood may suggest early genetic or gene-environment interaction mechanisms. Further insights into the ontogeny and causes of variability differences in the brain may provide clues for understanding male biased neurodevelopmental disorders.

---

*Supplemental Figure 1.* Boxplot visualization of comparison of right hippocampal volume, and parahippocampal surface area and thickness before and after adjustment. As age ranges differed for each cohort adjustments were performed in two steps: initially, a linear model was used to account for cohort and non-linear age effects. Next, random forest regression modelling was used to additionally account for field strength and FreeSurfer version. In the left panel, volumes were not adjusted, this displays the raw data for each cohort. In the right panel, volumes were adjusted.

*Supplemental Figure 2.* Correlation between variance ratio and vector of d-values for each region. Results show a significant association for subcortical volumes (left), but no significant relation for regional cortical surface area (middle), or thickness (right).

Supplemental Figure 3A. Sex differences in variability interacted with age in 50% of the subcortical volumes. Absolute residual values are modeled across the age range. Effects showed larger male than female variance in the younger age group, and a general trend of decreasing sex differences in variance with increasing age.

*Supplemental Figure 3B.* Sex differences in variability interacted with age in 30% of cortical surface area measures. Absolute residual values are modeled across the age range. Effects showed larger male than female variance in the younger age group, and a general trend of decreasing sex differences in variance with increasing age.

## Acknowledgements

ADHD NF-Study: The Neurofeedback study was partly funded by the project D8 of the Deutsche Forschungsgesellschaft collaborative research center 636. Barcelona 1.5T, Barcelona 3T: The Marató TV3 Foundation (#01/2010, #091710). Barcelona-Sant Pau: Miguel Servet Research Contract CPII16/0020 (Spanish Government, National Institute of Health, Carlos III); the Generalitat de Catalunya (2017SGR01343). Betula - Umea University: KA Wallenberg Foundation to LN. BIG - Nijmegen 1.5T; BIG - Nijmegen 3T: The BIG database, established in Nijmegen in 2007, is now part of Cognomics, a joint initiative by researchers of the Donders Centre of cognitive Neuroimaging, the Human Genetics and Cognitive Neuroscience departments of the Radboud university medical centre, and the Max Planck Institute for Psycholinguistics. The Cognomics Initiative is supported by the participating departments and centres and by external grants, including grants from the Biobanking and Biomolecular Resources Research Infrastructure (Netherlands) (BBMRI-NL) and the Hersenstichting Nederland. The authors also acknowledge grants supporting their work from the Netherlands Organization for Scientific Research (NWO), i.e. the NWO Brain & Cognition Excellence Program (grant 433-09-229), the Vici Innovation Program (grant 016–130-669 to BF) and #91619115. Additional support is received from the European Community’s Seventh Framework Programme (FP7/2007 – 2013) under grant agreements n° 602805 (Aggressotype), n° 603016 (MATRICS), n° 602450 (IMAGEMEND), and n° 278948 (TACTICS), and from the European Community’s Horizon 2020 Programme (H2020/2014 – 2020) under grant agreements n° 643051 (MiND) and n° 667302 (CoCA). Brain and Development Research Center, Leiden University: European Research Council (ERC-2010-StG-263234 to EAC); Research Council of Norway (#223273, #288083, #230345); South-Eastern Norway Regional Health Authority (#2017112, #2019069). BRAINSCALE: Nederlandse Organisatie voor Wetenschappelijk Onderzoek (NWO 51.02.061 to H.H., NWO 51.02.062 to D.B., NWO-NIHC Programs of excellence 433-09-220 to H.H., NWO-MagW 480-04-004 to D.B., and NWO/SPI 56-464-14192 to D.B.); FP7 Ideas: European Research Council (ERC-230374 to D.B. Universiteit Utrecht (High Potential Grant to H.H.); KNAW Academy Professor Award (PAH/6635). BRCATLAS: National Institute for Health Research (NIHR) Biomedical Research Centre at South London and Maudsley NHS Foundation Trust and King’s College London. CAMH: BBRF; Canadian Institutes of Health Research; Natural Sciences and Engineering Research Council; National Institute of Mental Health; CAMH Foundation; University of Toronto. Cardiff University: We are grateful to all researchers within Cardiff University who contributed to the MBBrains panel, and Cardiff University Brain Research Imaging Centre (CUBRIC) and the National Centre for Mental Health (NCMH) for their support. CEG (London): UK Medical Research Council Grant G03001896 to J Kuntsi; NIHR Biomedical Research Centre for Mental Health, NIHR/MRC (14/23/17); NIHR senior investigator award (NF-SI-0616-10040). CIAM: University Research Committee, University of Cape Town; National Research Foundation; South African Medical Research Council. CODE – Berlin: Lundbeck; the German Research Foundation (WA 1539/4-1, SCHN 1205/3-1). Conzelmann Study: Deutsche Forschungsgemeinschaft (KFO 125, TRR 58/A1 and A5, SFB-TRR 58/B01, B06 and Z02, RE1632/5-1); EU H2020 (#667302); German Research Foundation (KFO 125). ENIGMA Core: NIA T32AG058507; NIH/NIMH 5T32MH073526; NIH grant U54EB020403 from the Big Data to Knowledge (BD2K) Program; Core funding NIH Big Data to Knowledge (BD2K) program under consortium grant U54 EB020403; ENIGMA World Aging Center (R56 AG058854; PI PMT); ENIGMA Sex Differences Initiative (R01 MH116147; PI PMT); ENIGMA Suicidal Thoughts and Behavior Working Group (R01 MH117601; PI NJ). ENIGMA Lifespan: National Institute of Mental Health (R01MH113619, R01MH116147, R01 MH104284); National Institute for Health Research (NIHR) Biomedical Research Centre at South London and Maudsley NHS Foundation Trust and King’s College London; Psychiatry Research Trust; 2014 NARSAD Young Investigator Award. ENIGMA-HIV (NHIV; HIV-R01): NIH grant MH085604. ENIGMA-OCD (IDIBELL): FI17/00294 (Carlos III Health Institute). PI16/00889; CPII16/00048 (Carlos III Health Institute). ENIGMA-OCD (London Cohort/Mataix-Cols): Wellcome Trust and a pump priming grant from the South London and Maudsley Trust, London, UK (Project grant no. 064846). ENIGMA-OCD (van den Heuvel 1.5T; van den Heuvel 3T): The Dutch Organization for Scientific Research (NWO-ZonMw) VENI grant 916.86.036; NARSAD Young Investigators Award; Netherlands Brain Foundation grant 2010(1)-50. ENIGMA-OCD-3T-CONTROLS: South African Medical Research Council (SA MRC); South African National Research Foundation (NRF). FIDMAG: Generalitat de Catalunya (2017SGR01271); several grants funded by Instituto de Salud Carlos III co-funded by the European Regional Development Fund/European Social Fund “Investing in your future”: Miguel Servet Research Contract (CPII16/00018 to EP-C, CPII19/00009 to JR) and Research Projects (PI18/00810 to EP-C, PI18/00877 to RS, and PI19/00394 to JR); AGAUR; CIBERSAM. GSP: R01MH120080, K01MH099232, R00MH101367, R01MH119243; R01MH101486; K24 MH094614. We thank Randy Buckner for access to this dataset. Homburg Multidiagnosis Study (HMS) - Gottingen, CLING: CLING/HMS: The CliNG study sample was partially supported by the Deutsche Forschungsgemeinschaft (DFG) via the Clinical Research Group 241 ‘Genotype-phenotype relationships and neurobiology of the longitudinal course of psychosis’, TP2 (PI Gruber; http://www.kfo241.de; grant number GR 1950/5-1); data storage service SDS@hd supported by the Ministry of Science, Research and the Arts Baden-Württemberg (MWK) and the German Research Foundation (DFG) through grant INST 35/1314-1 FUGG and INST 35/1503-1 FUGG. HUBIN: Swedish Research Council (2003-5485, 2006-2992, 2006-986, 2008-2167, K2012-61X-15078-09-3, 521-2011-4622, 521-2014-3487, 2017-00949); regional agreement on medical training and clinical research between Stockholm County Council and the Karolinska Institutet; Knut and Alice Wallenberg Foundation; HUBIN project. IDIBELL: Carlos III Health Institute (PI13/01958, PI16/00889, CPII16/00048); FEDER funds/European Regional Development Fund (ERDF) - a way to build Europe-; the Department of Health of the Generalitat de Catalunya (PERIS SLT006/17/249); AGAUR (2017 SGR 1262). IMpACT-NL: The Netherlands Organization for Scientific Research (NWO), i.e. the Veni Innovation Program (grant 016-196-115 to MH) and the Vici Innovation Program (grant 016–130-669 to BF); U54 EB020403 to the ENIGMA Consortium from the BD2K Initiative, a cross-NIH partnership, and by the European College of Neuropsychopharmacology (ECNP) Network “ADHD Across the Lifespan”; The Dutch National Science Agenda NeurolabNL project (grant 400-17-602). Indiana 1.5T; Indiana 3T: NIH grants P30 AG010133, R01 AG019771 and R01 CA129769; Siemens Medical Solutions; the members of the Partnership for Pediatric Epilepsy Research, which includes the American Epilepsy Society, the Epilepsy Foundation, the Epilepsy Therapy Project, Fight Against Childhood Epilepsy and Seizures (F.A.C.E.S.), and Parents Against Childhood Epilepsy (P.A.C.E.); the GE/NFL Head Health Challenge I; the Indiana State Department of Health Spinal Cord and Brain Injury Fund Research Grant Program; a Project Development Team within the ICTSI NIH/NCRR Grant Number RR025761. Institute of Mental Health, Singapore: Singapore Bioimaging Consortium (RP C-009/06) and NMRC CSSSP (Jun17033) awarded to KS. KaSP: Swedish Medical Research Council (SE: 2009-7053; 2013-2838; SC: 523-2014-3467); the Swedish Brain Foundation; Svenska Läkaresällskapet; Torsten Söderbergs Stiftelse; Söderbergs Königska Stiftelse; Knut and Alice Wallenberg Foundation; Stockholm County Council (ALF and PPG); KID-funding from the Karolinska Institutet. MCIC: NIH P20GM103472; NIH R01EB020407; the Department of Energy DE-FG02-99ER62764 through its support of the Mind Research Network (MRN, formerly known as the MIND Institute); National Association for Research in Schizophrenia and Affective Disorders (NARSAD) Young Investigator Award (to SE); Blowitz-Ridgeway and Essel Foundations, NWO ZonMw TOP 91211021; the DFG research fellowship (to SE); the Mind Research Network, National Institutes of Health through NCRR 5 month-RR001066 (MGH General Clinical Research Center); NIMH K08 MH068540; the Biomedical Informatics Research Network with NCRR Supplements to P41 RR14075 (MGH), M01 RR 01066 (MGH), NIBIB R01EB006841 (MRN), R01EB005846 (MRN), 2R01 EB000840 (MRN), 1RC1MH089257 (MRN); U24 RR021992. METHCT: South African Medical Research Council. NETHERLANDS TWIN REGISTRY (NTR): Netherlands Organization for Scientific Research (NWO) MW904-61-193 (de Geus & Boomsma), MaGW-nr: 400-07-080 (van ‘t Ent), MagW 480-04-004 (Boomsma); NWO/SPI 56-464-14192 (Boomsma); the 646 European Research Council, ERC-230374 (Boomsma); Amsterdam Neuroscience; KNAW Academy Professor Award (PAH/6635) NeuroIMAGE: National Institutes of Health (R01MH62873 to SV Faraone); NWO Large Investment (1750102007010 to JK Buitelaar); NWO Brain & Cognition (433-09-242 to JK Buitelaar); Radboud University Medical Center, University Medical Center Groningen, Accare; VU University Amsterdam; the European Community’s Seventh Framework Programme (FP7/2007 – 2013) under grant agreements n° 602805 (Aggressotype), n° 278948 (TACTICS), and n° 602450 (IMAGEMEND); the European Community’s Horizon 2020 Programme (H2020/2014 – 2020) under grant agreements n° 643051 (MiND) and n° 667302 (CoCA); Research Council of Norway (#276082). Neuroventure: Canadian Institutes of Health Research (#287378, #FRN114887, #FRN126053). New York University: R01MH083246. Northwestern University: NIH grants P50 MH071616, R01 MH056584, R01 MH084803 (Wang PI), U01 MH097435 (Wang, Turner, Ambite, Potkin PIs), R01 EB020062 (Miller, Paulsen, Mostfosky, Wang PIs), NSF 1636893 (Pestilli, Wang, Saykin, Sporns PIs), NSF 1734853 (Pestilli, Garyfallidis, Henschel, Wang, Dinov PIs). NUI Galway: Health Research Board Ireland (HRA-POR-2013-324, HRA-POR-2011-100). Older Australian Twins Sample (OATS): NHMRC/ARC Strategic Award (ID401162); NHMRC Program Grants (ID568969, ID1093083); NHMRC Project Grants (ID1045325, ID1024224, ID1025243); we also thank Twins Research Australia. Oxford University: MRC G0500092. QTIM - University of Queensland: National Institute of Child Health and Human Development (R01 HD050735); National Health and Medical Research Council (NHMRC 486682, 1009064) Australia. São Paolo 1, São Paolo 3: Conselho Nacional de Desenvolvimento Científico e Tecnológico (CNPq, Brazil); Wellcome Trust, UK. SHIP: SHIP is part of the Community Medicine Research net of the University of Greifswald, Germany, funded by the Federal Ministry of Education and Research (grants no. 01ZZ9603, 01ZZ0103, and 01ZZ0403), the Ministry of Cultural Affairs and the Social Ministry of the Federal State of Mecklenburg-West Pomerania. MRI scans in SHIP and SHIP-TREND have been supported by a joint grant from Siemens Healthineers, Erlangen, Germany and the Federal State of Mecklenburg-West Pomerania. Stanford University: NIH Grant R37-MH101495; NIH Grant R01 MH059259 (to IHG). STROKEMRI: South-Eastern Norway Regional Health Authority (#2019107, #2015044); Norwegian ExtraFoundation for Health and Rehabilitation (#2015/FO5146). Sydney Memory and Aging Study (MAS): NHMRC Program Grants (ID350833, ID568969, ID1093083). TOP: Research Council of Norway (#223273, #248778, #249795, #300768); South-Eastern Norway Regional Health Authority (#2019-108, #2014097, #2019101); K.G. Jebsen Foundation (SKGJ-MED-008); EU (847776); European Research Council Starting Grant (#802998 to LTW); Department of Psychology, University of Oslo. UMC Utrecht 1 (CTR): zonmw 60-6360098602. University of Bari Aldo Moro (UNIBA): European Union Seventh Framework Programme for research, technological development and demonstration under grant agreement no. 602450 (IMAGEMEND) awarded to Alessandro Bertolino; “Ricerca Finalizzata” Grant PE-2011-02347951 awarded to Alessandro Bertolino; European Union’s Horizon 2020 research and innovation program under the Marie Skłodowska-Curie Grant No. 798181 (FLOURISH) awarded to Giulio Pergola. University of Bonn (Financial Risk Seeking Study): Frankfurt Institute for Risk Managemand and Regulation. University of Edinburgh: Wellcome Trust (104036/Z/14/Z, 216767/Z/19/Z); UKRI MRC (MC_PC_17209, MR/S035818/1); the European Union H2020 (SEP-210574971). University of Melbourne: National Health and Medical Research Council of Australia (NHMRC) (#1064643, #1024570); NHMRC Career Development Fellowships (1141738). University of Pennsylvania: R01MH120482; K23MH085096; R01MH101111; MH117014; MH119219. University of Sydney: NHMRC Research Fellowship. Yale University: K23 MH115206; IOCDF Award. Yale University (Olin): R01 MH106324; R01 MH096957.

## Author contributions

LMW developed the theoretical framework and prepared the manuscript with support from GED, PMT, EAC, SF, and CKT. LMW designed the models and scripts, GED and SF analyzed the data. All sites processed the imaging data and conducted quality control. GD, DD, and SF brought together and organized the datasets. *Cohort PI/ENIGMA core*: DD, IA, OAA, PA, TB, AB, DIB, SB, DB, HB, GFB, DMC, XC, TMCA, CRKC, VPC, PJC, AC, DvE, SEF, BF, ADG, DCG, IHG, HJG, OG, PG, REG, RCG, LdH, BJH, PJH, OAvdH, FMH, HEHP, CH, NJ, JAJ, AJK, JK, LL, ISL, CL, NGM, DM-C, BM, BCM, CMcD, AMM, KLM, JMM, LN, JO, PP, EP-C, MJP, JR, JLR, PGPR, MDS, PSS, TDS, AJS, KS, AS, JWS, IES, CS-M, AJS, DJS, SIT, JNT, DJV, HW, YW, BW, LTW, HCW, SCRW, MJW, MVZ, GIdZ, YW, PMT, EAC, SF. *Image data collection*: IA, TNA, AA-E, KIA, PA, SB, RB-S, AB, AB, SB, JB, AdB, AB, VDC, XC, FXC, TMCA, VPC, AC, FC, CGD, DvE, PF-C, EJCdG, ADG, DCG, IHG, HJG, PG, REG, LdH, BH, BJH, SNH, IBH, OAvdH, IBB, CAH, DJH, SH, AJH, MH, NH, FMH, CH, ACJ, EGJ, AJK, KKK, JL, LL, LdH, ISL, CL, MWJM, BM, BCM, YW, CMcD, AMM, GM, JN, YP, PP, GP, EP-C, JR, SS, AR, GR, JLR, PSS, RS, SS, TDS, AJS, MHS, KS, AS, LTS, PRS, AST, JNT, AU, N, HV, LW, YW, BW, WW, JDW, LTW, SCRW, DHW, YNY, MVZ, GCZ, EAC. *Image data processing/quality control*: GED, MA, TNA, AA-E, DA, KIA, AA, NB, SB, SE, AB, JB, AdB, RMB, VDC, EJC-R, XC, FXC, CRKC, AC, CGD, EWD, SE, DvE, JPF, PF-C, ADG, DCG, IHG, PG, TPG, BJH, SNH, OAvdH, AJH, MH, CH, ACJ, JJ, LK, BK, JL, ISL, PHL, MWJM, SM, IM-Z, BM, BCM, YW, GM, DvdM, JN, RS, EJC-R, YP, JR, GR, MDS, RS, TDS, KS, AS, LTS, PRS, SIT, AST, AU, IMV, LW, YW, WW, JDW, SCRW, KW, DHW, YNY, CKT. Manuscript revision: GED, IA, MA, AA-E, PA, AB, HB, RMB, JKB, VDC, EJC-R, XC, AC, CGD, DD, SE, PF-C, EJCdG, ADG, DCG, IHG, HJG, REG, RCG, TPG, BH, BJH, CAH, OAvdH, AJH, NH, FMH, ACJ, EGJ, JAJ, MK, JL, PHL, CL, DM-C, BM, BCM, AMM, DvdM, YP, GP, EP-C, MJP, JR, GR, PSS, RS, AJS, KS, AS, DJS, HST, AST, JNT, AU, N, HV, BW, LTW, KW, DHW.

## Collaborators

Members of the Karolinska Schizophrenia Project (KaSP) consortium: Farde L ^1^, Flyckt L ^1^, Engberg G ^2^, Erhardt S ^2^, Fatouros-Bergman H ^1^, Cervenka S ^1^, Schwieler L ^2^, Piehl F ^3^, Agartz I ^1,4,5^, Collste K ^1^, Sellgren CM ^2^, Victorsson P ^1^, Malmqvist A ^2^, Hedberg M ^2^, Orhan F ^2^. ^1^Centre for Psychiatry Research, Department of Clinical Neuroscience, Karolinska Institutet, & Stockholm Health Care Services, Stockholm County Council, Stockholm, Sweden; ^2^Department of Physiology and Pharmacology, Karolinska Institutet, Stockholm, Sweden; ^3^Neuroimmunology Unit, Department of Clinical Neuroscience, Karolinska Institutet, Stockholm, Sweden; ^4^NORMENT, Division of Mental Health and Addiction, Oslo University Hospital & Institute of Clinical Medicine, University of Oslo, Oslo, Norway; ^5^Department of Psychiatry, Diakonhjemmet Hospital, Oslo, Norway.

## Data Availability Statement

The data that support the findings of this study are available on request from the corresponding author. The data are not publicly available due to privacy or ethical restrictions.

