## Supplementary material for "Greater male than female variability in regional brain structure across the lifespan": Table 1

Table 1. Sex distributions and age of subjects by sample

| Sample | Total N | Sex | N | Age Mean | SD | Range |
|---|---|---|---|---|---|---|
| EDINBURGH | 55 | Male | 20 | 23.9 | 2.5 | 18.5 - 28.4 |
| | | Female | 35 | 23.7 | 3.1 | 18.6 - 30.6 |
| UNIBA | 131 | Male | 67 | 30.3 | 10.0 | 18.0 - 63.0 |
| | | Female | 64 | 24.3 | 6.8 | 18.0 - 52.0 |
| Tuebingen | 50 | Male | 22 | 38.4 | 11.1 | 26.0 - 61.0 |
| | | Female | 28 | 42.2 | 12.5 | 24.0 - 61.0 |
| GSP | 2009 | Male | 894 | 27.8 | 16.8 | 18.0 - 90.0 |
| | | Female | 1115 | 26.7 | 16.2 | 18.0 - 89.0 |
| Melbourne | 102 | Male | 54 | 19.5 | 2.9 | 15.0 - 25.0 |
| | | Female | 48 | 19.6 | 3.1 | 15.0 - 26.0 |
| HMS | 55 | Male | 21 | 41.3 | 11.2 | 24.0 - 59.0 |
| | | Female | 34 | 38.5 | 12.8 | 19.0 - 64.0 |
| ENIGMA-OCD (1) | 66 | Male | 30 | 30.6 | 8.9 | 19.0 - 56.0 |
| | | Female | 36 | 35.1 | 10.9 | 18.0 - 61.0 |
| NUIG | 93 | Male | 54 | 34.1 | 11.6 | 18.0 - 57.0 |
| | | Female | 39 | 39.0 | 11.0 | 18.0 - 58.0 |
| NeuroIMAGE | 383 | Male | 177 | 16.8 | 3.6 | 7.7 - 28.5 |
| | | Female | 206 | 17.0 | 3.8 | 7.8 - 28.6 |
| CAMH | 141 | Male | 72 | 43.2 | 18.9 | 18.0 - 86.0 |
| | | Female | 69 | 44.1 | 19.8 | 18.0 - 82.0 |
| Basel | 44 | Male | 17 | 25.7 | 4.5 | 19.0 - 35.0 |
| | | Female | 27 | 25.3 | 4.2 | 19.0 - 39.0 |
| Bordeaux | 452 | Male | 220 | 26.9 | 7.8 | 18.0 - 57.0 |
| | | Female | 232 | 26.6 | 7.7 | 18.0 - 56.0 |
| FBIRN | 174 | Male | 124 | 37.6 | 11.3 | 19.0 - 60.0 |
| | | Female | 50 | 37.4 | 11.3 | 19.0 - 58.0 |
| KaSP | 32 | Male | 15 | 27.4 | 5.5 | 21.0 - 43.0 |
| | | Female | 17 | 27.6 | 5.9 | 20.0 - 37.0 |
| CODE | 72 | Male | 31 | 43.7 | 12.4 | 25.0 - 64.0 |
| | | Female | 41 | 36.6 | 13.4 | 20.0 - 63.0 |
| Indiana (1) | 49 | Male | 9 | 71.9 | 6.6 | 63.0 - 80.0 |
| | | Female | 40 | 60.4 | 11.6 | 37.0 - 84.0 |
| COMPULS/TS EUROTRAIN | 53 | Male | 36 | 10.8 | 1.0 | 8.7 - 12.9 |
| | | Female | 17 | 11.0 | 1.1 | 9.2 - 12.9 |
| FIDMAG | 123 | Male | 54 | 36.4 | 8.5 | 19.0 - 63.0 |
| | | Female | 69 | 38.4 | 11.2 | 19.0 - 64.0 |
| NU | 79 | Male | 46 | 31.6 | 14.5 | 14.6 - 66.3 |
| | | Female | 33 | 34.4 | 15.3 | 14.2 - 67.9 |
| SHIP-2 | 818 | Male | 467 | 50.5 | 14.4 | 22.0 - 81.0 |
| | | Female | 351 | 49.6 | 14.0 | 21.0 - 81.0 |
| SHIP-TREND | 373 | Male | 207 | 55.6 | 12.8 | 31.0 - 84.0 |
| | | Female | 166 | 54.4 | 12.0 | 32.0 - 88.0 |
| QTIM | 340 | Male | 111 | 22.5 | 3.3 | 16.0 - 29.3 |
| | | Female | 229 | 22.7 | 3.4 | 16.1 - 30.0 |
| Betula | 287 | Male | 136 | 61.6 | 12.5 | 25.5 - 81.3 |
| | | Female | 151 | 64.1 | 13.1 | 25.7 - 80.9 |
| TOP | 303 | Male | 159 | 34.5 | 8.8 | 18.3 - 56.2 |
| | | Female | 144 | 36.3 | 10.9 | 19.3 - 73.4 |
| HUBIN | 102 | Male | 69 | 42.1 | 9.0 | 19.4 - 54.9 |
| | | Female | 33 | 41.7 | 8.5 | 19.9 - 56.2 |
| StrokeMRI | 52 | Male | 19 | 47.9 | 20.8 | 20.0 - 77.0 |
| | | Female | 33 | 43.6 | 23.0 | 18.0 - 78.0 |
| AMC | 99 | Male | 65 | 22.5 | 3.4 | 17.0 - 32.0 |
| | | Female | 34 | 23.6 | 3.3 | 18.0 - 29.0 |
| NESDA | 65 | Male | 23 | 40.7 | 9.7 | 23.0 - 56.0 |
| | | Female | 42 | 40.1 | 9.9 | 21.0 - 54.0 |
| Barcelona (1) | 30 | Male | 14 | 15.1 | 1.5 | 13.0 - 17.0 |
| | | Female | 16 | 14.9 | 2.1 | 11.0 - 17.0 |
| Barcelona (2) | 44 | Male | 24 | 14.4 | 1.8 | 11.0 - 17.0 |
| | | Female | 20 | 14.8 | 2.4 | 11.0 - 17.0 |
| Stages-Dep | 32 | Male | 9 | 46.6 | 8.4 | 37.0 - 58.0 |
| | | Female | 23 | 45.8 | 8.2 | 27.0 - 58.0 |
| IMpACT | 144 | Male | 57 | 34.2 | 11.0 | 19.0 - 62.0 |
| | | Female | 87 | 37.2 | 12.6 | 19.0 - 63.0 |
| BIG | 1319 | Male | 657 | 29.8 | 15.4 | 17.0 - 82.0 |
| | | Female | 662 | 26.9 | 12.9 | 13.0 - 79.0 |
| IMH | 56 | Male | 22 | 36.0 | 10.5 | 20.4 - 60.5 |
| Stanford | 34 | Female | 34 | 37.5 | 10.8 | 18.9 - 56.3 |
| MCIC (1) + (2) | 93 | Male | 63 | 32.8 | 12.2 | 18.0 - 58.0 |
| | | Female | 30 | 32.5 | 11.9 | 19.0 - 60.0 |
| OLIN | 599 | Male | 237 | 36.3 | 13.3 | 22.0 - 86.5 |
| | | Female | 362 | 35.9 | 12.8 | 21.0 - 74.0 |
| Neuroventure | 137 | Male | 62 | 13.7 | 0.6 | 12.4 - 14.9 |
| | | Female | 75 | 13.6 | 0.7 | 12.3 - 14.9 |
| CIAM | 30 | Male | 16 | 27.1 | 5.9 | 19.0 - 40.0 |
| | | Female | 14 | 26.1 | 3.8 | 20.0 - 33.0 |
| ENIGMA-HIV | 31 | Male | 16 | 25.6 | 4.7 | 19.0 - 33.0 |
| | | Female | 15 | 23.9 | 4.1 | 20.0 - 32.0 |
| Meth-CT | 62 | Female | 13 | 26.1 | 4.1 | 19.0 - 34.0 |
| | | Males | 49 | 27.0 | 7.9 | 18.0 - 53.0 |
| ENIGMA-OCD | 26 | Male | 10 | 34.6 | 13.6 | 19.0 - 56.0 |
| | | Female | 16 | 28.8 | 7.8 | 20.0 - 46.0 |
| Oxford | 38 | Male | 18 | 16.5 | 1.6 | 14.1 - 18.9 |
| | | Female | 20 | 15.9 | 1.1 | 13.7 - 17.7 |
| Yale | 23 | Male | 12 | 14.4 | 2.4 | 10.3 - 17.5 |
| | | Female | 11 | 14.0 | 2.0 | 9.9 - 16.5 |
| Sao Paulo-1 | 69 | Male | 45 | 27.1 | 5.6 | 18.0 - 42.0 |
| | | Female | 24 | 27.5 | 6.4 | 17.0 - 43.0 |
| Sao Paulo-3 | 85 | Male | 45 | 28.2 | 7.3 | 18.0 - 43.0 |
| | | Female | 40 | 32.7 | 8.8 | 18.0 - 50.0 |
| ENIGMA-OCD (2) | 49 | Male | 19 | 32.1 | 7.8 | 24.0 - 53.0 |
| | | Female | 30 | 31.3 | 7.7 | 21.0 - 50.0 |
| ENIGMA-OCD (3) | 35 | Male | 16 | 42.9 | 12.9 | 22.5 - 64.0 |
| | | Female | 19 | 36.0 | 8.8 | 21.5 - 49.3 |
| ENIGMA-OCD (4) | 23 | Male | 9 | 13.1 | 2.9 | 8.8 - 15.9 |
| | | Female | 14 | 13.8 | 2.4 | 8.7 - 16.8 |
| ENIGMA-OCD (5) | 33 | Male | 12 | 30.7 | 8.8 | 21.0 - 53.0 |
| | | Female | 21 | 39.2 | 11.5 | 24.0 - 63.0 |
| SYDNEY | 157 | Male | 65 | 42.0 | 22.4 | 12.0 - 84.0 |
| | | Female | 92 | 37.1 | 21.7 | 13.0 - 78.0 |
| IMH | 79 | Male | 50 | 30.7 | 8.3 | 23.0 - 53.9 |
| | | Female | 29 | 34.2 | 12.4 | 20.4 - 59.0 |
| UPENN | 187 | Male | 86 | 35.7 | 12.9 | 18.0 - 71.0 |
| | | Female | 101 | 35.8 | 14.7 | 16.0 - 85.0 |
| ADHD-NF | 13 | Male | 7 | 13.3 | 1.2 | 11.9 - 14.8 |
| | | Female | 6 | 13.4 | 0.8 | 12.1 - 14.2 |
| Indiana (2) | 66 | Male | 26 | 40.2 | 15.3 | 19.0 - 65.0 |
| | | Female | 40 | 39.4 | 14.1 | 20.0 - 65.0 |
| Sydney MAS | 523 | Male | 236 | 78.3 | 4.6 | 70.3 - 89.8 |
| | | Female | 287 | 78.5 | 4.7 | 70.5 - 90.1 |
| OADS (1) | 118 | Male | 39 | 73.8 | 5.5 | 65.0 - 84.0 |
| | | Female | 79 | 70.4 | 5.6 | 65.0 - 84.0 |
| Cardiff | 318 | Male | 89 | 28.1 | 7.8 | 19.0 - 57.0 |
| | | Female | 229 | 24.2 | 7.0 | 18.0 - 58.0 |
| CEG | 32 | Male | 32 | 15.6 | 1.7 | 13.0 - 19.0 |
| NYU | 51 | Male | 31 | 30.2 | 7.7 | 18.8 - 46.0 |
| | | Female | 20 | 31.4 | 10.3 | 19.8 - 51.9 |
| CLiNG | 321 | Male | 131 | 25.5 | 5.4 | 19.0 - 58.0 |
| | | Female | 190 | 24.9 | 5.1 | 18.0 - 57.0 |
| NTR (1) | 112 | Male | 42 | 28.5 | 8.0 | 19.0 - 56.0 |
| | | Female | 70 | 37.0 | 10.5 | 19.0 - 57.0 |
| NTR (2) | 30 | Male | 11 | 28.4 | 3.6 | 22.0 - 33.0 |
| | | Female | 19 | 28.6 | 9.8 | 1.0 - 42.0 |
| NTR (3) | 37 | Male | 14 | 15.1 | 1.5 | 12.0 - 17.0 |
| | | Female | 23 | 14.5 | 1.4 | 11.0 - 18.0 |
| Indiana (2) + (3) | 201 | Male | 97 | 21.6 | 14.4 | 6.0 - 79.0 |
| | | Female | 104 | 33.0 | 22.8 | 7.0 - 87.0 |
| BIG | 1291 | Male | 553 | 25.1 | 9.3 | 18.0 - 71.0 |
| | | Female | 738 | 23.3 | 6.9 | 18.0 - 66.0 |
| OADS (2) | 35 | Male | 15 | 70.1 | 5.7 | 65.0 - 81.0 |
| | | Female | 20 | 67.4 | 3.8 | 65.0 - 78.0 |
| OADS (3) | 153 | Male | 59 | 70.3 | 4.2 | 65.0 - 81.0 |
| | | Female | 94 | 69.7 | 4.6 | 65.0 - 81.0 |
| OADS (4) | 108 | Male | 30 | 69.8 | 4.5 | 65.0 - 85.0 |
| | | Female | 78 | 70.1 | 4.9 | 65.0 - 89.0 |
| MHRC | 52 | Male | 52 | 22.3 | 2.9 | 16.1 - 27.6 |
| BRAINSCALE | 277 | Male | 146 | 10.1 | 1.5 | 9.0 - 15.0 |
| | | Female | 131 | 9.9 | 1.2 | 9.0 - 14.1 |
| Leiden | 611 | Male | 299 | 16.2 | 4.7 | 8.3 - 28.1 |
| | | Female | 312 | 16.9 | 4.9 | 8.4 - 28.9 |
| IMAGEN | 1964 | Male | 952 | 14.5 | 0.4 | 13.2 - 15.7 |
| | | Female | 1012 | 14.5 | 0.4 | 13.3 - 16.0 |
| ENIGMA-HIV | 175 | Male | 175 | 38.8 | 6.5 | 29.0 - 50.0 |
| UMCU | 172 | Male | 84 | 40.2 | 16.5 | 18.0 - 80.0 |
| | | Female | 88 | 39.2 | 17.9 | 18.0 - 84.0 |
