## Supplementary material for "Greater male than female variability in regional brain structure across the lifespan": Table 2

### Tables Wierenga et al.

Table 2A

| Subcortical volume | Female (n=7141) | Male (n=6555) | Mean difference test | | Variance Ratio test | |
|---|---|---|---|---|---|---|
| | M | M | P | Cohen's D | VR | P |
| left thal | -328.287 | 357.024 | ** | 0.840 | 0.237 | ** |
| right thal | -317.358 | 345.963 | ** | 0.918 | 0.357 | ** |
| left caud | -139.573 | 152.488 | ** | 0.609 | 0.150 | ** |
| right caud | -147.366 | 160.706 | ** | 0.625 | 0.147 | ** |
| left put | -237.405 | 257.178 | ** | 0.757 | 0.197 | ** |
| right put | -233.415 | 252.623 | ** | 0.786 | 0.220 | ** |
| left pal | -86.166 | 93.761 | ** | 0.768 | 0.317 | ** |
| right pal | -74.910 | 81.507 | ** | 0.793 | 0.339 | ** |
| left hippo | -137.976 | 149.409 | ** | 0.673 | 0.173 | ** |
| right hippo | -134.745 | 145.724 | ** | 0.669 | 0.232 | ** |
| left amyg | -73.754 | 80.305 | ** | 0.765 | 0.154 | ** |
| right amyg | -80.242 | 87.372 | ** | 0.790 | 0.216 | ** |
| left accumb | -22.255 | 24.369 | ** | 0.414 | 0.168 | ** |
| right accumb | -22.755 | 24.685 | ** | 0.454 | 0.119 | ** |

### Table 2B

| Surface area | Female (n=6243) | Male (n=5092) | Mean difference test | | Variance Ratio test | |
|---|---|---|---|---|---|---|
| | M | M | P | Cohen's D | VR | P |
| left bankssts | -45.976 | 56.715 | ** | 0.596 | 0.282 | ** |
| left caudalanteriorcingulate | -25.875 | 31.956 | ** | 0.420 | 0.131 | ** |
| left caudalmiddlefrontal | -100.326 | 123.509 | ** | 0.589 | 0.163 | ** |
| left cuneus | -55.069 | 67.958 | ** | 0.605 | 0.188 | ** |
| left entorhinal | -19.379 | 23.824 | ** | 0.540 | 0.310 | ** |
| left fusiform | -142.081 | 174.977 | ** | 0.794 | 0.240 | ** |
| left inferiorparietal | -203.760 | 250.694 | ** | 0.751 | 0.288 | ** |
| left inferiortemporal | -158.709 | 195.821 | ** | 0.778 | 0.193 | ** |
| left isthmuscingulate | -54.544 | 67.228 | ** | 0.765 | 0.326 | ** |
| left lateraloccipital | -229.910 | 284.223 | ** | 0.893 | 0.240 | ** |
| left lateralorbitofrontal | -93.815 | 115.782 | ** | 0.771 | 0.194 | ** |
| left lingual | -114.132 | 141.130 | ** | 0.630 | 0.197 | ** |
| left medialorbitofrontal | -76.336 | 94.318 | ** | 0.741 | 0.288 | ** |
| left middletemporal | -139.909 | 172.666 | ** | 0.808 | 0.227 | ** |
| left parahippocampal | -24.273 | 30.139 | ** | 0.522 | 0.330 | ** |
| left paracentral | -46.588 | 57.790 | ** | 0.578 | 0.303 | ** |
| left parsopercularis | -63.862 | 78.461 | ** | 0.536 | 0.350 | ** |
| left parsorbitalis | -27.703 | 34.060 | ** | 0.755 | 0.223 | ** |
| left parstriangularis | -55.836 | 68.926 | ** | 0.633 | 0.262 | ** |
| left pericalcarine | -48.359 | 58.895 | ** | 0.485 | 0.151 | ** |
| left postcentral | -176.934 | 217.762 | ** | 0.867 | 0.286 | ** |
| left posteriorcingulate | -50.597 | 62.161 | ** | 0.651 | 0.253 | ** |
| left precentral | -207.652 | 255.826 | ** | 0.949 | 0.319 | ** |
| left precuneus | -163.276 | 200.728 | ** | 0.834 | 0.266 | ** |
| left rostralanteriorcingulate | -40.967 | 50.637 | ** | 0.619 | 0.160 | ** |
| left rostralmiddlefrontal | -297.267 | 365.653 | ** | 0.934 | 0.261 | ** |
| left superiorfrontal | -330.564 | 406.757 | ** | 0.962 | 0.269 | ** |
| left superiorparietal | -202.642 | 249.403 | ** | 0.730 | 0.241 | ** |
| left superiortemporal | -177.562 | 218.916 | ** | 0.970 | 0.262 | ** |
| left supramarginal | -205.547 | 254.230 | ** | 0.877 | 0.304 | ** |
| left frontalpole | -6.671 | 8.241 | ** | 0.439 | 0.249 | ** |
| left temporalpole | -15.185 | 18.664 | ** | 0.557 | 0.224 | ** |
| left transversetemporal | -19.898 | 24.463 | ** | 0.585 | 0.239 | ** |
| left insula | -84.765 | 104.782 | ** | 0.847 | 0.250 | ** |
| right bankssts | -42.654 | 52.655 | ** | 0.662 | 0.261 | ** |
| right caudalanteriorcingulate | -31.929 | 39.489 | ** | 0.465 | 0.275 | ** |
| right caudalmiddlefrontal | -95.924 | 117.705 | ** | 0.563 | 0.225 | ** |
| right cuneus | -61.606 | 75.541 | ** | 0.668 | 0.213 | ** |
| right entorhinal | -16.941 | 20.615 | ** | 0.467 | 0.339 | ** |
| right fusiform | -155.696 | 191.647 | ** | 0.900 | 0.225 | ** |
| right inferiorparietal | -278.411 | 342.870 | ** | 0.920 | 0.325 | ** |
| right inferiortemporal | -157.460 | 193.922 | ** | 0.827 | 0.187 | ** |
| right isthmuscingulate | -47.046 | 57.740 | ** | 0.723 | 0.314 | ** |
| right lateraloccipital | -227.765 | 282.023 | ** | 0.876 | 0.279 | ** |
| right lateralorbitofrontal | -99.594 | 122.823 | ** | 0.765 | 0.234 | ** |
| right lingual | -110.640 | 136.478 | ** | 0.644 | 0.225 | ** |
| right medialorbitofrontal | -70.180 | 86.695 | ** | 0.777 | 0.203 | ** |
| right middletemporal | -155.924 | 192.222 | ** | 0.857 | 0.224 | ** |
| right parahippocampal | -30.721 | 37.810 | ** | 0.708 | 0.357 | ** |
| right paracentral | -57.941 | 71.375 | ** | 0.609 | 0.349 | ** |
| right parsopercularis | -53.895 | 65.892 | ** | 0.506 | 0.312 | ** |
| right parsorbitalis | -35.086 | 43.159 | ** | 0.771 | 0.197 | ** |
| right parstriangularis | -69.557 | 85.138 | ** | 0.634 | 0.252 | ** |
| right pericalcarine | -56.327 | 68.894 | ** | 0.528 | 0.145 | ** |
| right postcentral | -168.595 | 208.307 | ** | 0.851 | 0.278 | ** |
| right posteriorcingulate | -52.836 | 65.327 | ** | 0.662 | 0.237 | ** |
| right precentral | -216.995 | 267.894 | ** | 0.950 | 0.341 | ** |
| right precuneus | -184.909 | 228.043 | ** | 0.878 | 0.248 | ** |
| right rostralanteriorcingulate | -33.179 | 41.005 | ** | 0.576 | 0.221 | ** |
| right rostralmiddlefrontal | -294.685 | 363.055 | ** | 0.898 | 0.228 | ** |
| right superiorfrontal | -325.198 | 400.002 | ** | 0.939 | 0.258 | ** |
| right superiorparietal | -205.624 | 252.962 | ** | 0.765 | 0.216 | ** |
| right superiortemporal | -132.506 | 163.787 | ** | 0.800 | 0.243 | ** |
| right supramarginal | -168.426 | 207.920 | ** | 0.754 | 0.285 | ** |
| right frontalpole | -9.712 | 11.996 | ** | 0.481 | 0.194 | ** |
| right temporalpole | -11.097 | 13.725 | ** | 0.422 | 0.228 | ** |
| right transversetemporal | -14.315 | 17.6[illegible]6 | ** | 0.564 | 0.194 | ** |
| right insula | -95.695 | 117.482 | ** | 0.863 | 0.238 | ** |

Table 2C

| Thickness | Female (n=6620) | Male (n=5913) | Mean difference test | | Variance Ratio test | |
|---|---|---|---|---|---|---|
| | M | M | P | Cohen's D | VR | P |
| left bankssts | 0.001 | -0.001 | n.s. | 0.011 | 0.039 | ** |
| left caudalanteriorcingulate | 0.026 | -0.028 | ** | 0.213 | -0.042 | n.s. |
| left caudalmiddlefrontal | 0.008 | -0.008 | ** | 0.103 | 0.061 | * |
| left cuneus | 0.000 | 0.000 | n.s. | 0.001 | 0.050 | * |
| left entorhinal | -0.013 | 0.015 | ** | 0.084 | 0.023 | n.s. |
| left fusiform | 0.001 | -0.001 | n.s. | 0.016 | 0.022 | n.s. |
| left inferiorparietal | 0.009 | -0.009 | ** | 0.128 | 0.092 | ** |
| left inferiortemporal | -0.002 | 0.003 | n.s. | 0.027 | 0.004 | n.s. |
| left isthmuscingulate | 0.009 | -0.009 | ** | 0.088 | -0.007 | ** |
| left lateraloccipital | 0.005 | -0.005 | ** | 0.074 | 0.079 | ** |
| left lateralorbitofrontal | -0.002 | 0.003 | n.s. | 0.036 | 0.101 | ** |
| left lingual | -0.003 | 0.004 | ** | 0.058 | 0.040 | n.s. |
| left medialorbitofrontal | -0.004 | 0.006 | ** | 0.058 | 0.027 | n.s. |
| left middletemporal | -0.003 | 0.004 | n.s. | 0.037 | 0.093 | * |
| left parahippocampal | 0.015 | -0.016 | ** | 0.098 | 0.016 | n.s. |
| left paracentral | 0.006 | -0.005 | ** | 0.067 | 0.030 | ** |
| left parsopercularis | -0.002 | 0.003 | n.s. | 0.027 | 0.087 | ** |
| left parsorbitalis | 0.013 | -0.014 | ** | 0.120 | 0.071 | ** |
| left parstriangularis | 0.004 | -0.004 | * | 0.049 | 0.084 | ** |
| left pericalcarine | 0.000 | 0.001 | n.s. | 0.006 | 0.043 | ** |
| left postcentral | 0.008 | -0.009 | ** | 0.133 | 0.078 | ** |
| left posteriorcingulate | 0.004 | -0.004 | ** | 0.052 | 0.080 | ** |
| left precentral | 0.007 | -0.007 | ** | 0.097 | 0.112 | ** |
| left precuneus | 0.000 | 0.000 | n.s. | 0.002 | 0.041 | ** |
| left rostralanteriorcingulate | 0.020 | -0.021 | ** | 0.170 | -0.046 | n.s. |
| left rostralmiddlefrontal | 0.005 | -0.004 | ** | 0.061 | 0.112 | ** |
| left superiorfrontal | 0.013 | -0.014 | ** | 0.168 | 0.048 | n.s. |
| left superiorparietal | 0.009 | -0.009 | ** | 0.136 | 0.098 | ** |
| left superiortemporal | -0.001 | 0.001 | n.s. | 0.014 | 0.052 | ** |
| left supramarginal | 0.009 | -0.009 | ** | 0.126 | 0.064 | ** |
| left frontalpole | 0.015 | -0.016 | ** | 0.100 | 0.036 | n.s. |
| left temporalpole | 0.004 | -0.004 | n.s. | 0.023 | 0.027 | n.s. |
| left transversetemporal | 0.020 | -0.021 | ** | 0.177 | 0.018 | n.s. |
| left insula | -0.009 | 0.011 | ** | 0.121 | 0.049 | n.s. |
| right bankssts | -0.001 | 0.002 | n.s. | 0.016 | 0.064 | ** |
| right caudalanteriorcingulate | 0.027 | -0.030 | ** | 0.242 | -0.029 | n.s. |
| right caudalmiddlefrontal | 0.008 | -0.009 | ** | 0.109 | 0.019 | ** |
| right cuneus | 0.003 | -0.002 | n.s. | 0.034 | 0.027 | * |
| right entorhinal | 0.005 | -0.005 | n.s. | 0.028 | 0.026 | n.s. |
| right fusiform | 0.001 | 0.000 | n.s. | 0.008 | 0.029 | n.s. |
| right inferiorparietal | 0.008 | -0.008 | ** | 0.110 | 0.103 | ** |
| right inferiortemporal | 0.000 | 0.001 | n.s. | 0.003 | 0.032 | n.s. |
| right isthmuscingulate | 0.010 | -0.010 | ** | 0.099 | -0.038 | ** |
| right lateraloccipital | 0.004 | -0.004 | ** | 0.057 | 0.078 | ** |
| right lateralorbitofrontal | 0.003 | -0.003 | n.s. | 0.036 | 0.074 | ** |
| right lingual | -0.002 | 0.003 | n.s. | 0.036 | 0.036 | n.s. |
| right medialorbitofrontal | 0.003 | -0.003 | n.s. | 0.033 | 0.056 | n.s. |
| right middletemporal | -0.003 | 0.004 | * | 0.047 | 0.065 | ** |
| right parahippocampal | 0.021 | -0.023 | ** | 0.162 | 0.028 | n.s. |
| right paracentral | 0.004 | -0.004 | ** | 0.055 | 0.065 | ** |
| right parsopercularis | 0.000 | 0.000 | n.s. | 0.001 | 0.037 | ** |
| right parsorbitalis | 0.018 | -0.019 | ** | 0.164 | 0.026 | n.s. |
| right parstriangularis | 0.004 | -0.004 | ** | 0.053 | 0.008 | ** |
| right pericalcarine | 0.001 | -0.001 | n.s. | 0.017 | 0.020 | n.s. |
| right postcentral | 0.009 | -0.009 | ** | 0.135 | 0.009 | ** |
| right posteriorcingulate | 0.007 | -0.007 | ** | 0.082 | 0.013 | ** |
| right precentral | 0.008 | -0.009 | ** | 0.119 | 0.084 | ** |
| right precuneus | -0.001 | 0.002 | n.s. | 0.018 | 0.063 | ** |
| right rostralanteriorcingulate | 0.009 | -0.010 | ** | 0.080 | 0.055 | n.s. |
| right rostralmiddlefrontal | 0.006 | -0.006 | ** | 0.078 | 0.085 | ** |
| right superiorfrontal | 0.013 | -0.013 | ** | 0.165 | 0.065 | * |
| right superiorparietal | 0.008 | -0.009 | ** | 0.132 | 0.065 | ** |
| right superiortemporal | -0.003 | 0.004 | * | 0.042 | 0.073 | ** |
| right supramarginal | 0.006 | -0.007 | ** | 0.086 | 0.096 | ** |
| right frontalpole | 0.021 | -0.022 | ** | 0.140 | 0.012 | n.s. |
| right temporalpole | -0.006 | 0.007 | * | 0.038 | 0.023 | n.s. |
| right transversetemporal | 0.011 | -0.011 | ** | 0.095 | 0.101 | * |
| right insula | -0.008 | 0.010 | ** | 0.107 | 0.092 | ** |
