## Supplementary material for "Greater male than female variability in regional brain structure across the lifespan": Table 3

Table 3A

| Subcortical | Intercept | (s.e.) | P | Age | (s.e.) | P | Sex | (s.e.) | P | Sex by age | (s.e.) | P |
|---|---|---|---|---|---|---|---|---|---|---|---|---|
| left thal | 587.987 | 6.178 | ** | 9398.523 | 652.185 | ** | 60.310 | 9.199 | ** | -3107.885 | 979.201 | ** |
| right thal | 515.416 | 5.524 | ** | 6424.232 | 583.119 | ** | 82.380 | 8.225 | ** | -3102.267 | 875.503 | ** |
| left caud | 361.790 | 3.729 | ** | 879.545 | 393.693 | * | 28.152 | 5.553 | ** | 270.769 | 591.096 | n.s. |
| right caud | 371.773 | 3.785 | ** | 1290.352 | 399.567 | ** | 31.395 | 5.636 | ** | -561.719 | 599.915 | n.s. |
| left put | 495.399 | 5.150 | ** | 4435.730 | 543.701 | ** | 54.586 | 7.669 | ** | -2966.533 | 816.321 | ** |
| right put | 460.842 | 4.887 | ** | 5622.177 | 515.939 | ** | 51.687 | 7.277 | ** | -3853.454 | 774.638 | ** |
| left pal | 165.039 | 1.816 | ** | 837.030 | 191.768 | ** | 26.852 | 2.705 | ** | -784.363 | 287.923 | * |
| right pal | 140.799 | 1.598 | ** | 910.463 | 168.695 | ** | 26.247 | 2.379 | ** | -850.994 | 253.281 | ** |
| left hippo | 309.722 | 3.308 | ** | 2755.892 | 349.231 | ** | 31.626 | 4.926 | ** | -1375.500 | 524.341 | * |
| right hippo | 305.607 | 3.264 | ** | 2615.969 | 344.571 | ** | 35.732 | 4.860 | ** | -890.970 | 517.345 | n.s. |
| left amyg | 148.932 | 1.598 | ** | 1378.267 | 168.734 | ** | 13.800 | 2.380 | ** | -233.236 | 253.340 | n.s. |
| right amyg | 154.218 | 1.645 | ** | 1621.298 | 173.675 | ** | 16.477 | 2.450 | ** | -540.141 | 260.758 | n.s. |
| left accumb | 82.473 | 0.875 | ** | 442.922 | 92.410 | ** | 7.382 | 1.303 | ** | -136.472 | 138.746 | n.s. |
| right accumb | 78.541 | 0.823 | ** | 539.975 | 86.850 | ** | 7.412 | 1.225 | ** | -106.522 | 130.398 | n.s. |

### Table 3B

| Surface area | Intercept | (s.e.) | P | Age | (s.e.) | P | Sex | (s.e.) | P | Sex by age | (s.e.) | P |
|---|---|---|---|---|---|---|---|---|---|---|---|---|
| left bankssts | 127.133 | 1.376 | ** | -437.616 | 142.554 | ** | 16.563 | 2.056 | ** | -574.105 | 219.785 | * |
| left caudalanteriorcingulate | 104.209 | 1.113 | ** | -302.669 | 115.254 | ** | 4.299 | 1.663 | ** | -277.614 | 177.695 | n.s. |
| left caudalmiddlefrontal | 293.750 | 2.943 | ** | -1359.284 | 304.791 | ** | 21.272 | 4.397 | ** | -660.300 | 469.918 | n.s. |
| left cuneus | 154.129 | 1.607 | ** | -360.698 | 166.430 | * | 13.158 | 2.401 | ** | -330.457 | 256.596 | n.s. |
| left entorhinal | 57.126 | 0.651 | ** | -458.398 | 67.397 | ** | 9.241 | 0.972 | ** | 1.893 | 103.911 | n.s. |
| left fusiform | 305.090 | 3.105 | ** | 250.591 | 321.575 | n.s. | 35.738 | 4.639 | ** | -2446.584 | 495.794 | ** |
| left inferiorparietal | 454.916 | 4.708 | ** | -614.521 | 487.682 | n.s. | 63.459 | 7.035 | ** | -2243.805 | 751.894 | * |
| left inferiortemporal | 352.394 | 3.540 | ** | -353.703 | 366.628 | n.s. | 31.482 | 5.289 | ** | -1652.239 | 565.256 | * |
| left isthmuscingulate | 116.771 | 1.249 | ** | -32.188 | 129.411 | n.s. | 19.544 | 1.867 | ** | -204.545 | 199.522 | n.s. |
| left lateraloccipital | 438.089 | 4.474 | ** | -1416.631 | 463.377 | ** | 50.571 | 6.685 | ** | -813.654 | 714.421 | n.s. |
| left lateralorbitofrontal | 208.173 | 2.120 | ** | 204.108 | 219.597 | n.s. | 20.633 | 3.168 | ** | -1428.745 | 338.567 | ** |
| left lingual | 310.573 | 3.141 | ** | -234.334 | 325.364 | n.s. | 29.898 | 4.694 | ** | -1268.288 | 501.636 | * |
| left medialorbitofrontal | 172.506 | 1.795 | ** | 3.188 | 185.938 | n.s. | 23.450 | 2.682 | ** | -213.946 | 286.673 | n.s. |
| left middletemporal | 296.794 | 2.997 | ** | -421.492 | 310.480 | n.s. | 31.627 | 4.479 | ** | -1014.822 | 478.689 | n.s. |
| left parahippocampal | 72.669 | 0.887 | ** | -211.577 | 91.839 | * | 10.825 | 1.325 | ** | -241.097 | 141.595 | n.s. |
| left paracentral | 133.446 | 1.419 | ** | -195.857 | 147.019 | n.s. | 19.139 | 2.121 | ** | -171.708 | 226.670 | n.s. |
| left parsopercularis | 193.582 | 2.113 | ** | -540.023 | 218.880 | * | 31.583 | 3.158 | ** | -459.911 | 337.462 | n.s. |
| left parsorbitalis | 61.886 | 0.643 | ** | -172.940 | 66.566 | ** | 7.120 | 0.960 | ** | -131.612 | 102.629 | n.s. |
| left parstriangularis | 148.566 | 1.524 | ** | -644.966 | 157.820 | ** | 19.173 | 2.277 | ** | -546.829 | 243.322 | n.s. |
| left pericalcarine | 171.607 | 1.690 | ** | -245.127 | 175.004 | n.s. | 13.803 | 2.525 | ** | -283.583 | 269.815 | n.s. |
| left postcentral | 340.927 | 3.572 | ** | -1033.492 | 370.007 | ** | 46.097 | 5.338 | ** | -1240.366 | 570.466 | n.s. |
| left posteriorcingulate | 130.459 | 1.363 | ** | -176.189 | 141.217 | n.s. | 13.905 | 2.037 | ** | -400.954 | 217.724 | n.s. |
| left precentral | 360.893 | 3.926 | ** | -1088.967 | 406.693 | ** | 47.580 | 5.867 | ** | -876.707 | 627.028 | n.s. |
| left precuneus | 329.439 | 3.386 | ** | -444.670 | 350.720 | n.s. | 44.718 | 5.060 | ** | -1691.713 | 540.730 | * |
| left rostralanteriorcingulate | 113.700 | 1.156 | ** | -6.807 | 119.754 | n.s. | 7.691 | 1.728 | ** | -80.447 | 184.632 | n.s. |
| left rostralmiddlefrontal | 541.319 | 5.553 | ** | -1574.677 | 575.208 | ** | 63.888 | 8.298 | ** | -2391.074 | 886.838 | * |
| left superiorfrontal | 577.465 | 6.015 | ** | -1306.494 | 623.063 | * | 75.007 | 8.988 | ** | -2320.740 | 960.620 | n.s. |
| left superiorparietal | 471.735 | 4.793 | ** | -1198.240 | 496.487 | * | 57.076 | 7.162 | ** | -2051.708 | 765.468 | * |
| left superiortemporal | 308.552 | 3.215 | ** | -864.236 | 333.037 | ** | 40.486 | 4.804 | ** | -1222.034 | 513.467 | n.s. |
| left supramarginal | 392.296 | 4.082 | ** | -1937.799 | 422.787 | ** | 58.041 | 6.099 | ** | -775.470 | 651.841 | n.s. |
| left frontalpole | 25.431 | 0.265 | ** | -114.432 | 27.425 | ** | 3.212 | 0.396 | ** | -7.992 | 42.283 | n.s. |
| left temporalpole | 45.410 | 0.478 | ** | -173.235 | 49.555 | ** | 5.115 | 0.715 | ** | -59.323 | 76.403 | n.s. |
| left transversetemporal | 56.992 | 0.594 | ** | -201.824 | 61.535 | ** | 6.690 | 0.888 | ** | -81.655 | 94.872 | n.s. |
| left insula | 164.339 | 1.842 | ** | -460.767 | 190.830 | * | 17.215 | 2.753 | ** | 6.824 | 294.215 | n.s. |
| right bankssts | 107.290 | 1.139 | ** | -392.600 | 117.986 | ** | 13.575 | 1.702 | ** | -493.453 | 181.908 | * |
| right caudalanteriorcingulate | 114.549 | 1.199 | ** | -266.524 | 124.192 | * | 14.948 | 1.792 | ** | -8.218 | 191.475 | n.s. |
| right caudalmiddlefrontal | 288.671 | 2.929 | ** | -1415.348 | 303.395 | ** | 30.576 | 4.377 | ** | -360.883 | 467.765 | n.s. |
| right cuneus | 152.647 | 1.656 | ** | -146.322 | 171.565 | n.s. | 16.151 | 2.475 | ** | -436.462 | 264.513 | n.s. |
| right entorhinal | 57.865 | 0.641 | ** | -455.979 | 66.351 | ** | 10.302 | 0.957 | ** | -50.231 | 102.298 | n.s. |
| right fusiform | 295.259 | 3.000 | ** | 43.695 | 310.723 | n.s. | 32.408 | 4.483 | ** | -1812.528 | 479.064 | ** |
| right inferiorparietal | 504.767 | 5.239 | ** | -577.142 | 542.646 | n.s. | 82.015 | 7.828 | ** | -2767.949 | 836.635 | ** |
| right inferiortemporal | 327.236 | 3.331 | ** | -482.481 | 345.043 | n.s. | 28.512 | 4.978 | ** | -1116.568 | 531.977 | n.s. |
| right isthmuscingulate | 105.700 | 1.157 | ** | -228.263 | 119.818 | n.s. | 16.311 | 1.729 | ** | -192.830 | 184.732 | n.s. |
| right lateraloccipital | 436.925 | 4.537 | ** | -1283.916 | 469.975 | ** | 58.726 | 6.780 | ** | -1927.057 | 724.593 | * |
| right lateralorbitofrontal | 220.527 | 2.284 | ** | 236.472 | 236.616 | n.s. | 24.442 | 3.413 | ** | -1470.759 | 364.808 | ** |
| right lingual | 289.568 | 3.001 | ** | -299.806 | 310.855 | n.s. | 34.596 | 4.484 | ** | -1128.138 | 479.266 | n.s. |
| right medialorbitofrontal | 154.743 | 1.568 | ** | 74.312 | 162.424 | n.s. | 15.452 | 2.343 | ** | -964.430 | 250.420 | ** |
| right middletemporal | 309.733 | 3.171 | ** | -517.078 | 328.408 | n.s. | 34.194 | 4.738 | ** | -1188.068 | 506.329 | n.s. |
| right parahippocampal | 70.171 | 0.781 | ** | -155.100 | 80.940 | n.s. | 11.822 | 1.168 | ** | -420.498 | 124.790 | ** |
| right paracentral | 156.024 | 1.669 | ** | -273.907 | 172.868 | n.s. | 25.570 | 2.494 | ** | -271.297 | 266.523 | n.s. |
| right parsopercularis | 174.570 | 1.866 | ** | -1036.595 | 193.296 | ** | 25.454 | 2.789 | ** | -231.029 | 298.018 | n.s. |
| right parsorbitalis | 77.607 | 0.794 | ** | -103.424 | 82.287 | n.s. | 7.160 | 1.187 | ** | -311.879 | 126.867 | * |
| right parstriangularis | 184.989 | 1.887 | ** | -925.697 | 195.494 | ** | 21.344 | 2.820 | ** | -662.628 | 301.407 | n.s. |
| right pericalcarine | 184.490 | 1.818 | ** | -314.748 | 188.350 | n.s. | 13.276 | 2.717 | ** | -264.356 | 290.392 | n.s. |
| right postcentral | 330.886 | 3.494 | ** | -1175.639 | 361.875 | ** | 44.061 | 5.220 | ** | -907.204 | 557.928 | n.s. |
| right posteriorcingulate | 133.953 | 1.413 | ** | 42.583 | 146.371 | n.s. | 14.739 | 2.112 | ** | -695.150 | 225.670 | * |
| right precentral | 374.619 | 4.131 | ** | -1039.063 | 427.849 | * | 53.576 | 6.172 | ** | -579.997 | 659.645 | n.s. |
| right precuneus | 355.783 | 3.685 | ** | -894.373 | 381.705 | * | 42.292 | 5.507 | ** | -1788.652 | 588.501 | * |
| right rostralanteriorcingulate | 97.009 | 1.005 | ** | 198.486 | 104.078 | n.s. | 10.668 | 1.501 | ** | -140.756 | 160.464 | n.s. |
| right rostralmiddlefrontal | 560.924 | 5.691 | ** | -2015.333 | 589.514 | ** | 60.682 | 8.504 | ** | -1467.830 | 908.895 | n.s. |
| right superiorfrontal | 586.059 | 6.054 | ** | -748.583 | 627.121 | n.s. | 72.274 | 9.047 | ** | -3613.685 | 966.876 | ** |
| right superiorparietal | 453.081 | 4.716 | ** | -1983.725 | 488.528 | ** | 49.530 | 7.048 | ** | 42.170 | 753.197 | n.s. |
| right superiortemporal | 281.023 | 2.898 | ** | -481.481 | 300.133 | n.s. | 31.844 | 4.330 | ** | -1005.995 | 462.736 | n.s. |
| right supramarginal | 376.538 | 3.839 | ** | -1315.029 | 397.627 | ** | 51.001 | 5.736 | ** | -1362.209 | 613.049 | n.s. |
| right frontalpole | 34.322 | 0.352 | ** | -93.541 | 36.451 | * | 2.974 | 0.526 | ** | -112.046 | 56.199 | n.s. |
| right temporalpole | 44.173 | 0.457 | ** | -144.791 | 47.330 | ** | 5.067 | 0.683 | ** | -32.370 | 72.972 | n.s. |
| right transversetemporal | 43.342 | 0.436 | ** | -122.601 | 45.112 | ** | 4.348 | 0.651 | ** | -76.872 | 69.553 | n.s. |
| right insula | 185.386 | 1.947 | ** | 167.564 | 201.684 | n.s. | 22.970 | 2.910 | ** | -270.419 | 310.950 | n.s. |

#### Table 3C

| Thickness | Intercept | (s.e.) | P | Age | (s.e.) | P | Sex | (s.e.) | P | Sex by age | (s.e.) | P |
|---|---|---|---|---|---|---|---|---|---|---|---|---|
| left bankssts | 0.138 | 0.001 | ** | 0.012 | 0.150 | n.s. | 0.002 | 0.002 | n.s. | 0.345 | 0.217 | n.s. |
| left caudalanteriorcingulate | 0.204 | 0.002 | ** | 1.405 | 0.217 | ** | -0.005 | 0.003 | n.s. | 0.207 | 0.314 | n.s. |
| left caudalmiddlefrontal | 0.119 | 0.001 | ** | 0.375 | 0.131 | ** | 0.002 | 0.002 | n.s. | -0.108 | 0.190 | n.s. |
| left cuneus | 0.108 | 0.001 | ** | -0.194 | 0.118 | n.s. | 0.003 | 0.002 | n.s. | -0.386 | 0.171 | n.s. |
| left entorhinal | 0.263 | 0.003 | ** | 0.348 | 0.288 | n.s. | 0.001 | 0.004 | n.s. | -0.414 | 0.417 | n.s. |
| left fusiform | 0.114 | 0.001 | ** | 0.484 | 0.125 | ** | 0.000 | 0.002 | n.s. | -0.340 | 0.181 | n.s. |
| left inferiorparietal | 0.109 | 0.001 | ** | 0.329 | 0.122 | ** | 0.005 | 0.002 | ** | 0.023 | 0.176 | n.s. |
| left inferiortemporal | 0.128 | 0.001 | ** | 0.515 | 0.138 | ** | 0.000 | 0.002 | n.s. | -0.327 | 0.199 | n.s. |
| left isthmuscingulate | 0.165 | 0.002 | ** | 0.491 | 0.175 | ** | -0.003 | 0.002 | n.s. | -0.076 | 0.254 | n.s. |
| left lateraloccipital | 0.096 | 0.001 | ** | 0.132 | 0.106 | n.s. | 0.004 | 0.001 | ** | 0.057 | 0.154 | n.s. |
| left lateralorbitofrontal | 0.124 | 0.001 | ** | 0.212 | 0.138 | n.s. | 0.006 | 0.002 | ** | -0.438 | 0.201 | n.s. |
| left lingual | 0.099 | 0.001 | ** | 0.343 | 0.109 | ** | 0.001 | 0.001 | n.s. | -0.308 | 0.157 | n.s. |
| left medialorbitofrontal | 0.135 | 0.001 | ** | 0.067 | 0.150 | n.s. | 0.004 | 0.002 | n.s. | -0.425 | 0.217 | n.s. |
| left middletemporal | 0.129 | 0.001 | ** | 0.493 | 0.140 | ** | 0.004 | 0.002 | * | -0.012 | 0.203 | n.s. |
| left parahippocampal | 0.248 | 0.002 | ** | 0.441 | 0.254 | n.s. | 0.002 | 0.003 | n.s. | -0.372 | 0.368 | n.s. |
| left paracentral | 0.126 | 0.001 | ** | 0.321 | 0.138 | * | 0.003 | 0.002 | n.s. | -0.017 | 0.199 | n.s. |
| left parsopercularis | 0.123 | 0.001 | ** | 0.497 | 0.134 | ** | 0.005 | 0.002 | ** | -0.358 | 0.194 | n.s. |
| left parsorbitalis | 0.178 | 0.002 | ** | -0.413 | 0.192 | * | 0.004 | 0.003 | n.s. | 0.266 | 0.278 | n.s. |
| left parstriangularis | 0.134 | 0.001 | ** | 0.145 | 0.144 | n.s. | 0.004 | 0.002 | * | -0.073 | 0.209 | n.s. |
| left pericalcarine | 0.101 | 0.001 | ** | 0.202 | 0.114 | n.s. | 0.001 | 0.002 | n.s. | -0.325 | 0.165 | n.s. |
| left postcentral | 0.097 | 0.001 | ** | 0.340 | 0.106 | ** | 0.004 | 0.001 | ** | 0.222 | 0.154 | n.s. |
| left posteriorcingulate | 0.131 | 0.001 | ** | 0.308 | 0.142 | * | 0.005 | 0.002 | ** | -0.236 | 0.205 | n.s. |
| left precentral | 0.110 | 0.001 | ** | 1.223 | 0.122 | ** | 0.004 | 0.002 | * | 0.181 | 0.177 | n.s. |
| left precuneus | 0.111 | 0.001 | ** | 0.521 | 0.121 | ** | 0.003 | 0.002 | n.s. | -0.056 | 0.176 | n.s. |
| left rostralanteriorcingulate | 0.193 | 0.002 | ** | 0.470 | 0.205 | * | -0.005 | 0.003 | n.s. | -0.378 | 0.298 | n.s. |
| left rostralmiddlefrontal | 0.109 | 0.001 | ** | 0.153 | 0.122 | n.s. | 0.005 | 0.002 | ** | 0.039 | 0.177 | n.s. |
| left superiorfrontal | 0.124 | 0.001 | ** | 0.505 | 0.137 | ** | 0.002 | 0.002 | n.s. | 0.083 | 0.198 | n.s. |
| left superiorparietal | 0.099 | 0.001 | ** | 0.158 | 0.109 | n.s. | 0.004 | 0.001 | ** | 0.224 | 0.158 | n.s. |
| left superiortemporal | 0.129 | 0.001 | ** | 0.832 | 0.139 | ** | 0.004 | 0.002 | * | -0.123 | 0.201 | n.s. |
| left supramarginal | 0.114 | 0.001 | ** | 0.396 | 0.122 | ** | 0.005 | 0.002 | ** | 0.063 | 0.177 | n.s. |
| left frontalpole | 0.241 | 0.002 | ** | -1.236 | 0.266 | ** | 0.004 | 0.004 | n.s. | 0.112 | 0.386 | n.s. |
| left temporalpole | 0.268 | 0.003 | ** | -2.010 | 0.301 | ** | 0.006 | 0.004 | n.s. | -0.518 | 0.436 | n.s. |
| left transversetemporal | 0.182 | 0.002 | ** | 0.027 | 0.194 | n.s. | -0.001 | 0.003 | n.s. | -0.168 | 0.281 | n.s. |
| left insula | 0.125 | 0.001 | ** | 1.184 | 0.135 | ** | 0.002 | 0.002 | n.s. | -0.700 | 0.195 | * |
| right bankssts | 0.146 | 0.001 | ** | -0.094 | 0.157 | n.s. | 0.003 | 0.002 | n.s. | 0.217 | 0.228 | n.s. |
| right caudalanteriorcingulate | 0.186 | 0.002 | ** | 0.936 | 0.198 | ** | -0.008 | 0.003 | ** | -0.105 | 0.288 | n.s. |
| right caudalmiddlefrontal | 0.120 | 0.001 | ** | 0.226 | 0.130 | n.s. | 0.002 | 0.002 | n.s. | 0.179 | 0.189 | n.s. |
| right cuneus | 0.110 | 0.001 | ** | 0.037 | 0.118 | n.s. | 0.001 | 0.002 | n.s. | -0.334 | 0.170 | n.s. |
| right entorhinal | 0.288 | 0.003 | ** | 0.122 | 0.310 | n.s. | 0.004 | 0.004 | n.s. | -0.746 | 0.449 | n.s. |
| right fusiform | 0.114 | 0.001 | ** | 0.657 | 0.125 | ** | 0.001 | 0.002 | n.s. | -0.171 | 0.181 | n.s. |
| right inferiorparietal | 0.109 | 0.001 | ** | 0.390 | 0.120 | ** | 0.005 | 0.002 | ** | 0.233 | 0.174 | n.s. |
| right inferiortemporal | 0.124 | 0.001 | ** | 0.539 | 0.135 | ** | 0.003 | 0.002 | n.s. | -0.132 | 0.196 | n.s. |
| right isthmuscingulate | 0.162 | 0.002 | ** | 0.401 | 0.172 | * | -0.002 | 0.002 | n.s. | 0.223 | 0.249 | n.s. |
| right lateraloccipital | 0.101 | 0.001 | ** | 0.280 | 0.110 | * | 0.005 | 0.001 | ** | 0.023 | 0.159 | n.s. |
| right lateralorbitofrontal | 0.129 | 0.001 | ** | -0.174 | 0.144 | n.s. | 0.004 | 0.002 | * | -0.110 | 0.208 | n.s. |
| right lingual | 0.102 | 0.001 | ** | 0.172 | 0.111 | n.s. | 0.000 | 0.002 | n.s. | -0.201 | 0.161 | n.s. |
| right medialorbitofrontal | 0.142 | 0.001 | ** | -0.424 | 0.156 | ** | 0.003 | 0.002 | n.s. | -0.201 | 0.227 | n.s. |
| right middletemporal | 0.123 | 0.001 | ** | 0.067 | 0.137 | n.s. | 0.006 | 0.002 | ** | 0.400 | 0.198 | n.s. |
| right parahippocampal | 0.207 | 0.002 | ** | 0.554 | 0.224 | * | 0.005 | 0.003 | n.s. | -0.115 | 0.325 | n.s. |
| right paracentral | 0.124 | 0.001 | ** | 0.492 | 0.134 | ** | 0.002 | 0.002 | n.s. | -0.050 | 0.194 | n.s. |
| right parsopercularis | 0.131 | 0.001 | ** | 0.330 | 0.139 | * | 0.001 | 0.002 | n.s. | -0.056 | 0.201 | n.s. |
| right parsorbitalis | 0.175 | 0.002 | ** | -0.470 | 0.188 | * | 0.002 | 0.003 | n.s. | 0.159 | 0.273 | n.s. |
| right parstriangularis | 0.131 | 0.001 | ** | -0.016 | 0.141 | n.s. | 0.002 | 0.002 | n.s. | 0.052 | 0.204 | n.s. |
| right pericalcarine | 0.102 | 0.001 | ** | 0.199 | 0.112 | n.s. | 0.002 | 0.002 | n.s. | -0.336 | 0.163 | n.s. |
| right postcentral | 0.102 | 0.001 | ** | 0.121 | 0.111 | n.s. | 0.002 | 0.002 | n.s. | 0.251 | 0.161 | n.s. |
| right posteriorcingulate | 0.129 | 0.001 | ** | 0.442 | 0.139 | ** | 0.000 | 0.002 | n.s. | -0.014 | 0.202 | n.s. |
| right precentral | 0.110 | 0.001 | ** | 0.992 | 0.124 | ** | 0.005 | 0.002 | ** | 0.411 | 0.179 | n.s. |
| right precuneus | 0.110 | 0.001 | ** | 0.473 | 0.121 | ** | 0.004 | 0.002 | * | -0.148 | 0.176 | n.s. |
| right rostralanteriorcingulate | 0.185 | 0.002 | ** | 0.390 | 0.205 | n.s. | 0.009 | 0.003 | ** | -0.713 | 0.298 | n.s. |
| right rostralmiddlefrontal | 0.108 | 0.001 | ** | 0.084 | 0.120 | n.s. | 0.003 | 0.002 | n.s. | -0.162 | 0.174 | n.s. |
| right superiorfrontal | 0.120 | 0.001 | ** | 0.499 | 0.131 | ** | 0.003 | 0.002 | n.s. | -0.189 | 0.190 | n.s. |
| right superiorparietal | 0.099 | 0.001 | ** | 0.231 | 0.110 | * | 0.003 | 0.002 | * | 0.154 | 0.160 | n.s. |
| right superiortemporal | 0.127 | 0.001 | ** | 0.738 | 0.138 | ** | 0.005 | 0.002 | * | 0.153 | 0.201 | n.s. |
| right supramarginal | 0.117 | 0.001 | ** | 0.723 | 0.127 | ** | 0.004 | 0.002 | * | -0.037 | 0.184 | n.s. |
| right frontalpole | 0.236 | 0.002 | ** | -0.642 | 0.255 | * | 0.002 | 0.003 | n.s. | -0.248 | 0.369 | n.s. |
| right temporalpole | 0.274 | 0.003 | ** | -2.088 | 0.317 | ** | 0.007 | 0.004 | n.s. | 0.219 | 0.459 | n.s. |
| right transversetemporal | 0.181 | 0.002 | ** | 0.511 | 0.198 | * | 0.010 | 0.003 | ** | -0.175 | 0.287 | n.s. |
| right insula | 0.130 | 0.001 | ** | 1.079 | 0.146 | ** | 0.005 | 0.002 | * | -0.468 | 0.211 | n.s. |
