## Supplemental Figure 1 for "Greater male than female variability in regional brain structure across the lifespan"

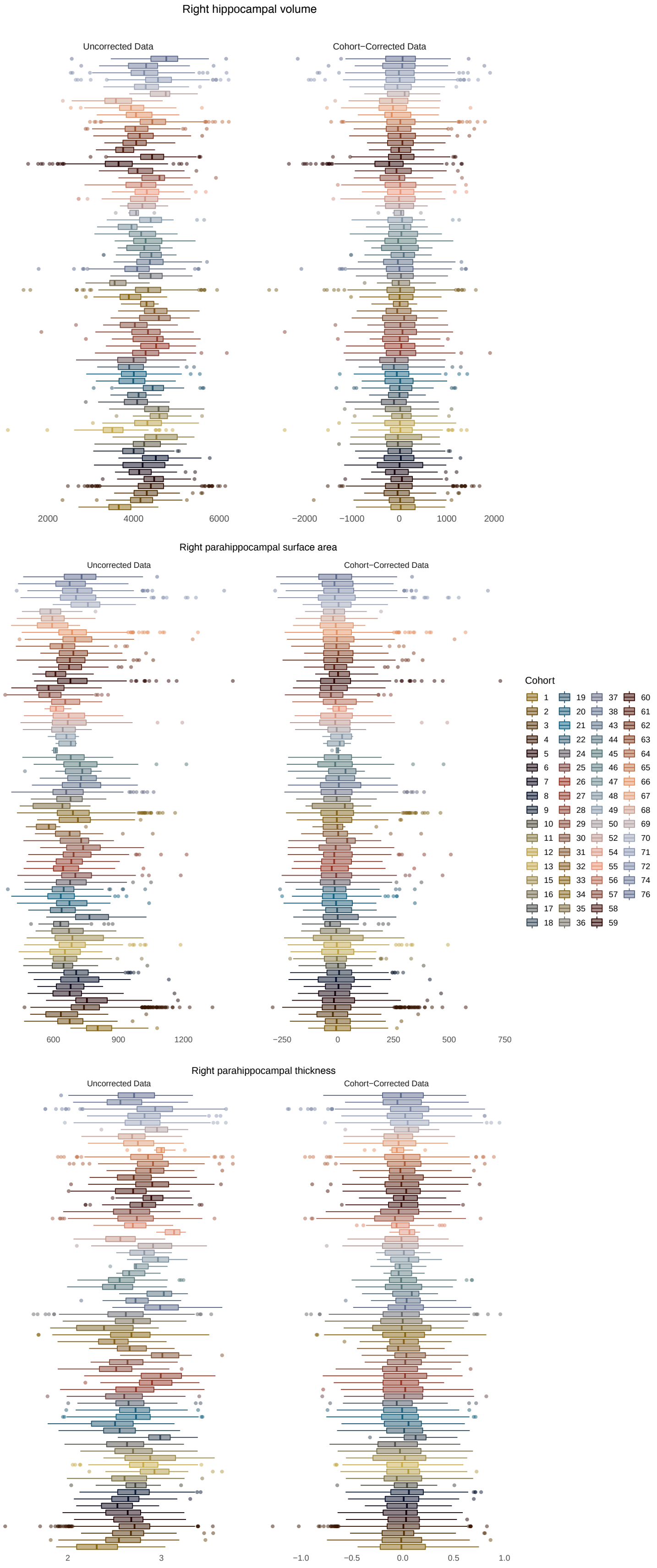


Supplemental Figure 1. Boxplot visualization of comparison of right hippocampal volume, parahippocampal surface area and thickness before and after adjustment. As age ranges differed for each cohort this was done in two steps: initially, a linear model was used to account for cohort effects and non-linear age effects, using a third degree polynomial function. Next, random forest regression modelling was used to additionally account for field strength and FreeSurfer version. In the left panel, volumes were not adjusted, this displays the raw data for each cohort. In the right panel, volumes were adjusted.
