## Supplementary figures and images for "Greater male than female variability in regional brain structure across the lifespan"

### Supplemental Figure 2

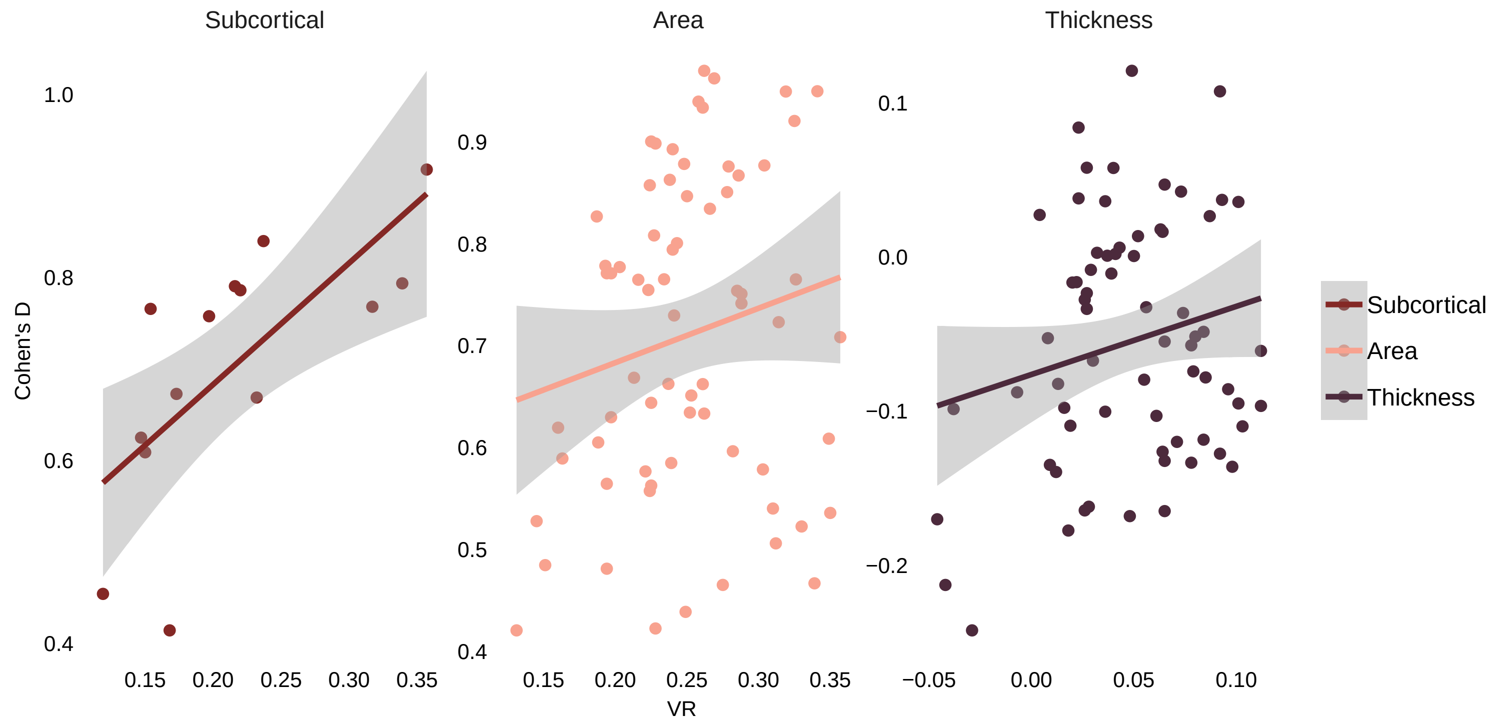
