## Supplemental Figure 3 for "Greater male than female variability in regional brain structure across the lifespan"

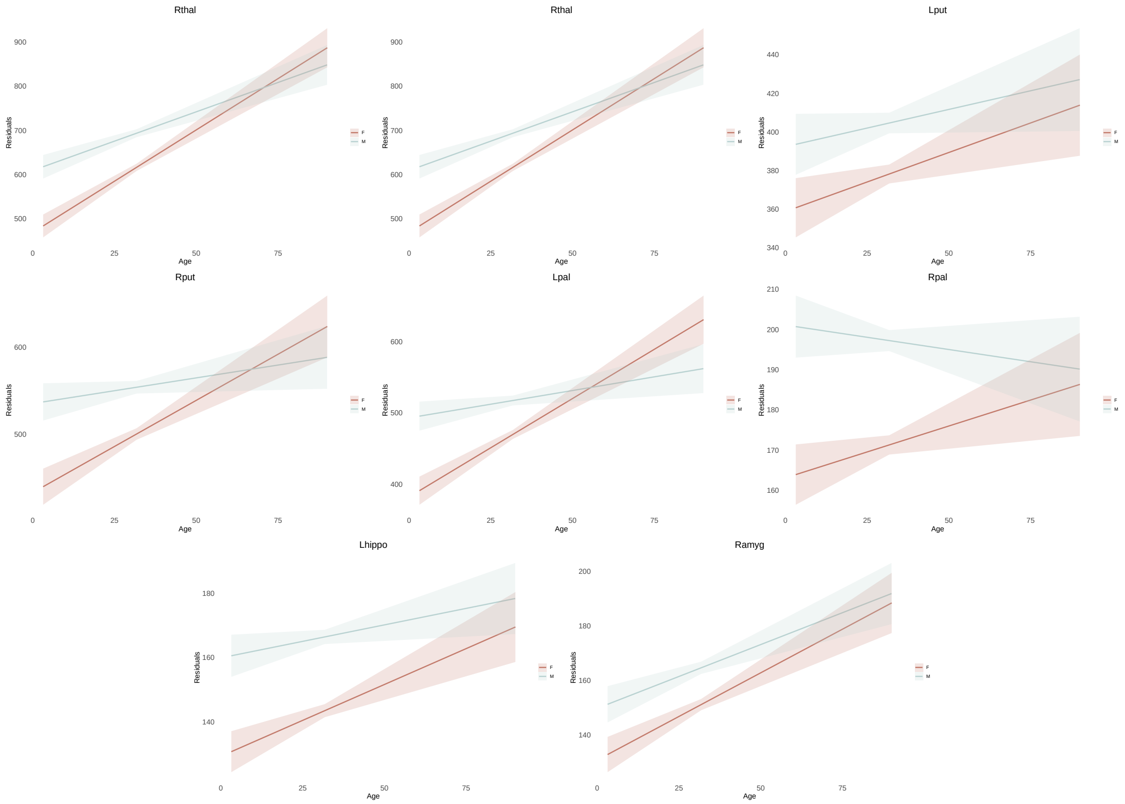


Supplemental Figure 3A. Sex differences in variability interacted with age in 50% of the subcortical volumes. Absolute residual values are modeled across the age range. Effects showed larger male than female variance in the younger age group, and a general trend of decreasing sex differences in variance with increasing age.

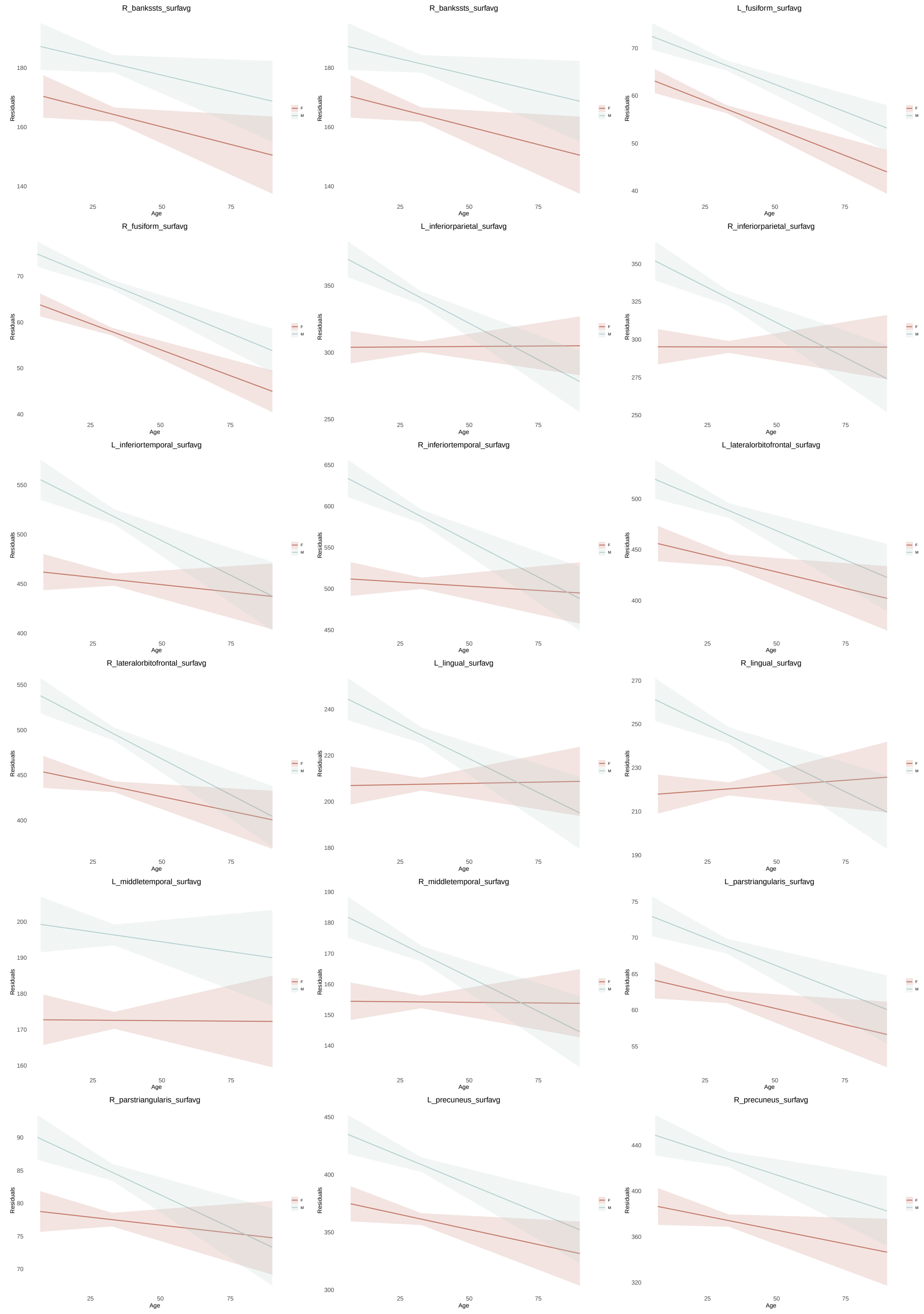


Supplemental Figure 3B-1. Sex differences in variability interacted with age in 30 % of cortical surface area measures. Absolute residual values are modeled across the age range. Effects showed larger male than female variance in the younger age group, and a general trend of decreasing sex differences in variance with increasing age.

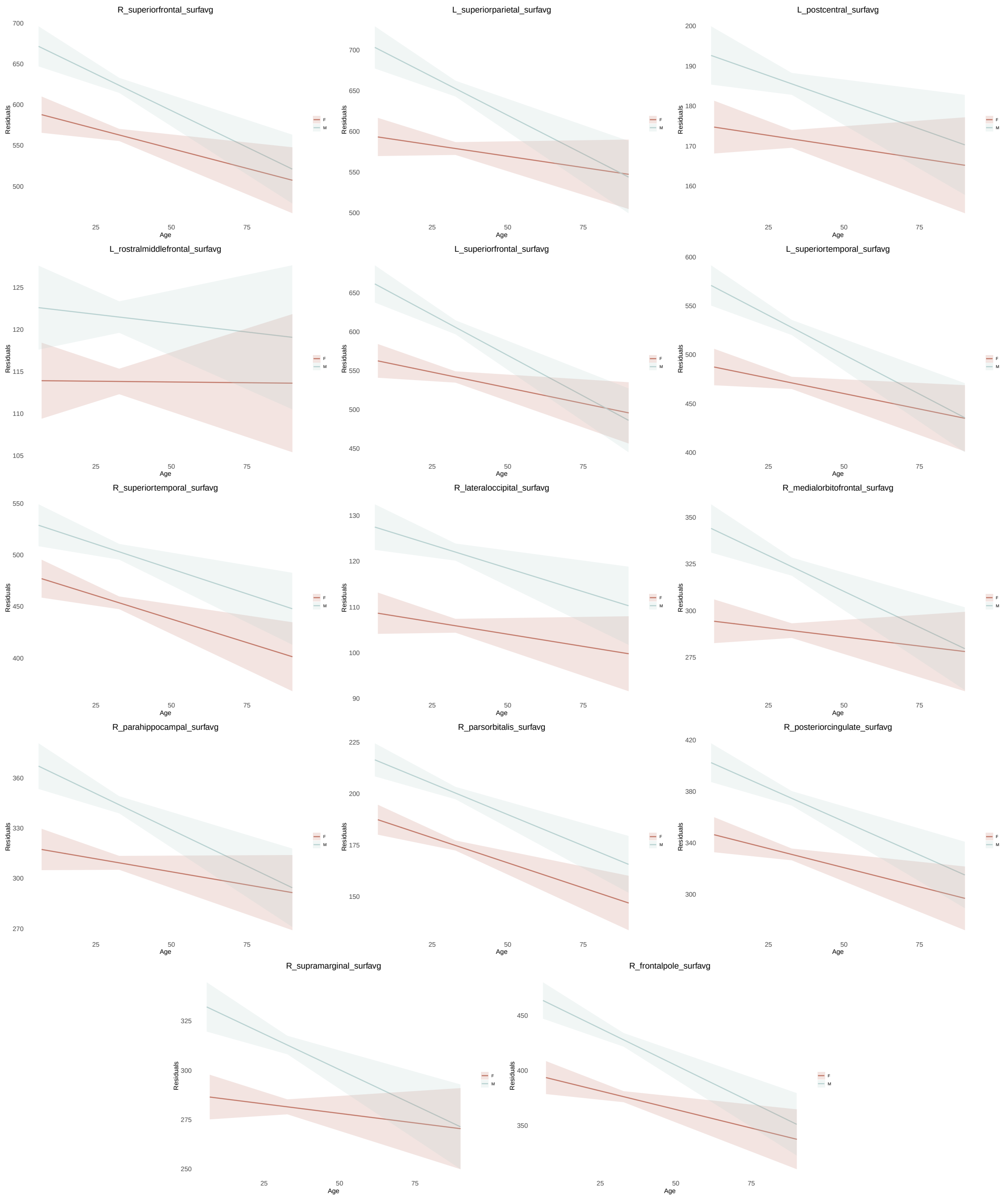


Supllemental Figure 3B-2. Sex differences in variablity interacted with age in 30% of cortical surface area measures. Absolute residual values are modeled across the age range. Effects showed larger male than female variance in the younger age group, and a general trend of decreasing sex differences in variance with increasing age.
