## Supplemental Table 1 for "Greater male than female variability in regional brain structure across the lifespan"

**Supplemental Table 1.** Screening Process and Eligibility Criteria, Scanner, Image Acquisition Parameters and Image Segmentation Software

| Sample | Screening Process | Eligibility Criteria | Magnet strength/ Scanner Vendor | Acquisition parameters | Freesurfer version |
|---|---|---|---|---|---|
| **ADHD-NF** | KSADS | No head trauma, no neurological and psychiatric history, no lifetime alcohol or substance abuse, no previous or current use of psychotropic medication, IQ>75. | 3T Siemens Tim Trio | T1-weighted 3D MPRAGE; TR/TE/TI/FA=2300ms/3030ms/900ms/9$^o$; Image matrix= 256 × 256; 192 sagittal slices; voxel size=1mm$^3$ | 5.3 |
| **AMC** | Personal Interview | No head trauma, no medical, neurological or psychiatric history, no lifetime alcohol or substance abuse, no previous or current use of psychotropic medication, IQ>75. No history of any psychiatric disorders in 1st degree family members. | 3T Philips Intera | T1-weighted 3D MPRAGE; TE range= 3.5-4.6ms, TR range= 9-9.663ms, FA= | 5.1 |
| **Basel** | Personal Interview | No head trauma, no medical, neurological, or psychiatric history, no alcohol or substance abuse ever, no previous or current use of psychotropic medication, IQ>75. No history of any psychiatric disorders in 1st degree family members. | 3T Siemens | T1-weighted MPRAGE; TR/TE/TI/FA= 2000 ms/3.37 ms/1000 ms/8$^o$; image matrix= 256×256x160; voxel size=1x1x1 mm$^3$; sagittal plane | 6.0 |
| **Betula** | Personal Interview | No head trauma, no medical, neurological or psychiatric history, no lifetime alcohol or substance abuse, no previous or current use of psychotropic medication. | 3T General Electric Discovery MR750 | T1-weighted MPRAGE; TR/TE/TI/FA=8.1240 ms/3.2000 ms/450ms/ 12°; image matrix = 256x256 | 5.3 |
| **Barcelona (1)** | KSADS | No head trauma, no medical, neurological or psychiatric history, no lifetime alcohol or substance abuse, no previous or current use of psychotropic medication, IQ>75. No history of any psychiatric disorders in 1st degree family members. | 1.5 T General Electric Signa | T1-weighted; image matrix = 256 x 256; 128 slices; voxel size 1 x 1 x 1 mm$^3$. | 5.3 |
| **Barcelona (2)** | KSADS | No head trauma, no medical, neurological or psychiatric history, no lifetime alcohol or substance abuse, no previous or current use of psychotropic medication, IQ>75. No history of any psychiatric disorders in 1st degree family members. | 3 T Siemens MAGNETOM TIM Trio | T1-weighted; image matrix = 256 x 256; 240 slices; voxel size 1 x 1 x 1 mm$^3$. | 5.3 |

| | | | | | |
|---|---|---|---|---|---|
| **BIG** | Questionnaire for psychiatric history | No head trauma, no medical, neurological or psychiatric history, no lifetime alcohol or substance abuse, no previous or current use of psychotropic medication, IQ>70. No history of any psychiatric disorders in 1st or 2nd degree family members. | 1,5 T Siemens Sonata and Avanto and 3 T Siemens Trio, TimTrio and Skyra | T1-weighted 3D MPRAGE; TR/TE/TI/sagittal slices = 1940-2730 ms/850-110 ms/2.92-4.58 ms; 176-192 sagittal slices; voxel size= 1.0x1.0x1.0 $mm^3$ | 5.3 |
| **Bonn** | Personal interview | No head trauma, no medical, neurological or psychiatric history, no previous or current use of psychotropic medication. | 3T Siemens Trio | TR/TE/FA= 1570-1660ms/2.75-3.42ms/8-9° | NA |
| **Bordeaux** | Personal interview | No head trauma, no previous or current use of psychotropic medication, IQ>75. | 3T Phillips Achieva | T1-weighted 3D TFE; TR/TE/TI/FA= 20 ms/4.6 ms/800 ms/10°; image matrix = 256x256x180$mm^3$; voxel size=1$mm^3$ | 5.3 |
| **BrainSCALE** | Personal interview | No head trauma, no medical, neurological or psychiatric history, no lifetime alcohol or substance abuse, no previous or current use of psychotropic medication. | 1.5T Philips Achieva | T1-weighted 3D SPGR; TR/TE/ FA= 30 ms/4.6 ms/30°; image matrix=256x256; 160–180 contiguous coronal slices; voxel size=1 x 1 x 1.2 $mm^3$ | 5.1 |
| **BRCATLAS** | Telephone interview | No head trauma, no medical, neurological or psychiatric history, no lifetime alcohol or substance abuse, no mild cognitive impairment, no previous or current use of psychotropic medication, IQ>75. | 3T General Electric Signa HDx | T1 weighted: TR/TE/TI/FA= 6.9 ms/2.8 ms/650 ms/8°; image matrix = 256x256x180$mm^3$; voxel size=1$mm^3$; sagittal plane acquisition | 5.3 |
| **CAMH** | SCID | No head trauma, no neurological or psychiatric history, no alcohol or substance abuse preceding 6 months, no previous or current use of psychotropic medication, IQ>75. No history of any psychotic disorders in 1st degree family members. | GE 1.5 T (echospeed) | 124 Axial inversion recovery–prepared spoiled gradient recall images, 1.5-mm-thick slice acquisition TE=5.3ms; TR=12.3ms; time to inversion, 300.0ms; flip angle, 20°. | 5.3 |
| **Cardiff** | MINI | No head trauma, no medical history, including neurological and psychiatric history, no alcohol or substance abuse in the preceding 6 months, no previous or current use of psychotropic medication. | 3T General Electric Signa | T1-weighted 3D FSPGR; TR/TE/TI/FA=7.9ms/3.0ms/450ms/20°; Image matrix= 256 × 192 x 172; voxel size=1$mm^3$ | 5.3 |
| **CEG** | Teacher and parent Conners' | No head trauma, no medical history, including neurological and psychiatric history, no alcohol or substance abuse in the preceding 6 months, no previous or current use of psychotropic medication. IQ>75 | 3T General Electric Signa | T1-weighted 3D SPGR; TR/TE/ FA=2000ms/30ms/90°; Image matrix= 128 × 128; 43 slices | 5.3 |

| | | | | | |
|---|---|---|---|---|---|
| **CIAM** | SCID | No head trauma or psychiatric history, no previous or current use of psychotropic medication, IQ>75. | 3T Siemens Allegra | T1-weighted 3D MPRAGE; TR/TE/TI/FA= 2530 ms/1.53, 3.21, 4.89, 6.57/2.91 ms/ ms/7$^{o}$; image matrix= 256x256; 128 sagittal slices; voxel size= 1.3x1.0x1.3 $mm^3$ | 5.3 |
| **CLiNG** | Personal Interview | No head trauma, no medical, neurological or psychiatric history, no lifetime alcohol or substance abuse, no previous or current use of psychotropic medication, IQ>75. No history of any psychiatric disorders in 1st degree family members. | 3T Siemens Tim Trio | T1-weighted 3D MPRAGE; TR/TE/TI/FA=2250 ms/3.26 ms/900 ms/9°; image matrix = 256 x 256; 192 sagittal slices; voxel size= 1 $mm^3$ | 5.3 |
| **CODE** | SCID Interview | No history of or current Axis-1 or 2 disorders, history of or current neurological disorder or brain injury, Serious medical condition, Severe cognitive impairment, Substance-related abuse or dependence disorder, Use of psychotropic medication, Use of central-acting medication, Pregnancy, General MRI contraindications. | 3T Siemens Trio (4 Sites), 3 T Philips Achieva (1 site) | Siemens: T1 mprage, voxel size 1 mm x 1 mm x 1 mm; TR=1900 msec; TE=2.52 msec; Sample 1: 192 slices, Sample 2: 170 slices<br>Philips: T1 3D-TFE, voxel size 1 mm x 1 mm x 1 mm; TR=8.3 msec; TE=3.8 msec; 170 slices. | 5.3 |
| **COMPULS/TS EUROTRAIN** | KSADS | No head trauma, no medical, neurological or psychiatric history, no alcohol or substance abuse preceding 6 months, no previous or current use of psychotropic medication, IQ>75. No history of any psychiatric disorders in 1$^{st}$ or 2$^{nd}$ degree family members. | 3T Siemens Tim Trio and Prisma | T1-weighted 3D MPRAGE; TR/TE/FA=2300 ms/2.98 ms/9°; image matrix = 256 x 256; 176 sagittal slices; voxel size= 1x1x1.2 $mm^3$ | 5.3 |
| **ENIGMA-HIV** | MINI | No head trauma, no medical, neurological or psychiatric history, no alcohol or substance abuse preceding 6 months, no previous or current use of psychotropic medication, IQ>75. No mild cognitive impairment | 3T Siemens Allegra | T1-weighted MPRAGE; TR/TE/TI/FA=2400 ms/2.38 ms/1000 ms/ 8°; 162 slices; voxel size= 1 $mm^3$ | 5.1 |
| **ENIGMA-OCD (1)** | MINI-Plus | No head trauma, no medical, neurological or psychiatric history, no lifetime alcohol or substance abuse, no previous or current use of psychotropic medication, IQ>75. No cognitive impairment | 3T Siemens Allegra | T1-weighted 3D MPRAGE; TR/TE/TI/FA=2300 ms/3.93 ms/ 1100 ms/12$^{o}$; image matrix =256×240; 160 contiguous sagittal slices; voxel size=1.3 x 1 x 1 $mm^3$ | 5.3 |

| | | | | | |
|---|---|---|---|---|---|
| **ENIGMA-OCD (2)** | SCID-I | No medical or psychiatric history | 1.5T Siemens Sonata | T1-weighted 3D MPRAGE; TR/TE/TI/FA=2700 ms/4 ms/ 950 ms/8$^{o}$; image matrix =256×192; 160 slices; voxel size= 1 $mm^3$ | 5.3 |
| **ENIGMA-OCD (3)** | SCID-I | No medical or psychiatric history | 3T General Electric Signa | T1-weighted 3D MPRAGE; image matrix =256×256; 172 slices; voxel size= 1x0.977x0.977 $mm^3$ | 5.3 |
| **ENIGMA-OCD (4)** | Personal Interview | No head trauma, no medical, neurological or psychiatric history, no lifetime alcohol or substance abuse, no previous or current use of psychotropic medication, IQ>75. | 3T Phillips Intera | T1-weighted 3D MPRAGE; TR/TE/ FA=9.69 ms/4.60 ms/8$^{o}$; image matrix =256×256; 182 slices; voxel size=1 x 1 x 1.2 $mm^3$ | 5.3 |
| **ENIGMA-OCD (5)** | SCID | No head trauma, no neurological or psychiatric history, no lifetime alcohol or substance abuse, no previous or current use of psychotropic medication, IQ>75. | 1.5T General Electric Signa | T1-weighted 3D SPGR; TR/TE/FA= 14.8 ms/ 1.7 ms/ 20º; image matrix= 256 x 256 x 124; voxel size: 0.94 x 0.94 x 1.50 mm | 5.3 |
| **ENIGMA-OCD (7)** | SCID-I/NP | No head trauma, no medical, neurological or psychiatric history, no alcohol or substance abuse in the preceding 6 months, no previous or current use of psychotropic medication, IQ>70. No history of any psychiatric disorders in 1st or 2$^{nd}$ degree family members | 1.5 T General Electric Signa | T1-weighted 3D FSPGR; TR/TE/ FA=11.8ms/4.2ms/90$^{o}$; Image matrix= 256 × 256 x 130; voxel size=1.2$mm^3$ | 5.3 |
| **FIDMAG** | Personal interview; structured interview in part of the sample | No head trauma, no medical, neurological or psychiatric history, no lifetime alcohol or substance abuse, no previous or current use of psychotropic medication, IQ>70. | 1.5 T General Electric Signa | T1-weighted MPRAGE; TR/TE/FA=2000 ms/4 ms/ 9$^{o}$; image matrix=512 x 512; 180 contiguous sagittal slices; voxel size=0.56 x 0.56 x 1 mm3 | 5.3 |
| **GSP** | Structured phone screen and study specific self-report battery and clinical | No head trauma, no medical, neurological or psychiatric history, no lifetime alcohol or substance abuse, no current use of psychotropic medication, normal brain anatomy following brain scan. | 3T Siemens Tim Trio | T1-weighted 3D multi-echo MPRAGE; TR/TE/TI/FA =2200 ms/1.54-7 ms/ 1100/7 $^{o}$; voxel size=1.2x1.2x1.2 mm | 4.5 |

| | | | | | |
|---|---|---|---|---|---|
| | screen | | | | |
| **HUBIN** | SCID-I | No head trauma, no medical, neurological or psychiatric history, no lifetime alcohol or substance abuse, no previous or current use of psychotropic medication, IQ>75. No history of any psychiatric disorders in 1st degree family members. | 1.5 T General Electric Signa | T1-weighted SPGR; TR/TE/FA= 24 ms/6 ms/35 $^{o}$; 124 coronal slices; voxel size 0.86 x 0.86 x 1.50 mm3. | 5.3 |
| **HMS** | Personal Interview | No head trauma, no medical, neurological or psychiatric history, no lifetime alcohol or substance abuse, no previous or current use of psychotropic medication, IQ>75. No history of any psychiatric disorders in 1st degree family members. | 1.5T Siemens Magnetom Sonata | T1-weighted 3D MPRAGE; TR/TE/TI/FA=1900 ms/4.0 ms/700 ms/15°; image matrix = 256 x 256; 176 consecutive sagittal slices; voxel size=1 $mm^3$ | 5.3 |
| **IDIVAL (1) + (2)** | CASH | No head trauma, no medical, neurological or psychiatric history, no lifetime alcohol or substance abuse, no previous or current use of psychotropic medication, IQ>75. No history of any psychiatric disorders in 1st degree family members. | 3T Siemens Alegra, Phillips Achieva 1.5T General Electric Signa | T1-weighted SPGR; TR/TE/FA=24 ms/5 ms/ 5$^{o}$; image matrix=256x192; T1-weighted SPGR; TR/TE/FA=3000 ms/3.9 ms/8$^{o}$; image matrix=256x256; voxel size=1$mm^3$; sagittal plane acquisition | 5.3 |
| **IDIVAL (3)** | Personal Interview | No lifetime history of Axis I psychiatric disorders, no mild cognitive | 3T Phillips Achieva | T1-weighted SPGR; TR/TE/FA=3000 ms/4.6 ms/8$^{o}$; image matrix=321x312; voxel size=1$mm^3$; sagittal plane acquisition | 5.3 |
| **IMAGEN** | DAWBA questionnaire clinician interview | No head trauma, no medical, neurological or psychiatric history, no previous or current use of psychotropic medication. IQ>75 | 3T Siemens Verio and TimTrio, Philips Achieva, General Electric Signa Excite, and Signa HDx | T1-weighted 3D MPRAGE; TR/TE/TI/FA=2300ms/3030ms/900ms/9$^{o}$; Image matrix= 256 × 256;192 sagittal slices; voxel size=1$mm^3$ | 5.3 |
| **IMH** | SCID-I/NP | No head trauma, no medical history, neurological or psychiatric history, no lifetime alcohol or substance abuse, as well as no previous or current use of psychotropic medication, IQ>75. No cognitive impairment | 3T Phillips Achieva | T1-weighted 3D MPRAGE; TR/TE/FA= 7.2ms/ 3.8ms/8$^{o}$; image matrix=256 x 256; 180 axial slices; voxel size=0.9$mm^3$ | 5.3 |
| **IMpACT** | SCID-I (and | No head trauma, no medical, neurological or | 1.5 T Siemens | T1-weighted 3D-MPRAGE; TR/TE/TI/FA | 5.3 |

| | | | | | |
|---|---|---|---|---|---|
| | SCID-II) | psychiatric history, no alcohol or substance abuse in the preceding 6 months, no previous or current use of psychotropic medication, IQ>70. No history of any psychiatric disorders in 1st or 2nd degree family members. | | =2730 ms/2.95 ms/1000 ms; 176 consecutive sagittal slices; voxel size= 1 $mm^3$ | |
| **Indiana (1)** | Personal Interview | No head trauma, no medical, neurological or psychiatric history, no lifetime alcohol or substance abuse, no previous or current use of psychotropic medication, IQ>75. | 1.5T General Electric Signa Horizon LX | T1-weighted 3D SPGR; TR/TE/FA=25 ms/3 ms/ 45$^o$; image matrix= 256 x 256; 124 contiguous coronal slices | 5.1 |
| **Indiana (2)** | Personal interview Structured phone screen | No head trauma, no medical, neurological or psychiatric history, no alcohol or substance abuse in the preceding 6 months, no previous or current use of psychotropic medication, IQ>75. | 3T Siemens Magnetom TrioTim | T1-weighted MPRAGE; TR/TE/FA=2300 ms/2.95 ms/ 9$^o$; image matrix=256 x 240; 176 contiguous sagittal slices | 5.1 |
| **Indiana (3)** | Personal interview Structured phone screen | No head trauma, no medical, neurological or psychiatric history, no alcohol or substance abuse in the preceding 6 months, no previous or current use of psychotropic medication, IQ>75. | 3T Siemens Skyra | T1-weighted MPRAGE GRAPPA2; TR/TE/FA=2300 ms/2.91 ms/ 9$^o$; image matrix=256 x 240; 160 contiguous sagittal slices | 5.1 |
| **Johns Hopkins** | Personal Interview | No head trauma, no medical, neurological or psychiatric history, no alcohol or substance abuse preceding 6 months, never prescribed with psychotropic medication, IQ>75. No mild cognitive impairment | 1.5T General Electric Signa | T1-weighted SPGR; TR/TE/FA=35 ms/5 ms/ 45$^o$; image matrix=256x256; 124 slices | 5.3 |
| **KaSP** | MINI | No head trauma, no medical, neurological or psychiatric history, no lifetime alcohol or substance abuse, no previous or current use of psychotropic medication, IQ>75. No history of any psychiatric disorders in 1st or 2nd degree family members. | 3T General Electric | T1-weighted SPGR; TR/TI/FA=7.904 ms/450 ms/12 $^o$; image matrix= 256 x 256 $mm^3$; 145 sagittal slices ; voxel size=0.934 x 0.934 x 1.2 mm3 | 5.3 |
| **Leiden** | Self-report | No psychiatric or neurological disorders, no use of psychotropic medications | 3T Philips Achieva | T1-weighted 3D SPGR; TR/TE = 9.76 ms/4.59 ms; image matrix=256x256; 160–180 contiguous coronal slices; voxel size=0.875x 0.875 x 1.2 $mm^3$ | 5.3 |
| **MCIC (1) + (2)** | SCID, SCID-I/NP, CASH | No head trauma, no medical, neurological or psychiatric history, no lifetime alcohol or | 1.5T Siemens Sonata-3T Siemens | T1-weighted MPRAGE sequence; TR/TE/TI/FA=2530 ms/4.76 ms/1100 | 5.3 |

| | | | | | |
|---|---|---|---|---|---|
| | | substance abuse, no previous or current use of psychotropic medication, IQ>75. | Trio | ms/20°; image matrix=256×256×128 cm; voxel size=0.625 $mm^3$ | |
| **Meth-CT** | SCID DSM-IV | No head trauma, no medical, neurological or psychiatric history, no lifetime alcohol or substance abuse, no previous or current use of psychotropic medication, IQ>70. | 3T Siemens Allegra | T1-weighted 3D MPRAGE; TR/graded TE/FA=2530 ms/ 1.53, 3.21, 4.89, 6.57 ms/ 7°; 160 contiguous sagittal slices; voxel size=1 x 1 *x 1 mm3 | 5.3 |
| **MHRC** | Personal interview | No head trauma, no medical history, including neurological and psychiatric history. No Family History of neurological or psychiatric disorders | 3T Philips Achieva | T1-weighted TFE; TR/TE/ FA=8.2ms/3.7ms/8°; voxel size=0.83 x 0.83 x 1 $mm^3$ | 5.3 |
| **Muenster** | SCID | No head trauma, no medical, neurological or psychiatric history, no lifetime alcohol or substance abuse, no previous or current use of psychotropic medication, IQ>75. | 3T Phillips Intera | T1 weighted TFE: TR/TE/FA= 7.4 ms/3.4 ms/9°; image matrix = 256x204x160$mm^3$; voxel size=0.5$mm^3$; sagittal plane acquisition | 5.3 |
| **NESDA** | CIDI | No lifetime history of Axis-I diagnoses, no lifetime medical or neurological morbidity including hypertension, no lifetime substance dependence, no substance abuse in the preceding year, no medication use. | 3T Philips Achieva SENSE-6 to 8 channel head coil | T1-weighted 3D MPRAGE; TR/TE/FA= 9 ms/3.5 ms/8°; image matrix=256x256; 170 sagittal slices; voxel size=1$mm^3$ | 5.0 |
| **NeuroIMAGE** | KSADS-PL | No head trauma, no mild cognitive impairment, neurological or psychiatric history, no previous or current use of psychotropic medication, IQ>75. No history of any psychiatric disorders in 1st and 2nd degree family members. | 1.5 T Siemens AVANTO (Donders Centre for Cognitive Neuroimaging) 1.5 T Siemens SONATA (VU University Amsterdam) | MPRAGE 176 sagittal slices, repetition time=2,730ms, echo time=2.95ms, voxel size=1.0x1.0x1.0mm, field of view=256 mm | 5.3 |
| **Neuroventure** | DAWBA and BSI | No head trauma, no medical, neurological or psychiatric history, no lifetime alcohol or substance abuse, no previous or current use of psychotropic medication, IQ>75. | 3T SIEMENS TrioTim | T1-weighted 3D MPRAGE; TR/TE/ FA= 2300 ms/2.96 ms/9°; image matrix= 256x256; voxel size= 1.0x1.0x1.0 $mm^3$ | 5.3 |
| **NTR (1)** | MINI, BDI, STAI, STAS, YBOCS | No head trauma, no previous or current use of psychotropic medication, normal IQ. | 3T Philips Intera | T1-weighted 3D MPRAGE; TR/TE/FA=9.64 ms/4.60 ms/8 °; image matrix=256 x 256; 182 coronal slices; voxel size=1 x1x1.2 $mm^3$ | 5.1 |
| **NTR (2)** | CIDI, MADRS, BDI, | No current psychiatric disorder, no current use of psychotropic medication, normal IQ. | 1.5 T Siemens Sonata | T1-weighted 3D MPRAGE; TR/TE/TI/FA= 15 ms/7 ms/300 ms/8°; image | 5.1 |

| | | | | | |
|---|---|---|---|---|---|
| | STAI | | | matrix=256x176; 160 coronal slices; voxel size=1x1x1.5 $mm^3$ | |
| **NTR (3)** | DISC-IV | No head trauma, no medical, neurological or psychiatric history, no lifetime alcohol or substance abuse, no mild cognitive impairment, no previous or current use of psychotropic medication, IQ>75. | 1.5T Siemens Sonata | T1-weighted 3D MPRAGE; TR/TE/TI/FA=1900 ms/3.93 ms/1100 ms/ 15°; image matrix=256 x 224; 160 sagittal slices; voxel size=1 $mm^3$ | 5.1 |
| **NU** | SCID | No head trauma, no medical, neurological or psychiatric history, no lifetime alcohol or substance abuse, no previous or current use of psychotropic medication, IQ>75. No history of any psychiatric disorders in 1st degree family members. | 1.5T SIEMENS Vision | T1-weighted 3D MPRAGE; TR/TE/TI/FA=2200 ms/4.13 ms/766 ms/13°; voxel size =0.8$mm^3$; axial plane acquisition. | 5.3 |
| **NUIG** | SCID | No history of head trauma or injury; No family history of ANY psychiatric disorders in 1st degree family members; IQ>75 | 1.5T Siemens Magnetom Symphony | T1-weighted 3D MPRAGE; TR/TE=1140/4.38 mm Image matrix = 256x256; FOV=230mm; voxel size: 0.45x0.45x0.9mm | 5.1 |
| **NYU** | SCID-NP for DSM-IV | No head trauma, no medical history, including neurological and psychiatric history, no lifetime alcohol or substance abuse, no previous or current use of psychotropic medication. IQ>75. | 3T Siemens Allegra | T1-weighted 3D MPRAGE; TR/TE/TI/FA=2530ms/3.25ms/1100ms/7$^o$ | 5.3 |
| **OATS (1-4)** | Personal interview | No head trauma, no current diagnosis of a psychotic disorder, no neurological disorder, no malignancy (other than skin cancer) or other severe medical comorbidity, no mild cognitive impairment, IQ>75. | 1.5T Philips Gyroscan, Siemens Magnetom Avanto, Siemens Sonata; 3T Philips Achieva Quasar Dual, a | T1-weighted 3D acquisition; TR/TE/TI/FA=15370 ms/3.24 ms/780 ms/8$^o$; 144 slices; voxel size=1 x 1 x 1.5 $mm^3$ | 5.3 |
| **OLIN** | SCID I | No head trauma, no medical, neurological or psychiatric history, no alcohol or substance abuse preceding 6 months, never prescribed with psychotropic medication, IQ>75. | 3T Siemens Alegra | T1-weighted 3D MPRAGE; TR/TE/TI/FA= 2300 ms/2.91 ms/900 ms/9$^o$; image matrix= 256x240x192; 160 sagittal slices; voxel size= 1.0x1.0x1.2 $mm^3$ | 5.1 |
| **QTIM** | CIDI | No head trauma, no medical history, neurological and psychiatric history, no alcohol or substance abuse in the preceding 6 months, no antidepressant medication or medication affecting cognition. | 4T Bruker | T1-weighted 3D MPRAGE: TR/TE/TI/FA = 1500 ms/3.35 ms/ 700 ms/ 8°; image matrix= 256 × 256 × 256 or 256 × 256 × 240; 256 coronal slices; voxel size= 0.9 $mm^3$ | 5.1 |

| | | | | | |
|---|---|---|---|---|---|
| **Oxford** | KSADS | No head trauma, no medical, neurological or psychiatric history, no lifetime alcohol or substance abuse, no previous or current use of psychotropic medication, IQ>75. | 1.5T Siemens Sonata | T1-weighted 3D MPRAGE; TR/TE =12 ms/5.6 ms; image matrix =256×240x 208 $mm^3$; voxel size=1 $mm^3$ | 5.3 |
| **Sao Paulo-1** | SCID | No head trauma, neurological or psychiatric history, no lifetime alcohol or substance abuse. | 1.5T Siemens Espree | T1-weighted 3D MPRAGE; TR/TE/TI/FA=2400 ms/3.65 ms/ 0 ms/8$^o$; 160 contiguous sagittal slices; voxel size=1.3 x 1.3x 1.2 mm3 | 5.3 |
| **Sao Paulo-3** | SCID | No head trauma, neurological or psychiatric history, no lifetime alcohol or substance abuse. IQ>75 | 1.5T General Electric Signa | T1-weighted FSPGR ; TR/TE/TI/FA=21.7 ms/52 ms /20$^o$; 124 axial slices; voxel size= 0.86 x 0.86 x 1.5 mm3 | 5.3 |
| **SHIP-2** | Personal Interview | No head trauma, no neurological and psychiatric history, no alcohol or substance abuse in the preceding 6 months, no previous or current use of psychotropic medication. IQ>75. | 1.5T Siemens Avanto | T1-weighted 3D MPRAGE; TR/TE/ FA=1900ms/3.4ms/15$^o$; voxel size=1$mm^3$ | 5.3 |
| **SHIP-TREND** | Personal Interview | No head trauma, no neurological and psychiatric history, no alcohol or substance abuse in the preceding 6 months, no previous or current use of psychotropic medication. IQ>75. | 1.5T Siemens Avanto | T1-weighted 3D MPRAGE; TR/TE/ FA=1900ms/3.4ms/15$^o$; voxel size=1 $mm^3$ | 5.3 |
| **Stages-Dep** | SCID-I | No head trauma, no medical, neurological or psychiatric history, no lifetime alcohol or substance abuse, no previous or current use of psychotropic medication, IQ>75. No history of any psychiatric disorders in 1st degree family members. | 3T Phillips Achieva | T1-weighted 3D-MPRAGE; TR/TE/TI/FA =6.7 ms/3.2 ms/200 ms/88$^o$; °; image matrix = 288 x 288; 170 consecutive sagittal slices; voxel size= 0.896×0.896×1.2 $mm^3$ | 5.1 |
| **Stanford** | SCID | No head trauma, no medical, neurological or psychiatric history, no lifetime alcohol or substance abuse, no previous or current use of psychotropic medication. no mild cognitive impairment. | 1.5T General Electric Signa Excite | T1-weighted SPGR; TR/TE/TI/FA=8.3-10.3 ms/1.7-3.0 ms/300 ms/15$^o$; image matrix= 256 x 192; 176 contiguous sagittal slices; voxel size=0.86x0.86x1.5 $mm^3$; sagittal plan acquisition | 5.3 |
| **StrokeMRI** | Personal | No head trauma, no medical, neurological or | 3T General Electric | T1-weighted FSPGR (BRAVO); | 5.3 |

| | | | | | |
|---|---|---|---|---|---|
| | interview | psychiatric history, no lifetime alcohol or substance abuse, no previous or current use of psychotropic medication, IQ>75. | 750 Discovery MRI scanner | TR/TE/TI/FA=8.16 s/3.18 ms/450 ms/12°; 188 sagittal slices; voxel size= 1.0x1.0x1.2 mm; FOV: 256 x 256 mm; scan time: 4:43 min | |
| **SYDNEY** | SCID | No head trauma, no medical history, neurological or psychiatric history, no alcohol or substance abuse preceding 6 months, as well as no previous or current use of psychotropic medication, IQ>75. | 3T General Electric Discovery MR750 | T1-weighted 3D MPRAGE; TR/TE/FA= 7264ms/ 2784ms/15$^{o}$; image matrix =256 x 256 x 196; voxel size=0.9mm$^{3}$ | 5.1 |
| **Sydney MAS** | Personal interview | No head trauma, no diagnosis of dementia, schizophrenia, bipolar disorder no psychotic symptoms, no neurological disorder, no mild cognitive impairment, IQ>75. | 3T Philips Achieva Quasar Dual | TR/TE = 6.39 ms/2.9 ms; 190 coronal slices; voxel size = 1mm$^{3}$ | 5.3 |
| **TOP** | PRIME-MD | No head trauma, no organic or other psychotic disorder (ICD codes 290-299), no substance abuse in the preceding 6 months, no previous or current use of psychotropic medication, IQ>75. No history of any psychiatric disorders in 1st degree family members. | 1.5T Siemens Magnetom Sonata | T1-weighted SPGR; TR/TE/TI/FA=2730 ms/3.93 ms/1000 ms/71$^{o}$; voxel size = 1.33x0.94x1mm$^{3}$; sagittal plane acquisition | 5.3 |
| **Tuebingen** | SCID I and II | No head trauma, no medical history, neurological or psychiatric history, no lifetime alcohol or substance abuse as well as no previous or current use of psychotropic medication, IQ>75. No history of any psychiatric disorders in 1st degree family members. | 1.5T Siemens Avanto | T1-weighted 3D MPRAGE; TR/TE/FA= 2250ms/ 3.93ms/8$^{o}$; image matrix =256 x 256; voxel size=1mm$^{3}$ | 5.3 |
| **UMCU** | CASH | No head trauma, no medical, neurological or psychiatric history, no lifetime alcohol or substance abuse, no previous or current use of psychotropic medication, IQ>75. No history of any psychiatric disorders in 1st degree family members. | 3T Philips Achieva | T1-weighted 3D TFE; TE/TR/FA= 4.50 ms/9.96 ms/ 8˚; FOV=224mm; image matrix = 256x256 mm; voxel size=.875x.875x1 mm$^{3}$ | 5.3 |
| **UNIBA** | SCID-NP | No head trauma, no medical, neurological or psychiatric history, no lifetime alcohol or substance abuse, no previous or current use of psychotropic medication, IQ>75. No | 3T General Electric | T1-weighted 3D SPGR; TE/FA = min full/ 6°; image matrix= 256×256 x124 | 5.3 |

| | | | | | |
|---|---|---|---|---|---|
| | | history of any psychiatric disorders in 1st degree family members. | | | |
| **UPENN** | SCID | No head trauma, no medical history, including neurological and psychiatric history, no alcohol or substance abuse preceding 6 months, no previous or current use of psychotropic medication, IQ>75. No history of any psychiatric disorders in 1st degree family members. | 3T Siemens Tim Trio | T1-weighted 3D MPRAGE; TR/TE/TI/FA=1810 ms/3.51 ms/1100 ms/9$^{o}$; image matrix= 256 × 192;160 axial slices | 5.3 |
| **Yale** | KSADS-PL | No head trauma, neurological or psychiatric history, no alcohol or substance abuse in the preceding 6 months, no previous or current use of psychotropic medication, IQ>75. | 3T General Electric Signa | T1-weighted 3D MPRAGE; image matrix =256×256; voxel size=0.976 x 0.976 x 1 mm3 | 5.3 |

Abbreviations of Terms: BDI = Behavioural Descriptive Interview; BSI = Brief Symptom Inventory; CASH = Comprehensive assessment of symptoms and history; CDR = Clinical Dementia Rating; CIDI = Composite International Diagnostic Interview; DAWBA = Development and Well-Being Assessment; DISC-IV = Diagnostic Interview Schedule for Children; DSM = Diagnostic and Statistical Manual of Mental Disorders (DSM); FA=flip angle; FSPGR=fast spoiled gradient echo sequence; GRE=spoiled gradient echo sequence; ICD= International Classification of Diseases; IR= inversion recovery; KSADS-PL= Kiddie Schedule for Affective Disorders and Schizophrenia-Present and Lifetime; MADRS = Montgomery-Asberg Depression Rating Scale; MINI = Mini International Neuropsychiatric Interview; MMSE = Mini Mental State Exam; PRIME-MD = Primary Care Evaluation of Mental Disorder; SCID = Structured Clinical Interview for DSM Disorders; SCID-I/NP = SCID Non-Patient version; SPGR=spoiled gradient recalled sequence; STAI = State-Trait Anxiety Inventory; STAS = State-trait anger scale; TE=echo time; TI=inversion time; TR=repetition time; TFE=turbo field echo sequence; YBOCS = Yale-Brown Obsessive Compulsive Scale

Abbreviations of studies: **ADHD-NF = Attention Deficit Hyperactivity Disorder- Neurofeedback Study; AMC = Amsterdam Medisch Centrum; Basel = University of Basel**; **Betula = Swedish longitudinal study on aging, memory, and dementia**; **BIG = Brain Imaging Genetics; Bonn = University of Bonn; Bordeaux = University of Bordeaux; BrainSCALE=Brain Structure and Cognition: an Adolescence Longitudinal twin study into Etiology; BRCATLAS= NIHR Biomedical Research Centre/ Mapping the relationship between the white matter and executive function across the adult lifespan; CAMH = Centre for Addiction and Mental Health; Cardiff = Cardiff University; CEG = Cognitive-experimental and Genetic study of ADHD and Control Sibling Pairs; CIAM = Cortical Inhibition and Attentional Modulation study; CLiNG = Clinical Neuroscience Göttingen**; **CODE = Chronic Depression Study; ENIGMA-HIV = Enhancing NeuroImaging Genetics through Meta-Analysis-Human Immunodeficiency Virus Working Group; ENIGMA-OCD = Enhancing NeuroImaging Genetics through Meta-Analysis- Obsessive Compulsive Disorder Working Group; FBIRN = Function Biomedical Informatics Research Network; EDINBURGH = The University of Edinburgh**; **GSP = Brain Genomics Superstruct Project (http://neuroinformatics.harvard.edu/gsp/; Holmes et al., Sci Data 2015; http://www.nature.com/articles/sdata201531); HMS = Homburg Multidiagnosis Study**; **ENIGMA-OCD (IDIBELL)**; **FIDMAG = Fundación para la Investigación y Docencia Maria Angustias Giménez**; **HUBIN = Human Brain Informatics; IDIVAL = Valdecilla Biomedical Research Institute; IMAGEN = the IMAGEN Consortium; IMH=Institute of Mental Health, Singapore; IMpACT = The International Multicentre persistent ADHD Genetics Collaboration; Indiana = Indiana University School of Medicine; KaSP= The Karolinska Schizophrenia Project; Leiden = Leiden University; MCIC = MIND Clinical Imaging Consortium formed by the Mental Illness and Neuroscience Discovery (MIND) Institute now the Mind Research Network (MRN; http//ww.mrn.org); Melbourne = University of Melbourne**; **Meth-CT = methamphetamine use; University of Cape Town; MHRC = Mental Health Research Center; Muenster = Muenster University; NESDA = The Netherlands Study of Depression and Anxiety**; Neuroventure = ; **NTR = Netherlands Twin Register**; **NU = Northwestern University; NYU = New York University**; **OLIN = Olin Neuropsychiatric Research Center; NUIG = National University of Ireland Galway**; **OATS = Older Australian Twins Study**; **Oxford =Oxford University; QTIM = Queensland Twin Imaging; Sao Paulo = University of Sao Paulo; SHIP = Study of Health in Pomerania; Stages-Dep= Stages of Depression Study; Stanford = Stanford University**; **StrokeMRI = Stroke Magnetic Resonance Imaging; SYDNEY = University of Sydney; Sydney MAS = Sydney Memory and Ageing Study; TOP = Tematisk Område Psykoser (Thematically Organized Psychosis Research); Tuebingen = University of Tuebingen; UMCU = Universitair Medisch Centrum Utrecht**; **UNIBA = University of Bari Aldo Moro**; **UPENN=University of Pennsylvania; Yale = Yale University**
