## Supplemental Table 2 for "Greater male than female variability in regional brain structure across the lifespan"

### Supplementary Table 2A

| Subcortical volume | Female (n=7141) | Male (n=6555) | Mean difference test | | Variance Ratio test | |
|---|---|---|---|---|---|---|
| | M | M | P | Cohen's D | VR | P |
| left thal | -174.193 | 189.936 | ** | 0.504 | 0.233 | ** |
| right thal | -175.412 | 191.140 | ** | 0.576 | 0.364 | ** |
| left caud | -65.407 | 71.566 | ** | 0.311 | 0.163 | ** |
| right caud | -70.631 | 78.224 | ** | 0.328 | 0.133 | ** |
| left put | -133.575 | 144.498 | ** | 0.453 | 0.205 | ** |
| right put | -132.828 | 144.790 | ** | 0.483 | 0.218 | ** |
| left pal | -54.597 | 59.591 | ** | 0.510 | 0.321 | ** |
| right pal | -44.439 | 48.627 | ** | 0.504 | 0.357 | ** |
| left hippo | -73.545 | 79.192 | ** | 0.387 | 0.119 | ** |
| right hippo | -67.811 | 74.070 | ** | 0.368 | 0.221 | ** |
| left amyg | -45.046 | 48.757 | ** | 0.492 | 0.156 | ** |
| right amyg | -49.629 | 53.719 | ** | 0.515 | 0.216 | ** |
| left accumb | -10.863 | 11.819 | ** | 0.208 | 0.162 | ** |
| right accumb | -11.997 | 13.134 | ** | 0.248 | 0.133 | ** |

### Supplementary Table 2B

| Surface area | Female (n=6243) | Male (n=5092) | Mean difference test | | Variance Ratio test | |
|---|---|---|---|---|---|---|
| | M | M | P | Cohen's D | VR | P |
| left bankssts | -21.428 | 26.514 | ** | 0.303 | 0.288 | ** |
| left caudalanteriorcingulate | -7.761 | 9.655 | ** | 0.138 | 0.144 | ** |
| left caudalmiddlefrontal | -32.171 | 40.016 | ** | 0.218 | 0.174 | ** |
| left cuneus | -23.913 | 28.961 | ** | 0.288 | 0.213 | ** |
| left entorhinal | -9.573 | 11.775 | ** | 0.282 | 0.318 | ** |
| left fusiform | -59.732 | 73.637 | ** | 0.401 | 0.218 | ** |
| left inferiorparietal | -89.239 | 110.767 | ** | 0.388 | 0.290 | ** |
| left inferiortemporal | -69.130 | 84.488 | ** | 0.403 | 0.198 | ** |
| left isthmuscingulate | -26.185 | 32.319 | ** | 0.424 | 0.309 | ** |
| left lateraloccipital | -116.996 | 144.255 | ** | 0.534 | 0.241 | ** |
| left lateralorbitofrontal | -33.519 | 41.514 | ** | 0.348 | 0.170 | ** |
| left lingual | -44.627 | 54.474 | ** | 0.276 | 0.210 | ** |
| left medialorbitofrontal | -32.826 | 40.488 | ** | 0.373 | 0.276 | ** |
| left middletemporal | -55.369 | 68.345 | ** | 0.396 | 0.261 | ** |
| left parahippocampal | -10.153 | 12.606 | ** | 0.234 | 0.324 | ** |
| left paracentral | -19.674 | 24.133 | ** | 0.270 | 0.294 | ** |
| left parsopercularis | -23.321 | 28.224 | ** | 0.214 | 0.365 | ** |
| left parsorbitalis | -12.910 | 15.872 | ** | 0.401 | 0.195 | ** |
| left parstriangularis | -26.699 | 32.436 | ** | 0.326 | 0.267 | ** |
| left pericalcarine | -18.170 | 22.274 | ** | 0.198 | 0.139 | ** |
| left postcentral | -75.090 | 92.913 | ** | 0.462 | 0.314 | ** |
| left posteriorcingulate | -21.743 | 26.882 | ** | 0.317 | 0.254 | ** |
| left precentral | -97.158 | 120.713 | ** | 0.551 | 0.316 | ** |
| left precuneus | -67.764 | 83.523 | ** | 0.437 | 0.283 | ** |
| left rostralanteriorcingulate | -14.519 | 17.886 | ** | 0.256 | 0.169 | ** |
| left rostralmiddlefrontal | -136.648 | 169.107 | ** | 0.549 | 0.293 | ** |
| left superiorfrontal | -141.866 | 175.228 | ** | 0.559 | 0.228 | ** |
| left superiorparietal | -79.266 | 98.252 | ** | 0.344 | 0.227 | ** |
| left superiortemporal | -85.038 | 104.778 | ** | 0.592 | 0.208 | ** |
| left supramarginal | -94.686 | 117.258 | ** | 0.487 | 0.304 | ** |
| left frontalpole | -3.241 | 3.968 | ** | 0.221 | 0.231 | ** |
| left temporalpole | -8.189 | 10.093 | ** | 0.316 | 0.219 | ** |
| left transversetemporal | -8.304 | 10.195 | ** | 0.268 | 0.217 | ** |
| left insula | -39.534 | 48.789 | ** | 0.474 | 0.229 | ** |
| right bankssts | -19.472 | 24.007 | ** | 0.337 | 0.269 | ** |
| right caudalanteriorcingulate | -12.282 | 15.246 | ** | 0.193 | 0.283 | ** |
| right caudalmiddlefrontal | -31.934 | 39.450 | ** | 0.213 | 0.229 | ** |
| right cuneus | -26.831 | 32.861 | ** | 0.330 | 0.236 | ** |
| right entorhinal | -9.061 | 11.120 | ** | 0.260 | 0.331 | ** |
| right fusiform | -75.126 | 91.735 | ** | 0.525 | 0.197 | ** |
| right inferiorparietal | -135.284 | 166.645 | ** | 0.540 | 0.306 | ** |
| right inferiortemporal | -64.768 | 79.764 | ** | 0.413 | 0.188 | ** |
| right isthmuscingulate | -21.334 | 26.370 | ** | 0.375 | 0.321 | ** |
| right lateraloccipital | -114.231 | 139.966 | ** | 0.515 | 0.267 | ** |
| right lateralorbitofrontal | -36.896 | 46.038 | ** | 0.349 | 0.221 | ** |
| right lingual | -44.954 | 55.584 | ** | 0.300 | 0.245 | ** |
| right medialorbitofrontal | -30.524 | 37.532 | ** | 0.404 | 0.223 | ** |
| right middletemporal | -66.294 | 81.941 | ** | 0.463 | 0.234 | ** |
| right parahippocampal | -14.183 | 17.365 | ** | 0.367 | 0.319 | ** |
| right paracentral | -22.852 | 28.213 | ** | 0.271 | 0.332 | ** |
| right parsopercularis | -20.480 | 25.363 | ** | 0.211 | 0.323 | ** |
| right parsorbitalis | -16.572 | 20.550 | ** | 0.414 | 0.178 | ** |
| right parstriangularis | -34.599 | 42.647 | ** | 0.341 | 0.270 | ** |
| right pericalcarine | -19.021 | 23.521 | ** | 0.198 | 0.151 | ** |
| right postcentral | -70.424 | 87.713 | ** | 0.443 | 0.292 | ** |
| right posteriorcingulate | -20.435 | 24.929 | ** | 0.295 | 0.240 | ** |
| right precentral | -110.167 | 136.382 | ** | 0.587 | 0.339 | ** |
| right precuneus | -84.731 | 104.341 | ** | 0.501 | 0.255 | ** |
| right rostralanteriorcingulate | -13.300 | 16.469 | ** | 0.257 | 0.203 | ** |
| right rostralmiddlefrontal | -124.963 | 153.957 | ** | 0.488 | 0.225 | ** |
| right superiorfrontal | -141.040 | 173.327 | ** | 0.543 | 0.270 | ** |
| right superiorparietal | -85.848 | 104.679 | ** | 0.382 | 0.212 | ** |
| right superiortemporal | -52.348 | 64.673 | ** | 0.408 | 0.231 | ** |
| right supramarginal | -68.219 | 83.861 | ** | 0.362 | 0.287 | ** |
| right frontalpole | -4.746 | 5.920 | ** | 0.247 | 0.186 | ** |
| right temporalpole | -5.293 | 6.502 | ** | 0.208 | 0.240 | ** |
| right transversetemporal | -5.637 | 6.915 | ** | 0.245 | 0.178 | ** |
| right insula | -48.677 | 60.048 | ** | 0.513 | 0.224 | ** |

### Supplementary Table 2C

| Thickness | Female (n=6620) | Male (n=5913) | Mean difference test | | Variance Ratio test | |
|---|---|---|---|---|---|---|
| | M | M | P | Cohen's D | VR | P |
| left bankssts | -0.001 | 0.002 | n.s. | 0.014 | 0.023 | ** |
| left caudalanteriorcingulate | 0.024 | -0.026 | ** | 0.203 | -0.035 | n.s. |
| left caudalmiddlefrontal | 0.005 | -0.006 | ** | 0.084 | 0.062 | n.s. |
| left cuneus | -0.001 | 0.002 | n.s. | 0.028 | 0.081 | * |
| left entorhinal | -0.016 | 0.018 | ** | 0.100 | 0.011 | n.s. |
| left fusiform | -0.001 | 0.001 | n.s. | 0.015 | 0.016 | n.s. |
| left inferiorparietal | 0.006 | -0.006 | ** | 0.115 | 0.126 | ** |
| left inferiortemporal | -0.004 | 0.005 | ** | 0.067 | -0.040 | n.s. |
| left isthmuscingulate | 0.007 | -0.007 | ** | 0.072 | -0.028 | ** |
| left lateraloccipital | 0.002 | -0.002 | * | 0.044 | 0.111 | ** |
| left lateralorbitofrontal | -0.004 | 0.006 | ** | 0.071 | 0.082 | ** |
| left lingual | -0.005 | 0.006 | ** | 0.090 | 0.051 | n.s. |
| left medialorbitofrontal | -0.006 | 0.008 | ** | 0.087 | 0.000 | n.s. |
| left middletemporal | -0.005 | 0.007 | ** | 0.087 | 0.045 | n.s. |
| left parahippocampal | 0.013 | -0.013 | ** | 0.085 | 0.011 | n.s. |
| left paracentral | 0.004 | -0.003 | * | 0.048 | 0.041 | ** |
| left parsopercularis | -0.004 | 0.005 | ** | 0.066 | 0.063 | ** |
| left parsorbitalis | 0.010 | -0.011 | ** | 0.102 | 0.037 | ** |
| left parstriangularis | 0.002 | -0.002 | n.s. | 0.025 | 0.027 | ** |
| left pericalcarine | -0.001 | 0.002 | n.s. | 0.026 | 0.068 | ** |
| left postcentral | 0.006 | -0.007 | ** | 0.122 | 0.055 | ** |
| left posteriorcingulate | 0.003 | -0.003 | n.s. | 0.037 | 0.063 | ** |
| left precentral | 0.004 | -0.004 | ** | 0.066 | 0.077 | ** |
| left precuneus | -0.002 | 0.003 | n.s. | 0.040 | 0.058 | ** |
| left rostralanteriorcingulate | 0.018 | -0.020 | ** | 0.166 | -0.066 | n.s. |
| left rostralmiddlefrontal | 0.002 | -0.002 | n.s. | 0.031 | 0.073 | ** |
| left superiorfrontal | 0.010 | -0.011 | ** | 0.174 | 0.037 | n.s. |
| left superiorparietal | 0.006 | -0.006 | ** | 0.119 | 0.138 | ** |
| left superiortemporal | -0.004 | 0.005 | ** | 0.062 | 0.042 | ** |
| left supramarginal | 0.007 | -0.007 | ** | 0.119 | 0.067 | ** |
| left frontalpole | 0.013 | -0.014 | ** | 0.091 | 0.024 | n.s. |
| left temporalpole | 0.000 | -0.001 | n.s. | 0.003 | 0.011 | n.s. |
| left transversetemporal | 0.017 | -0.018 | ** | 0.169 | 0.011 | n.s. |
| left insula | -0.011 | 0.012 | ** | 0.163 | 0.042 | n.s. |
| right bankssts | -0.003 | 0.004 | * | 0.046 | 0.052 | ** |
| right caudalanteriorcingulate | 0.025 | -0.028 | ** | 0.233 | -0.061 | n.s. |
| right caudalmiddlefrontal | 0.006 | -0.006 | ** | 0.090 | 0.011 | ** |
| right cuneus | 0.000 | 0.000 | n.s. | 0.001 | 0.045 | * |
| right entorhinal | 0.002 | -0.003 | n.s. | 0.013 | 0.010 | n.s. |
| right fusiform | -0.001 | 0.002 | n.s. | 0.027 | 0.005 | n.s. |
| right inferiorparietal | 0.005 | -0.005 | ** | 0.098 | 0.109 | ** |
| right inferiortemporal | -0.002 | 0.003 | n.s. | 0.035 | 0.002 | n.s. |
| right isthmuscingulate | 0.008 | -0.009 | ** | 0.088 | -0.045 | ** |
| right lateraloccipital | 0.002 | -0.001 | n.s. | 0.022 | 0.098 | ** |
| right lateralorbitofrontal | 0.001 | 0.000 | n.s. | 0.010 | 0.049 | ** |
| right lingual | -0.004 | 0.005 | ** | 0.070 | 0.061 | n.s. |
| right medialorbitofrontal | 0.001 | 0.000 | n.s. | 0.009 | 0.037 | n.s. |
| right middletemporal | -0.006 | 0.007 | ** | 0.099 | 0.060 | ** |
| right parahippocampal | 0.019 | -0.020 | ** | 0.153 | 0.032 | n.s. |
| right paracentral | 0.002 | -0.002 | n.s. | 0.028 | 0.051 | ** |
| right parsopercularis | -0.002 | 0.003 | n.s. | 0.031 | 0.000 | ** |
| right parsorbitalis | 0.015 | -0.016 | ** | 0.155 | 0.007 | n.s. |
| right parstriangularis | 0.002 | -0.002 | n.s. | 0.029 | -0.016 | ** |
| right pericalcarine | 0.000 | 0.000 | n.s. | 0.002 | 0.028 | n.s. |
| right postcentral | 0.007 | -0.007 | ** | 0.121 | -0.012 | ** |
| right posteriorcingulate | 0.005 | -0.005 | ** | 0.067 | -0.021 | * |
| right precentral | 0.005 | -0.005 | ** | 0.091 | 0.052 | ** |
| right precuneus | -0.003 | 0.004 | ** | 0.059 | 0.048 | ** |
| right rostralanteriorcingulate | 0.008 | -0.008 | ** | 0.067 | 0.033 | n.s. |
| right rostralmiddlefrontal | 0.003 | -0.003 | ** | 0.053 | 0.073 | ** |
| right superiorfrontal | 0.010 | -0.011 | ** | 0.169 | 0.045 | n.s. |
| right superiorparietal | 0.006 | -0.006 | ** | 0.119 | 0.102 | ** |
| right superiortemporal | -0.006 | 0.007 | ** | 0.094 | 0.077 | ** |
| right supramarginal | 0.004 | -0.004 | ** | 0.069 | 0.051 | ** |
| right frontalpole | 0.018 | -0.019 | ** | 0.131 | -0.009 | n.s. |
| right temporalpole | -0.010 | 0.011 | ** | 0.060 | 0.005 | n.s. |
| right transversetemporal | 0.008 | -0.0[illegible]8 | ** | 0.077 | 0.104 | * |
| right insula | -0.010 | 0.012 | ** | 0.142 | 0.069 | ** |

### Supplementary Table 3A

| Subcortical | Intercept | (s.e.) | P | Age2 | (s.e.) | P | Sex | (s.e.) | P | Sex by age2 | (s.e.) | P |
|---|---|---|---|---|---|---|---|---|---|---|---|---|
| left thal | 613.049 | 6.008 | ** | 8695.484 | 716.539 | ** | 74.940 | 8.682 | ** | -1187.243 | 1014.991 | n.s. |
| right thal | 531.871 | 5.364 | ** | 4171.752 | 639.774 | ** | 95.209 | 7.752 | ** | -1244.554 | 906.251 | n.s. |
| left caud | 364.783 | 3.521 | ** | 2190.319 | 419.998 | ** | 29.834 | 5.089 | ** | 834.754 | 594.934 | n.s. |
| right caud | 377.897 | 3.603 | ** | 3303.418 | 429.729 | ** | 28.284 | 5.207 | ** | 479.254 | 608.719 | n.s. |
| left put | 495.407 | 4.834 | ** | 6160.964 | 576.548 | ** | 57.544 | 6.986 | ** | -323.645 | 816.691 | n.s. |
| right put | 464.079 | 4.595 | ** | 6698.088 | 547.991 | ** | 50.995 | 6.640 | ** | -614.940 | 776.239 | n.s. |
| left pal | 170.576 | 1.763 | ** | 1939.804 | 210.284 | ** | 27.772 | 2.548 | ** | -317.629 | 297.871 | n.s. |
| right pal | 143.455 | 1.497 | ** | 1092.346 | 178.526 | ** | 23.127 | 2.163 | ** | 90.990 | 252.886 | n.s. |
| left hippo | 320.138 | 3.201 | ** | 3512.741 | 381.787 | ** | 31.862 | 4.626 | ** | -1374.551 | 540.807 | n.s. |
| right hippo | 311.867 | 3.121 | ** | 2672.383 | 372.203 | ** | 39.214 | 4.510 | ** | -605.863 | 527.232 | n.s. |
| left amyg | 151.438 | 1.523 | ** | 1726.265 | 181.644 | ** | 14.425 | 2.201 | ** | -576.511 | 257.302 | n.s. |
| right amyg | 156.565 | 1.596 | ** | 1835.945 | 190.409 | ** | 16.531 | 2.307 | ** | -660.039 | 269.717 | n.s. |
| left accumb | 85.209 | 0.858 | ** | 1188.991 | 102.329 | ** | 6.094 | 1.240 | ** | -58.930 | 144.951 | n.s. |
| right accumb | 80.431 | 0.786 | ** | 1059.372 | 93.692 | ** | 5.613 | 1.135 | ** | -244.057 | 132.716 | n.s. |

Supplementary Table 3B

| Surface area | Intercept | (s.e.) | P | Age2 | (s.e.) | P | Sex | (s.e.) | P | Sex by age2 | (s.e.) | P |
|---|---|---|---|---|---|---|---|---|---|---|---|---|
| left bankssts | 126.857 | 1.375 | ** | 468.035 | 148.508 | ** | 16.754 | 2.055 | ** | -144.283 | 217.605 | n.s. |
| left caudalanteriorcingulate | 126.787 | 1.375 | ** | 439.050 | 148.445 | ** | 16.823 | 2.054 | ** | -122.407 | 217.513 | n.s. |
| left caudalmiddlefrontal | 104.429 | 1.114 | ** | -125.449 | 120.351 | n.s. | 3.901 | 1.666 | * | -127.352 | 176.348 | n.s. |
| left cuneus | 293.437 | 2.943 | ** | 502.730 | 317.810 | n.s. | 22.109 | 4.398 | ** | 63.451 | 465.680 | n.s. |
| left entorhinal | 154.473 | 1.609 | ** | -258.284 | 173.761 | n.s. | 13.102 | 2.405 | ** | 285.420 | 254.608 | n.s. |
| left fusiform | 57.088 | 0.651 | ** | 246.897 | 70.291 | ** | 9.333 | 0.973 | ** | -60.093 | 102.996 | n.s. |
| left inferiorparietal | 303.903 | 3.104 | ** | 733.041 | 335.160 | * | 36.622 | 4.638 | ** | -711.684 | 491.103 | n.s. |
| left inferiortemporal | 453.623 | 4.702 | ** | 1832.218 | 507.743 | ** | 64.389 | 7.027 | ** | -1264.030 | 743.985 | n.s. |
| left isthmuscingulate | 351.679 | 3.537 | ** | 2118.970 | 381.998 | ** | 33.152 | 5.287 | ** | -1028.874 | 559.733 | n.s. |
| left lateraloccipital | 116.786 | 1.250 | ** | 1.485 | 135.027 | n.s. | 19.631 | 1.869 | ** | 208.057 | 197.853 | n.s. |
| left lateralorbitofrontal | 439.073 | 4.476 | ** | 408.166 | 483.345 | n.s. | 49.659 | 6.689 | ** | 303.046 | 708.234 | n.s. |
| left lingual | 207.272 | 2.114 | ** | 837.294 | 228.332 | ** | 21.655 | 3.160 | ** | -397.922 | 334.570 | n.s. |
| left medialorbitofrontal | 310.562 | 3.142 | ** | 62.805 | 339.266 | n.s. | 29.594 | 4.695 | ** | -899.943 | 497.120 | n.s. |
| left middletemporal | 172.650 | 1.796 | ** | -186.564 | 193.995 | n.s. | 23.477 | 2.685 | ** | -138.893 | 284.257 | n.s. |
| left parahippocampal | 296.470 | 2.997 | ** | 1349.148 | 323.622 | ** | 32.342 | 4.479 | ** | -635.081 | 474.196 | n.s. |
| left paracentral | 72.659 | 0.887 | ** | 146.999 | 95.780 | n.s. | 10.826 | 1.326 | ** | -99.995 | 140.344 | n.s. |
| left parsopercularis | 133.455 | 1.420 | ** | -209.260 | 153.336 | n.s. | 18.981 | 2.122 | ** | 191.968 | 224.681 | n.s. |
| left parsorbitalis | 192.892 | 2.113 | ** | 240.095 | 228.190 | n.s. | 32.407 | 3.158 | ** | -19.421 | 334.363 | n.s. |
| left parstriangularis | 61.740 | 0.641 | ** | 174.928 | 69.257 | * | 7.148 | 0.958 | ** | -111.706 | 101.481 | n.s. |
| left pericalcarine | 148.648 | 1.525 | ** | 286.425 | 164.734 | n.s. | 19.352 | 2.280 | ** | -18.396 | 241.381 | n.s. |
| left postcentral | 171.902 | 1.691 | ** | -361.951 | 182.645 | * | 13.425 | 2.528 | ** | -136.405 | 267.625 | n.s. |
| left posteriorcingulate | 340.826 | 3.576 | ** | 88.942 | 386.228 | n.s. | 46.216 | 5.345 | ** | -999.934 | 565.932 | n.s. |
| left precentral | 130.418 | 1.364 | ** | -272.664 | 147.314 | n.s. | 13.878 | 2.039 | ** | 25.470 | 215.856 | n.s. |
| left precuneus | 361.433 | 3.930 | ** | -417.038 | 424.423 | n.s. | 46.837 | 5.874 | ** | -250.550 | 621.898 | n.s. |
| left rostralanteriorcingulate | 329.071 | 3.385 | ** | 150.942 | 365.610 | n.s. | 44.972 | 5.060 | ** | 256.184 | 535.721 | n.s. |
| left rostralmiddlefrontal | 113.811 | 1.158 | ** | 3.816 | 125.011 | n.s. | 7.655 | 1.730 | ** | 15.972 | 183.176 | n.s. |
| left superiorfrontal | 541.177 | 5.556 | ** | 1927.368 | 600.057 | ** | 65.054 | 8.304 | ** | -680.175 | 879.250 | n.s. |
| left superiorparietal | 578.286 | 6.021 | ** | 1700.014 | 650.221 | ** | 74.321 | 8.998 | ** | -1311.611 | 952.755 | n.s. |
| left superiortemporal | 470.859 | 4.787 | ** | 931.025 | 516.942 | n.s. | 57.562 | 7.154 | ** | 331.895 | 757.464 | n.s. |
| left supramarginal | 307.760 | 3.214 | ** | 819.335 | 347.036 | * | 40.690 | 4.803 | ** | 123.558 | 508.504 | n.s. |
| left frontalpole | 391.464 | 4.081 | ** | 966.018 | 440.730 | * | 58.784 | 6.099 | ** | -331.182 | 645.792 | n.s. |
| left temporalpole | 25.400 | 0.265 | ** | 47.772 | 28.616 | n.s. | 3.224 | 0.396 | ** | -83.854 | 41.930 | n.s. |
| left transversetemporal | 45.368 | 0.479 | ** | 194.716 | 51.710 | ** | 5.206 | 0.716 | ** | -137.908 | 75.770 | n.s. |
| left insula | 56.947 | 0.594 | ** | -7.686 | 64.171 | n.s. | 6.788 | 0.888 | ** | -6.444 | 94.028 | n.s. |
| right bankssts | 164.356 | 1.842 | ** | -399.121 | 198.942 | * | 16.907 | 2.753 | ** | 164.981 | 291.506 | n.s. |
| right caudalanteriorcingulate | 107.054 | 1.139 | ** | 374.190 | 122.986 | ** | 13.963 | 1.702 | ** | -141.376 | 180.209 | n.s. |
| right caudalmiddlefrontal | 114.635 | 1.199 | ** | -319.007 | 129.509 | * | 14.718 | 1.792 | ** | 67.915 | 189.767 | n.s. |
| right cuneus | 288.838 | 2.933 | ** | 364.976 | 316.703 | n.s. | 30.420 | 4.383 | ** | -269.229 | 464.058 | n.s. |
| right entorhinal | 152.670 | 1.657 | ** | -4.937 | 178.965 | n.s. | 16.423 | 2.477 | ** | 430.922 | 262.233 | n.s. |
| right fusiform | 57.903 | 0.641 | ** | 149.574 | 69.216 | * | 10.349 | 0.958 | ** | 60.903 | 101.421 | n.s. |
| right inferiorparietal | 294.737 | 2.995 | ** | 1058.200 | 323.467 | ** | 32.896 | 4.476 | ** | 100.632 | 473.969 | n.s. |
| right inferiortemporal | 506.192 | 5.248 | ** | 1296.520 | 566.710 | * | 81.216 | 7.843 | ** | -1318.390 | 830.387 | n.s. |
| right isthmuscingulate | 326.198 | 3.322 | ** | 1526.453 | 358.727 | ** | 29.511 | 4.964 | ** | -78.652 | 525.635 | n.s. |
| right lateraloccipital | 105.916 | 1.158 | ** | -81.030 | 125.054 | n.s. | 16.020 | 1.731 | ** | -82.175 | 183.240 | n.s. |
| right lateralorbitofrontal | 437.115 | 4.538 | ** | 27.875 | 490.026 | n.s. | 57.833 | 6.782 | ** | -959.819 | 718.025 | n.s. |
| right lingual | 220.136 | 2.285 | ** | 585.537 | 246.770 | * | 24.988 | 3.415 | ** | 106.253 | 361.586 | n.s. |
| right medialorbitofrontal | 289.218 | 3.000 | ** | 70.565 | 323.929 | n.s. | 34.006 | 4.483 | ** | -705.531 | 474.646 | n.s. |
| right middletemporal | 154.044 | 1.565 | ** | 530.659 | 169.031 | ** | 15.954 | 2.339 | ** | -334.799 | 247.678 | n.s. |
| right parahippocampal | 308.652 | 3.167 | ** | 1826.197 | 342.056 | ** | 35.701 | 4.734 | ** | -843.985 | 501.208 | n.s. |
| right paracentral | 69.859 | 0.780 | ** | 147.856 | 84.229 | n.s. | 12.062 | 1.166 | ** | -6.834 | 123.418 | n.s. |
| right parsopercularis | 156.203 | 1.670 | ** | -499.921 | 180.338 | ** | 25.403 | 2.496 | ** | 541.736 | 264.245 | n.s. |
| right parsorbitalis | 174.711 | 1.870 | ** | 50.267 | 201.936 | n.s. | 25.558 | 2.795 | ** | 38.219 | 295.892 | n.s. |
| right parstriangularis | 77.382 | 0.794 | ** | 363.966 | 85.693 | ** | 7.429 | 1.186 | ** | -146.078 | 125.564 | n.s. |
| right pericalcarine | 184.815 | 1.887 | ** | 262.275 | 203.816 | n.s. | 21.655 | 2.821 | ** | 94.098 | 298.647 | n.s. |
| right postcentral | 184.562 | 1.820 | ** | 86.236 | 196.575 | n.s. | 13.005 | 2.720 | ** | -583.514 | 288.038 | n.s. |
| right posteriorcingulate | 331.128 | 3.494 | ** | -256.202 | 377.343 | n.s. | 43.218 | 5.222 | ** | -322.962 | 552.913 | n.s. |
| right precentral | 133.855 | 1.413 | ** | 118.978 | 152.606 | n.s. | 14.929 | 2.112 | ** | 61.943 | 223.610 | n.s. |
| right precuneus | 373.987 | 4.130 | ** | 254.919 | 446.051 | n.s. | 53.178 | 6.173 | ** | -1474.581 | 653.589 | n.s. |
| right rostralanteriorcingulate | 355.954 | 3.689 | ** | 491.740 | 398.397 | n.s. | 42.230 | 5.513 | ** | -70.807 | 583.762 | n.s. |
| right rostralmiddlefrontal | 97.209 | 1.006 | ** | -322.561 | 108.606 | ** | 10.462 | 1.503 | ** | 194.938 | 159.138 | n.s. |
| right superiorfrontal | 562.107 | 5.703 | ** | 1950.479 | 615.924 | ** | 62.001 | 8.524 | ** | -562.373 | 902.501 | n.s. |
| right superiorparietal | 586.732 | 6.069 | ** | 1396.682 | 655.391 | * | 73.254 | 9.070 | ** | 99.951 | 960.330 | n.s. |
| right superiortemporal | 453.249 | 4.721 | ** | 768.519 | 509.852 | n.s. | 49.896 | 7.056 | ** | -407.728 | 747.076 | n.s. |
| right supramarginal | 281.280 | 2.903 | ** | 608.371 | 313.474 | n.s. | 31.507 | 4.338 | ** | -694.847 | 459.327 | n.s. |
| right frontalpole | 375.950 | 3.841 | ** | 732.416 | 414.824 | n.s. | 52.182 | 5.741 | ** | -29.682 | 607.832 | n.s. |
| right temporalpole | 34.365 | 0.353 | ** | -16.406 | 38.079 | n.s. | 2.976 | 0.527 | ** | 48.350 | 55.796 | n.s. |
| right transversetemporal | 44.135 | 0.457 | ** | 150.427 | 49.340 | ** | 5.188 | 0.683 | ** | 10.036 | 72.297 | n.s. |
| right insula | 43.318 | 0.436 | ** | -38.027 | 47.047 | n.s. | 4.407 | 0.651 | ** | 134.689 | 68.938 | n.s. |

### Supplementary Table 3C

| Thickness | Intercept | (s.e.) | P | Age2 | (s.e.) | P | Sex | (s.e.) | P | Sex by age2 | (s.e.) | P |
|---|---|---|---|---|---|---|---|---|---|---|---|---|
| left bankssts | 0.084 | 0.001 | ** | 2.320 | 0.095 | ** | 0.005 | 0.001 | ** | 0.066 | 0.135 | n.s. |
| left caudalanteriorcingulate | 0.141 | 0.001 | ** | 1.885 | 0.156 | ** | 0.003 | 0.002 | n.s. | -0.247 | 0.222 | n.s. |
| left caudalmiddlefrontal | 0.206 | 0.002 | ** | 2.014 | 0.223 | ** | -0.005 | 0.003 | n.s. | 0.158 | 0.317 | n.s. |
| left cuneus | 0.124 | 0.001 | ** | 1.968 | 0.140 | ** | 0.005 | 0.002 | ** | 0.366 | 0.198 | n.s. |
| left entorhinal | 0.111 | 0.001 | ** | 1.243 | 0.125 | ** | 0.003 | 0.002 | n.s. | 0.300 | 0.177 | n.s. |
| left fusiform | 0.266 | 0.003 | ** | 1.105 | 0.298 | ** | 0.002 | 0.004 | n.s. | 0.744 | 0.424 | n.s. |
| left inferiorparietal | 0.116 | 0.001 | ** | 1.049 | 0.131 | ** | 0.002 | 0.002 | n.s. | 0.428 | 0.186 | n.s. |
| left inferiortemporal | 0.112 | 0.001 | ** | 2.475 | 0.126 | ** | 0.006 | 0.002 | ** | -0.005 | 0.178 | n.s. |
| left isthmuscingulate | 0.131 | 0.001 | ** | 1.354 | 0.145 | ** | 0.002 | 0.002 | n.s. | 0.112 | 0.205 | n.s. |
| left lateraloccipital | 0.168 | 0.002 | ** | 2.249 | 0.181 | ** | -0.002 | 0.002 | n.s. | -0.062 | 0.257 | n.s. |
| left lateralorbitofrontal | 0.099 | 0.001 | ** | 1.086 | 0.111 | ** | 0.005 | 0.001 | ** | 0.336 | 0.158 | n.s. |
| left lingual | 0.130 | 0.001 | ** | 2.192 | 0.148 | ** | 0.008 | 0.002 | ** | 0.370 | 0.211 | n.s. |
| left medialorbitofrontal | 0.102 | 0.001 | ** | 1.786 | 0.115 | ** | 0.002 | 0.001 | n.s. | -0.019 | 0.163 | n.s. |
| left middletemporal | 0.140 | 0.001 | ** | 2.002 | 0.158 | ** | 0.002 | 0.002 | n.s. | -0.220 | 0.224 | n.s. |
| left parahippocampal | 0.132 | 0.001 | ** | 1.873 | 0.148 | ** | 0.006 | 0.002 | ** | 0.414 | 0.210 | n.s. |
| left paracentral | 0.249 | 0.002 | ** | 0.884 | 0.263 | ** | 0.004 | 0.003 | n.s. | 0.709 | 0.373 | n.s. |
| left parsopercularis | 0.129 | 0.001 | ** | 2.123 | 0.143 | ** | 0.004 | 0.002 | * | 0.110 | 0.203 | n.s. |
| left parsorbitalis | 0.127 | 0.001 | ** | 2.030 | 0.139 | ** | 0.006 | 0.002 | ** | -0.150 | 0.198 | n.s. |
| left parstriangularis | 0.183 | 0.002 | ** | 2.713 | 0.200 | ** | 0.006 | 0.003 | * | -0.017 | 0.284 | n.s. |
| left pericalcarine | 0.138 | 0.001 | ** | 2.138 | 0.152 | ** | 0.007 | 0.002 | ** | 0.439 | 0.216 | n.s. |
| left postcentral | 0.104 | 0.001 | ** | 0.810 | 0.121 | ** | 0.001 | 0.002 | n.s. | 0.196 | 0.171 | n.s. |
| left posteriorcingulate | 0.099 | 0.001 | ** | 1.541 | 0.111 | ** | 0.005 | 0.001 | ** | 0.312 | 0.157 | n.s. |
| left precentral | 0.134 | 0.001 | ** | 2.195 | 0.147 | ** | 0.004 | 0.002 | * | -0.051 | 0.209 | n.s. |
| left precuneus | 0.113 | 0.001 | ** | 1.752 | 0.126 | ** | 0.004 | 0.002 | ** | -0.019 | 0.179 | n.s. |
| left rostralanteriorcingulate | 0.113 | 0.001 | ** | 2.472 | 0.125 | ** | 0.004 | 0.002 | ** | 0.354 | 0.178 | n.s. |
| left rostralmiddlefrontal | 0.195 | 0.002 | ** | 2.076 | 0.213 | ** | -0.004 | 0.003 | n.s. | 0.088 | 0.302 | n.s. |
| left superiorfrontal | 0.115 | 0.001 | ** | 2.092 | 0.131 | ** | 0.007 | 0.002 | ** | 0.617 | 0.186 | * |
| left superiorparietal | 0.127 | 0.001 | ** | 2.679 | 0.142 | ** | 0.003 | 0.002 | n.s. | 0.265 | 0.202 | n.s. |
| left superiortemporal | 0.101 | 0.001 | ** | 2.096 | 0.113 | ** | 0.005 | 0.001 | ** | -0.027 | 0.160 | n.s. |
| left supramarginal | 0.131 | 0.001 | ** | 2.277 | 0.143 | ** | 0.005 | 0.002 | ** | 0.090 | 0.203 | n.s. |
| left frontalpole | 0.119 | 0.001 | ** | 2.386 | 0.129 | ** | 0.005 | 0.002 | ** | -0.027 | 0.184 | n.s. |
| left temporalpole | 0.248 | 0.002 | ** | 2.941 | 0.275 | ** | 0.005 | 0.003 | n.s. | 0.335 | 0.391 | n.s. |
| left transversetemporal | 0.267 | 0.003 | ** | 0.298 | 0.306 | n.s. | 0.006 | 0.004 | n.s. | 1.223 | 0.434 | n.s. |
| left insula | 0.184 | 0.002 | ** | 2.050 | 0.200 | ** | 0.000 | 0.003 | n.s. | 0.296 | 0.284 | n.s. |
| right bankssts | 0.128 | 0.001 | ** | 1.806 | 0.142 | ** | 0.003 | 0.002 | n.s. | 0.212 | 0.201 | n.s. |
| right caudalanteriorcingulate | 0.149 | 0.001 | ** | 2.170 | 0.163 | ** | 0.004 | 0.002 | n.s. | -0.116 | 0.231 | n.s. |
| right caudalmiddlefrontal | 0.188 | 0.002 | ** | 2.080 | 0.206 | ** | -0.005 | 0.003 | * | -0.227 | 0.292 | n.s. |
| right cuneus | 0.124 | 0.001 | ** | 1.943 | 0.136 | ** | 0.003 | 0.002 | n.s. | -0.065 | 0.193 | n.s. |
| right entorhinal | 0.113 | 0.001 | ** | 1.416 | 0.124 | ** | 0.002 | 0.002 | n.s. | 0.045 | 0.177 | n.s. |
| right fusiform | 0.290 | 0.003 | ** | 1.469 | 0.318 | ** | 0.003 | 0.004 | n.s. | 0.460 | 0.452 | n.s. |
| right inferiorparietal | 0.116 | 0.001 | ** | 1.602 | 0.131 | ** | 0.002 | 0.002 | n.s. | 0.178 | 0.186 | n.s. |
| right inferiortemporal | 0.113 | 0.001 | ** | 2.722 | 0.127 | ** | 0.007 | 0.002 | ** | 0.056 | 0.180 | n.s. |
| right isthmuscingulate | 0.128 | 0.001 | ** | 1.380 | 0.144 | ** | 0.004 | 0.002 | * | 0.281 | 0.205 | n.s. |
| right lateraloccipital | 0.164 | 0.002 | ** | 2.094 | 0.177 | ** | -0.001 | 0.002 | n.s. | -0.040 | 0.252 | n.s. |
| right lateralorbitofrontal | 0.104 | 0.001 | ** | 1.331 | 0.116 | ** | 0.007 | 0.001 | ** | 0.599 | 0.164 | * |
| right lingual | 0.135 | 0.001 | ** | 2.002 | 0.153 | ** | 0.005 | 0.002 | ** | 0.034 | 0.218 | n.s. |
| right medialorbitofrontal | 0.105 | 0.001 | ** | 1.756 | 0.117 | ** | 0.001 | 0.001 | n.s. | 0.186 | 0.167 | n.s. |
| right middletemporal | 0.148 | 0.001 | ** | 1.745 | 0.167 | ** | 0.005 | 0.002 | * | 0.194 | 0.236 | n.s. |
| right parahippocampal | 0.127 | 0.001 | ** | 1.952 | 0.142 | ** | 0.006 | 0.002 | ** | 0.103 | 0.202 | n.s. |
| right paracentral | 0.208 | 0.002 | ** | 1.224 | 0.230 | ** | 0.005 | 0.003 | n.s. | -0.332 | 0.326 | n.s. |
| right parsopercularis | 0.126 | 0.001 | ** | 1.900 | 0.139 | ** | 0.004 | 0.002 | * | 0.355 | 0.197 | n.s. |
| right parsorbitalis | 0.134 | 0.001 | ** | 1.732 | 0.145 | ** | 0.002 | 0.002 | n.s. | -0.161 | 0.205 | n.s. |
| right parstriangularis | 0.179 | 0.002 | ** | 2.199 | 0.196 | ** | 0.004 | 0.002 | n.s. | 0.389 | 0.278 | n.s. |
| right pericalcarine | 0.135 | 0.001 | ** | 1.844 | 0.147 | ** | 0.001 | 0.002 | n.s. | -0.122 | 0.208 | n.s. |
| right postcentral | 0.105 | 0.001 | ** | 0.793 | 0.120 | ** | 0.002 | 0.002 | n.s. | 0.050 | 0.171 | n.s. |
| right posteriorcingulate | 0.105 | 0.001 | ** | 1.942 | 0.118 | ** | 0.001 | 0.001 | n.s. | 0.118 | 0.167 | n.s. |
| right precentral | 0.133 | 0.001 | ** | 2.117 | 0.146 | ** | 0.000 | 0.002 | n.s. | 0.109 | 0.207 | n.s. |
| right precuneus | 0.113 | 0.001 | ** | 1.694 | 0.127 | ** | 0.005 | 0.002 | ** | 0.133 | 0.181 | n.s. |
| right rostralanteriorcingulate | 0.113 | 0.001 | ** | 2.499 | 0.125 | ** | 0.005 | 0.002 | ** | 0.244 | 0.178 | n.s. |
| right rostralmiddlefrontal | 0.188 | 0.002 | ** | 1.992 | 0.212 | ** | 0.008 | 0.003 | ** | -0.059 | 0.301 | n.s. |
| right superiorfrontal | 0.113 | 0.001 | ** | 1.736 | 0.130 | ** | 0.005 | 0.002 | ** | 0.250 | 0.184 | n.s. |
| right superiorparietal | 0.125 | 0.001 | ** | 2.589 | 0.139 | ** | 0.004 | 0.002 | * | 0.117 | 0.197 | n.s. |
| right superiortemporal | 0.103 | 0.001 | ** | 1.986 | 0.115 | ** | 0.004 | 0.001 | ** | 0.361 | 0.164 | n.s. |
| right supramarginal | 0.128 | 0.001 | ** | 2.116 | 0.141 | ** | 0.007 | 0.002 | ** | 0.191 | 0.201 | n.s. |
| right frontalpole | 0.121 | 0.001 | ** | 2.782 | 0.133 | ** | 0.005 | 0.002 | ** | 0.020 | 0.189 | n.s. |
| right temporalpole | 0.243 | 0.002 | ** | 2.417 | 0.269 | ** | 0.003 | 0.003 | n.s. | 0.592 | 0.382 | n.s. |
| right transversetemporal | 0.274 | 0.003 | ** | 0.825 | 0.323 | * | 0.003 | 0.004 | n.s. | 1.082 | 0.459 | n.s. |
| right insula | 0.183 | 0.002 | ** | 1.833 | 0.203 | ** | 0.010 | 0.003 | ** | 0.211 | 0.288 | n.s. |
